# *In silico* Proteome analysis of Severe acute respiratory syndrome coronavirus 2 (SARS-CoV-2)

**DOI:** 10.1101/2020.05.23.104919

**Authors:** Chittaranjan Baruah, Papari Devi, Dhirendra K. Sharma

## Abstract

Severe acute respiratory syndrome coronavirus 2 (SARS-CoV-2) (2019-nCoV), is a positive-sense, single-stranded RNA coronavirus. The virus is the causative agent of coronavirus disease 2019 (COVID-19) and is contagious through human-to-human transmission. The present study reports sequence analysis, complete coordinate tertiary structure prediction and *in silico* sequence-based and structure-based functional characterization of full SARS-CoV-2 proteome based on the NCBI reference sequence NC_045512 (29903 bp ss-RNA) which is identical to GenBank entry MN908947 and MT415321. The proteome includes 12 major proteins namely orf1ab polyprotein (includes 15 proteins), surface glycoprotein, ORF3a protein, envelope protein, membrane glycoprotein, ORF6 protein, ORF7a protein, orf7b, ORF8, Nucleocapsid phosphoprotein and ORF10 protein. Each protein of orf1ab polyprotein group has been studied separately. A total of 25 polypeptides have been analyzed out of which 15 proteins are not yet having experimental structures and only 10 are having experimental structures with known PDB IDs. Out of 15 newly predicted structures six (6) were predicted using comparative modeling and nine (09) proteins having no significant similarity with so far available PDB structures were modeled using *ab-initio* modeling. Structure verification using recent tools QMEANDisCo 4.0.0 and ProQ3 for global and local (per-residue) quality estimates indicate that the all-atom model of tertiary structure of high quality and may be useful for structure-based drug designing targets. The study has identified nine major targets (spike protein, envelop protein, membrane protein, nucleocapsid protein, 2’-O-ribose methyltransferase, endoRNAse, 3’-to-5’ exonuclease, RNA-dependent RNA polymerase and helicase) for which drug design targets could be considered. There are other 16 nonstructural proteins (NSPs), which may also be percieved from the drug design angle. The protein structures have been deposited to ModelArchive. Tunnel analysis revealed the presence of large number of tunnels in NSP3, ORF 6 protein and membrane glycoprotein indicating a large number of transport pathways for small ligands influencing their reactivity.

## INTRODUCTION

Severe acute respiratory syndrome coronavirus 2 (SARS-CoV-2), is a positive-sense, single-stranded RNA with genome size 26.2 and 31.7 kb coronavirus, covered by an enveloped structure (Yang *et al*., 2006), which is a major source of disaster in the 21^th^ century. The virus is the causative agent of coronavirus disease 2019 (COVID-19) and is contagious through human-to-human transmission. A typical CoV contains at least six ORFs in its genome. SARS-CoV-2 is the seventh coronavirus that is known to cause human disease. Coronaviruses (CoVs) are a group of large and enveloped viruses with positive-sense, single-stranded RNA genomes (Lai *et al*., 2007; Lu and Liu, 2012). Previously identified human CoVs that cause human disease include the alphaCoVs hCoV-NL63 and hCoV-229E and the betaCoVs HCoV-OC43, HKU1, severe acute respiratory syndrome CoV (SARS-CoV), and Middle East respiratory syndrome CoV (MERS-CoV) (Lu *et al*., 2015). Among these seven strains, three strains proved to be highly pathogenic (SARS-CoV, MERS-CoV, and 2019-nCoV), which caused endemic of severe CoV disease (Paules *et al*., 2020).The viruses can be classified into four genera: alpha, beta, gamma, and deltaCoVs (Woo *et al*., 2009). Both alphaCoVs and the betaCoVs HCoV-OC43 and HKU1 cause self-limiting common cold-like illnesses (Chiu *et al*., 2005; Jean *et al*., 2013). However, SARS-CoV and MERS-CoV infection can result in life-threatening disease and have pandemic potential. SARS-CoV-2 is responsible for the infection with special reference to the involvement of the respiratory tract (both lower and upper respiratory tract), e.g., common cold, pneumonia, bronchiolitis, rhinitis, pharyngitis, sinusitis, and other system symptoms such as occasional watery and diarrhea (Chang *et al*., 2016; Paules *et al*., 2020).

The current classification of coronaviruses recognizes 39 species in 27 subgenera, five genera and two subfamilies that belong to the family *Coronaviridae*, suborder *Cornidovirineae*, order *Nidovirales* and realm *Riboviria* (Ziebuhr *et al*., 2017; Siddell *et al*., 2019; Ziebuhr et *al*., 2019). The family classification and taxonomy are developed by the *Coronaviridae* Study Group (CSG), a working group of the ICTV (de Groot *et al*., 2012). To accommodate the wide spectrum of clinical presentations and outcomes of infections caused by SARS-CoV-2 (ranging from asymptomatic to severe or even fatal in some cases) (Huang *et al.*, 2020), the WHO recently introduced a rather unspecific name (coronavirus disease 19, also known as COVID-19 (World Health Organization, 2020) to denote this disease. Whole-genome sequencing results showed that the causative agent was a novel coronavirus that was initially named 2019-nCoV by the World Health Organization (WHO) (Wu *et al*., 2020; Zhou *et al*., 2020; Zhu *et al*., 2020). Later, the International Committee on Taxonomy of Viruses (ICTV) officially designated the virus SARS-CoV-2 (Coronaviridae Study Group of the International Committee on Taxonomy of Viruses, 2020), although many virologists argue that HCoV-19 is more appropriate (Jiang *et al*., 2020).

Understanding the complete proteome of SARS-CoV-2 is the need of the hour for the final destination of drug/medicine. The major structural proteins namely the E and M proteins, which form the viral envelope; the N protein, which binds to the virus’s RNA genome; and the S protein, which binds to human receptors, may be significant from the perspectives of drug design and development.. The nonstructural proteins get expressed as two long polypeptides, which gets chopped up by the virus’s main protease. This group of proteins includes the main protease (Nsp5) and RNA polymerase (Nsp12) has equal importance for structure based drug designing. In this regard, the present study reports sequence analysis and structure prediction (both Comparative and *ab-initio* modeling) of full SARS-CoV-2 proteome based on the NCBI reference sequence NC_045512 (29903 bp ss-RNA) which is identical to GenBank entry MN908947 and MT415321 (Table 1).

**Table 1.**
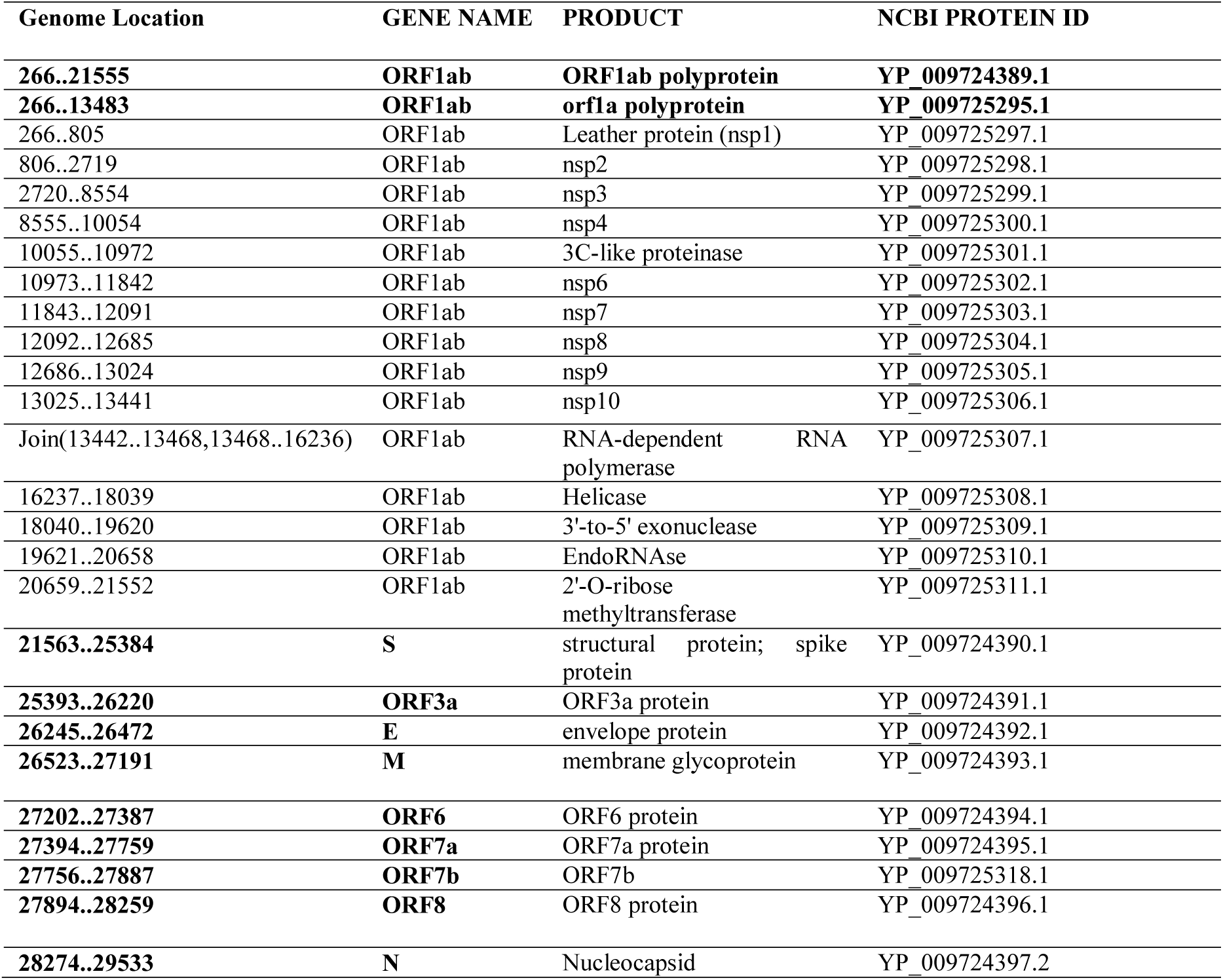
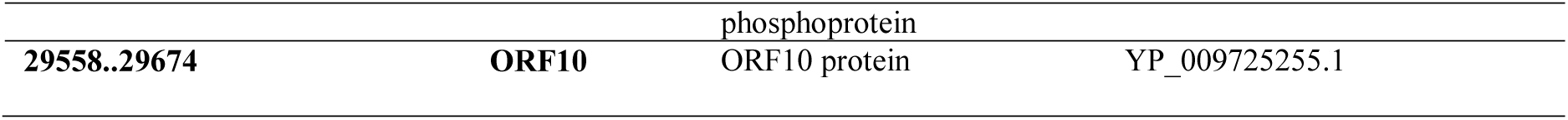
Complete Proteome of Severe acute respiratory syndrome coronavirus 2 (SARS-CoV-2) (Genome: NCBI LOCUS NC_045512; 29903 bp ss-RNA)

The viral genome encodes 29 proteins (*Nature* DOI: 10.1038/s41433-020-0790-7). In the present analysis 25 proteins of NCBI reference sequence NC_045512 (Figure 1) (including 15 proteins of orf1ab) from the proteome have been analyzed out of which 15 proteins are not yet having experimental structures and 10 are having experimental structures with known PDB IDs. Out of 15 predicted structures six (6) proteins namely NSP6, Surface glycoprotein, Envelope protein, ORF7a, ORF8 and Nucleoproteins were predicted using comparative modeling and nine (09) proteins namely NSP1, NSP2, NSP3, NSP4, ORF3a, Membrane protein, ORF6, ORF7b and ORF10 having no significant similarity with so far available PDB structures were modeled using *ab-initio* modeling.

**Figure 1.**
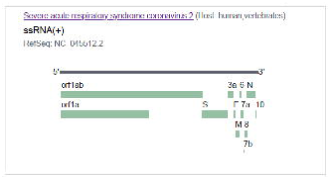
The structure of SARS-CoV-2 reference genome (<u>NC_045512.2</u>) used in the present study. The genome size is 29.9 Kb with GC% 38.0 and 11 genes which encodes 12 major proteins. SARS-CoV-2 has four structural proteins: the E and M proteins, which form the viral envelope; the N protein, which binds to the virus’s RNA genome; and the S protein, which binds to human receptors. (Source: NCBI)

## MATERIALS AND METHODS

### Acquisition and analysis of sequences

The sequence of full SARS-CoV-2 proteome based on the NCBI reference **NC_045512 (29903 bp ss-RNA)** along with GenBank entry MN908947 and **MT415321** were analyzed (**Table 1**). A total of 25 proteins from the proteome have been analyzed out of which 15 proteins are not yet having experimental structures based on BLASTp (Altschul *et al.*, 1997) and FASTA (Pearson, 1991) searches.

### Three-dimensional structure prediction (Comparative and ab-initio modeling)

3D structure predictions were carried out for fifteen (15) proteins which are without 3D co-ordinate files. BlastP and FASTA searches were performed independently to know the existing structure from the PDP, for suitable template for Comparative modeling and to select the proteins for which *ab-initio* modeling (**Table 2**). The significance of the BLAST results was assessed by expect values (e-value) generated by BLAST family of search algorithm and query coverage. Six (6) proteins namely NSP6, Surface glycoprotein, Envelope protein, ORF7a, ORF8 and Nucleoproteins had been predicted following Modeller9.23 program (Webb and Sali, 2016) and Baker Rosetta Server (https://robetta.bakerlab.org/); nine (09) proteins namely NSP1, NSP2, NSP3, NSP4, ORF3a, Membrane protein, ORF6, ORF7b and ORF10 having no similarity with the available PDB structures, were modeled for *ab-initio* following Baker Rosetta Server (https://robetta.bakerlab.org/). The loop regions were modeled using ModLoop server (Fiser and Sali, 2003). The final 3D structures with complete coordinates had been obtained by optimization of a molecular probability density function (pdf) of Modeller (Eswar *et al*., 2006). The molecular pdf for modelling was optimized with the variable target function procedure in Cartesian space that employed the method of conjugate gradients and molecular dynamics with simulated annealing (Sali and Blundell, 1993). The computational protein structures were verified by using Global and local (per-residue) quality estimates using ProQ3, ModFOLD6-TS, QMEANDisCo 4.0.0 (Haas *et al*., 2019). After verification, the coordinate files were successfully deposited to ModelArchive (https://www.modelarchive.org). All the graphic presentations of the 3D structures were prepared using Chimera version 1.8.1 (Pettersen *et al.*, 2004) and pyMOL 0.97rc (*DeLano, 2002*).

**Table 2.**
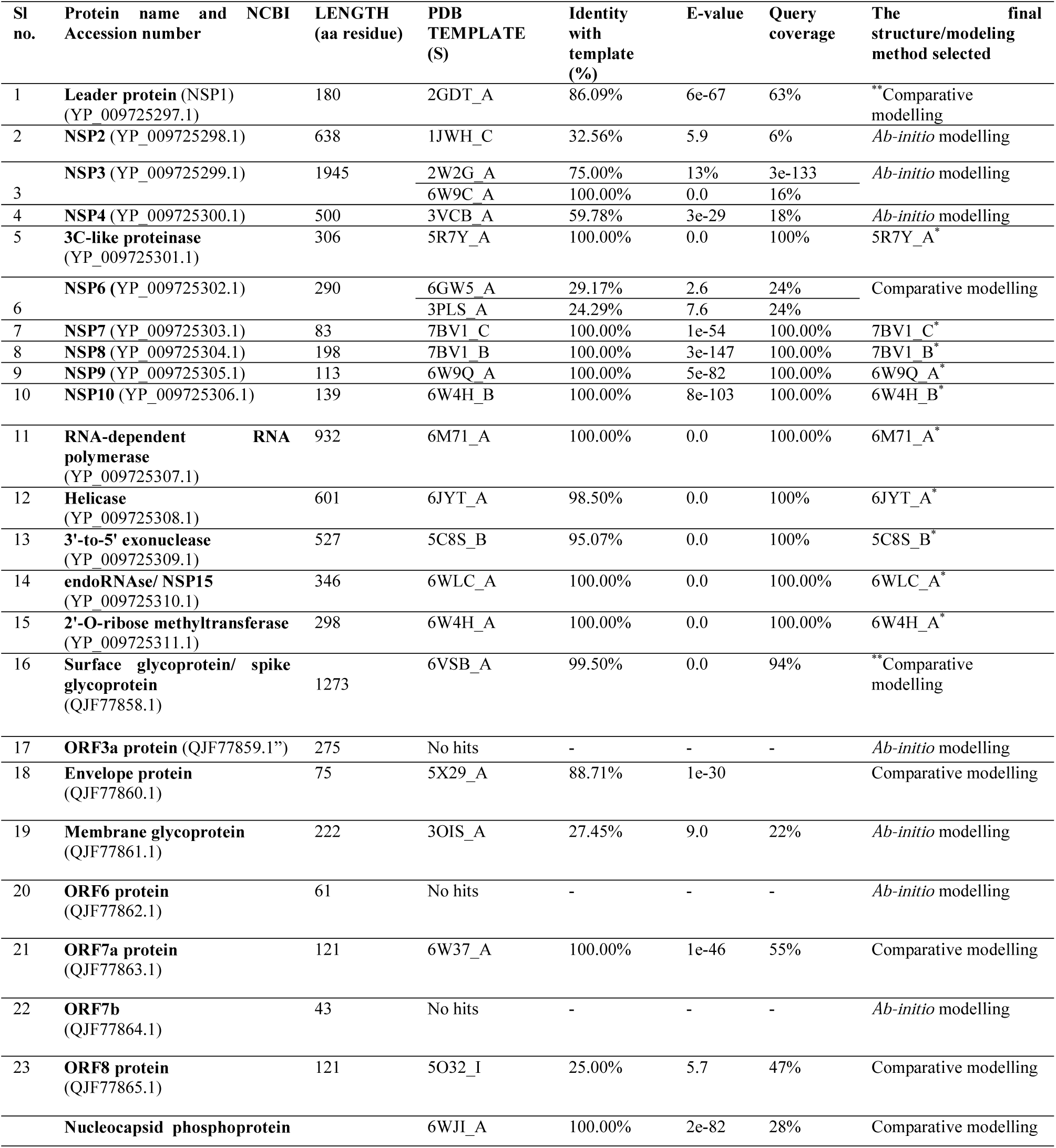
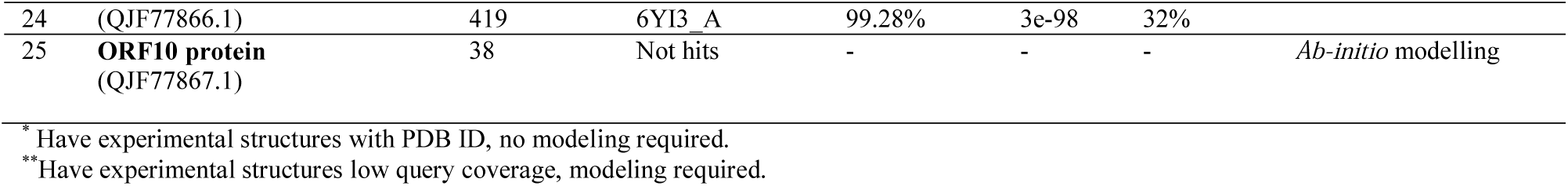
BLAST results against available PDB structures for selection of modeling method, template selection and existing PDB structures considered.

### Proteomics analysis

The proteomics analyses were carried out after ExPASy proteomic tools (https://www.expasy.org/tools). The Data mining, sequence analyses for physico-chemical parameters of SARS-CoV-2 proteomes were computed using ProtParam (Gasteiger *et al.*, 2005) and BioEdit. The important calculations for the amino acid composition, atomic composition, theoretical pI, molecular weight, Formula, extinction coefficients, half-life, instability index, aliphatic index, hydrophobicity, charge *vs.* pH were carried out under sequence analysis (**Table 3**).

**Table 3.**
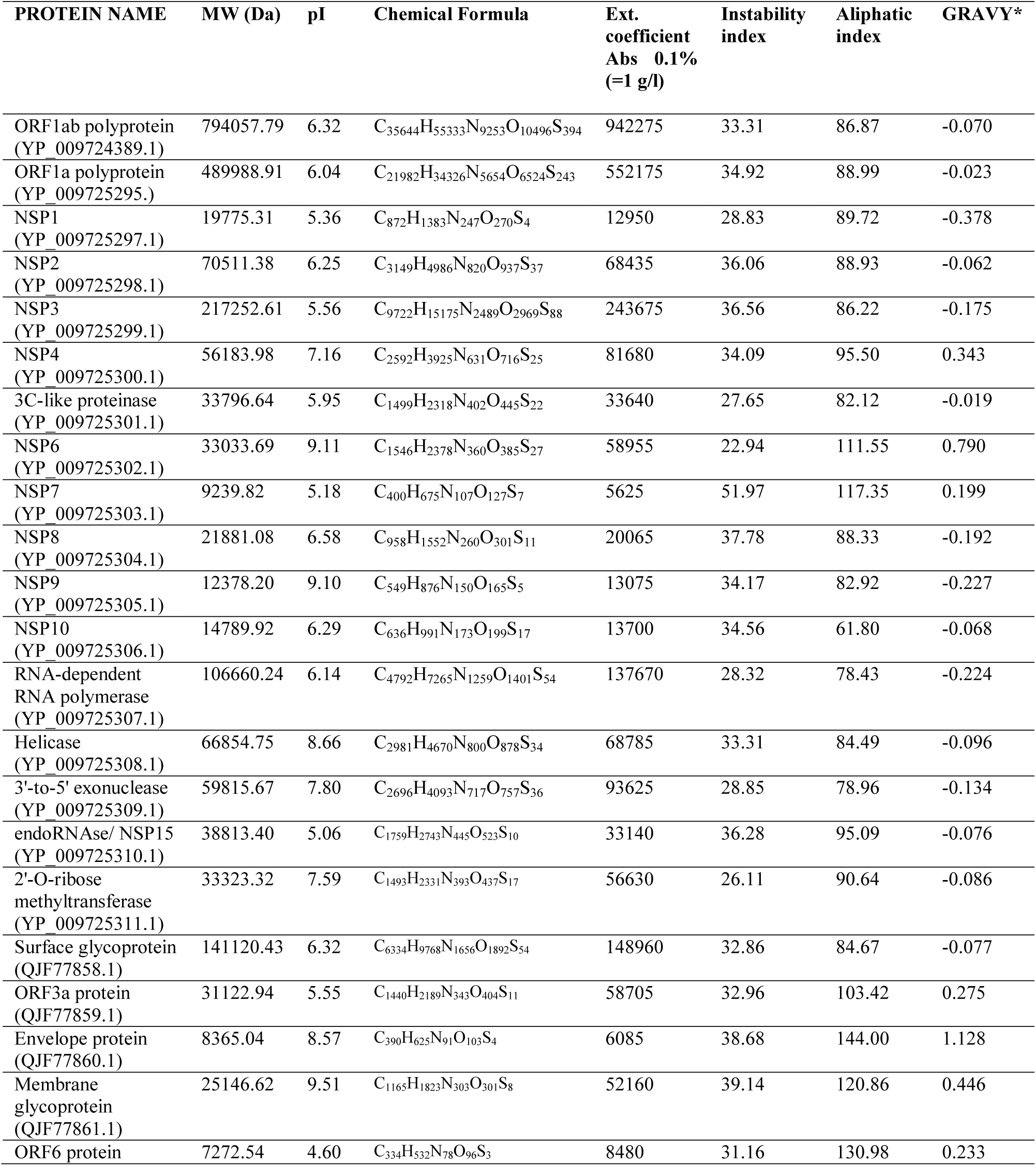
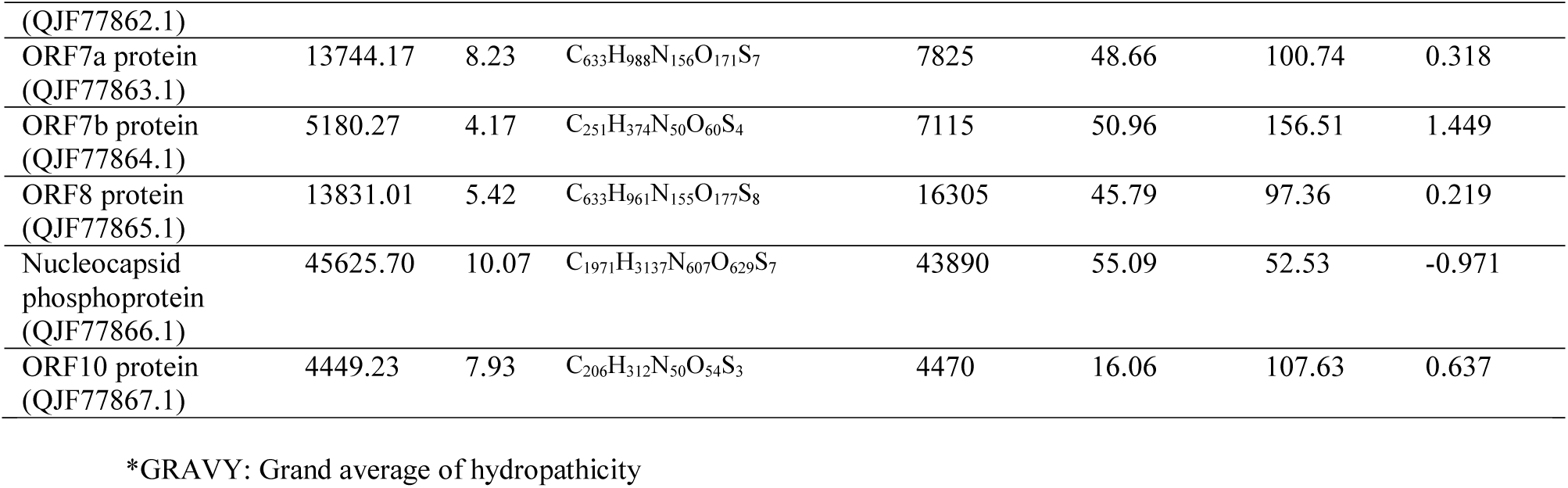
Physicochemical parameters of SARS-CoV-2 proteome

The sequence based functional annotation had been carried out for SARS-CoV-2 proteome in the sequence database, using Pfam (*pfam.sanger.ac.uk/-*), GO (www.geneontology.org/), KEGG database (www.genome.jp/kegg/). ProFunc server (Laskowski *et al*., 2005) was used to identify the likely biochemical function of proteins from the predicted 3D structure. Hmmer version 3.3 (Finn *et al*., 2011), PFam, PROSITE, PRINTS, ProDom, InterProScan were deployed for functional characterization. **MOLE 2.0** (Sehnal *et al*., 2013) and **Caver Web 1.0 (**Stourac *et al*., 2019) were used for advanced analysis of biomacromolecular channels. The theoretical structures of the present study are used for tunnels analysis using Caver Web 1.0. The tunnel bottleneck radius and lengthas have been calculated in Angstrom (Å) and throughput (estimated tunnel importance) calculated as *e*^*-cost*^, where *e* is Euler’s number.

## RESULTS AND DISCUSSION

The ORF1ab polyprotein is the largest protein (7096 amino acids; 794.017kDa) of the SARS-CoV-2, rich in Leucine (9.41%) and Valine (8.43%).The protein has pI of 6.32, Instability index 33.31, Aliphatic index 86.87 and Grand average of hydropathicity - 0.070 (Figure 2a; Table 3). It is a multifunctional protein involved in the transcription and replication of viral RNAs. It consists of proteinases responsible for the cleavages of the polyprotein.

**Figure 2(A).**
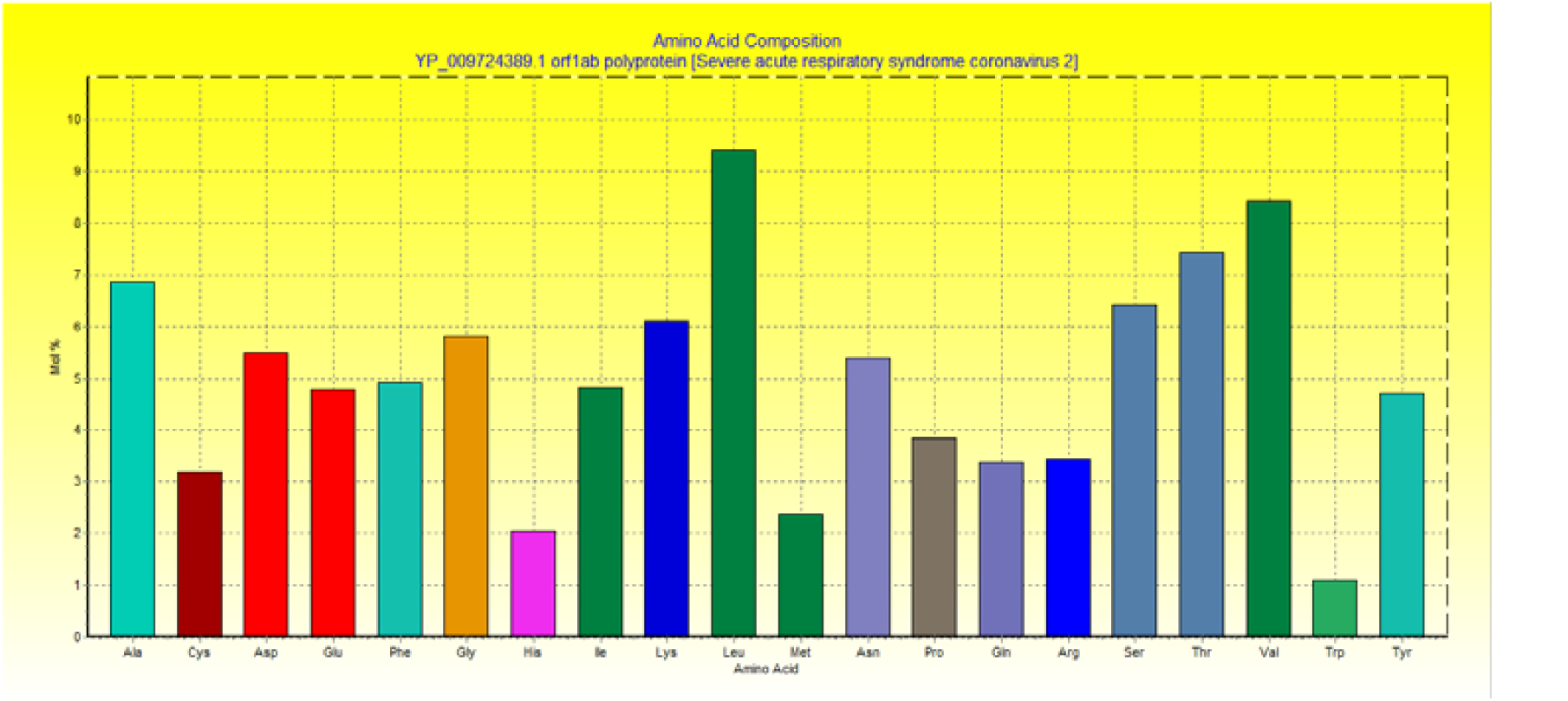
Amino acid composition of ORF 1ab protein. Amino acid distribution histogram for 12 mjor proteins of SARS-CoV-2 proteome

InterPro classification of protein families ORF1ab polyprotein belongs to Non-structural protein NSP1, betacoronavirus (IPR021590) protein family. The biologival process associated with ORF1ab protein includes-viral genome replication (*GO* :0019079), viral protein processing (*GO* :0019082), proteolysis (*GO* :0006508), transcription, DNA-templated (*GO* :0006351) and viral RNA genome replication (*GO* :0039694).

The molecular functions associated are zinc ion binding (*GO* :0008270), RNA binding (*GO* :0003723), omega peptidase activity (*GO* :0008242), cysteine-type endopeptidase activity (*GO* :0004197), transferase activity (*GO* :0016740), single-stranded RNA binding (*GO* :0003727), cysteine-type peptidase activity (*GO* :0008234), RNA-directed 5’-3’ RNA polymerase activity (*GO* :0003968), ATP binding (*GO* :0005524) and nucleic acid binding (*GO* :0003676).

Hits for all PROSITE (release 2020_02) motifs on sequence of ORF1ab (YP_009724389-1) has detected 8 hits: (i) Peptidase family C16 domain profile : PS51124| PEPTIDASE_C16 (profile) Peptidase family C16 domain profile :1634 - 1898: score=60.973; (ii) Corona virus main protease (M-pro) domain profile : PS51442|M_PRO (profile) Corona virus main protease (M-pro) domain profile : 3264 - 3569: score=154.193; (iii) RdRp of positive ssRNA viruses catalytic domain profile : PS50507|RDRP_SSRNA_POS (profile) RdRp of positive ssRNA viruses catalytic domain profile : 5004 - 5166: score=8.290; (iv) Coronaviridae zinc-binding (CV ZBD) domain profile : PS51653| CV_ZBD (profile) Coronaviridae zinc-binding (CV ZBD) domain profile : 5325 - 5408: score=33.035; (v) (+)RNA virus helicase core domain profile : PS51657|PSRV_HELICASE (profile) (+)RNA virus helicase core domain profile : 5581 - 5932: score=27.299; (vi) Carbamoyl-phosphate synthase subdomain signature 2 : PS00867|CPSASE_2 (pattern) Carbamoyl-phosphate synthase subdomain signature 2 : 2061 - 2068: [confidence level: (−1)]; vii) Lipocalin signature : PS00213| LIPOCALIN (pattern) Lipocalin signature : 4982 - 4993: [confidence level: (−1)].

The ORF1a polyprotein (Length = 4405 amino acids; Molecular Weight = 489.963kDa) is also rich in Leucine (9.88%) and Valine (8.42%). ORF1a polyprotein has theoretical Isoelectric point (pI) 6.04, Instability index 34.92, Aliphatic index 88.87 and Grand average of hydropathicity - 0.023 (Figure 2B; Table 3).

**Figure 2 (B).**
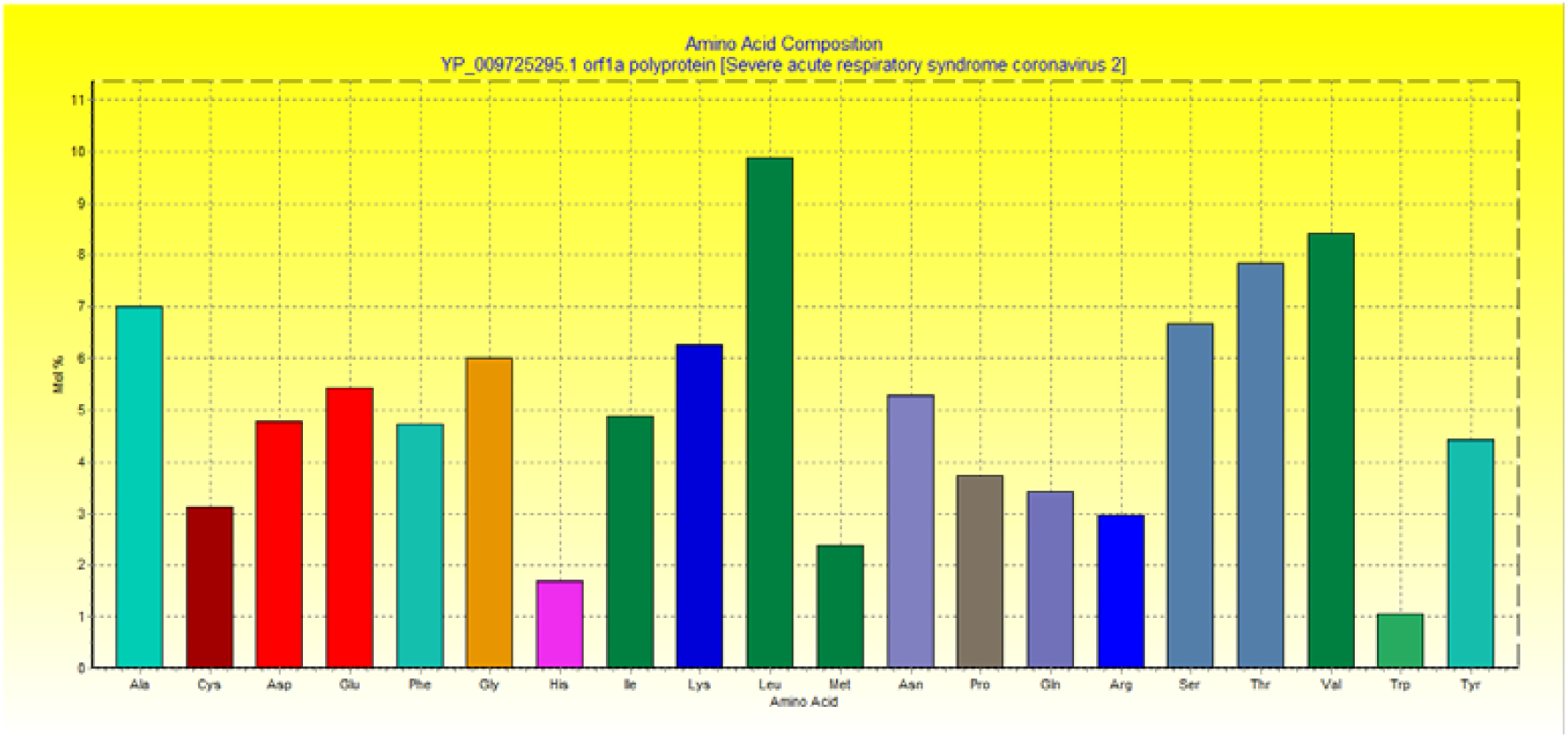
Amino acid composition of ORF 1a Protein. Amino acid distribution histogram for 12 mjor proteins of SARS-CoV-2 proteome

**Figure 2 (C).**
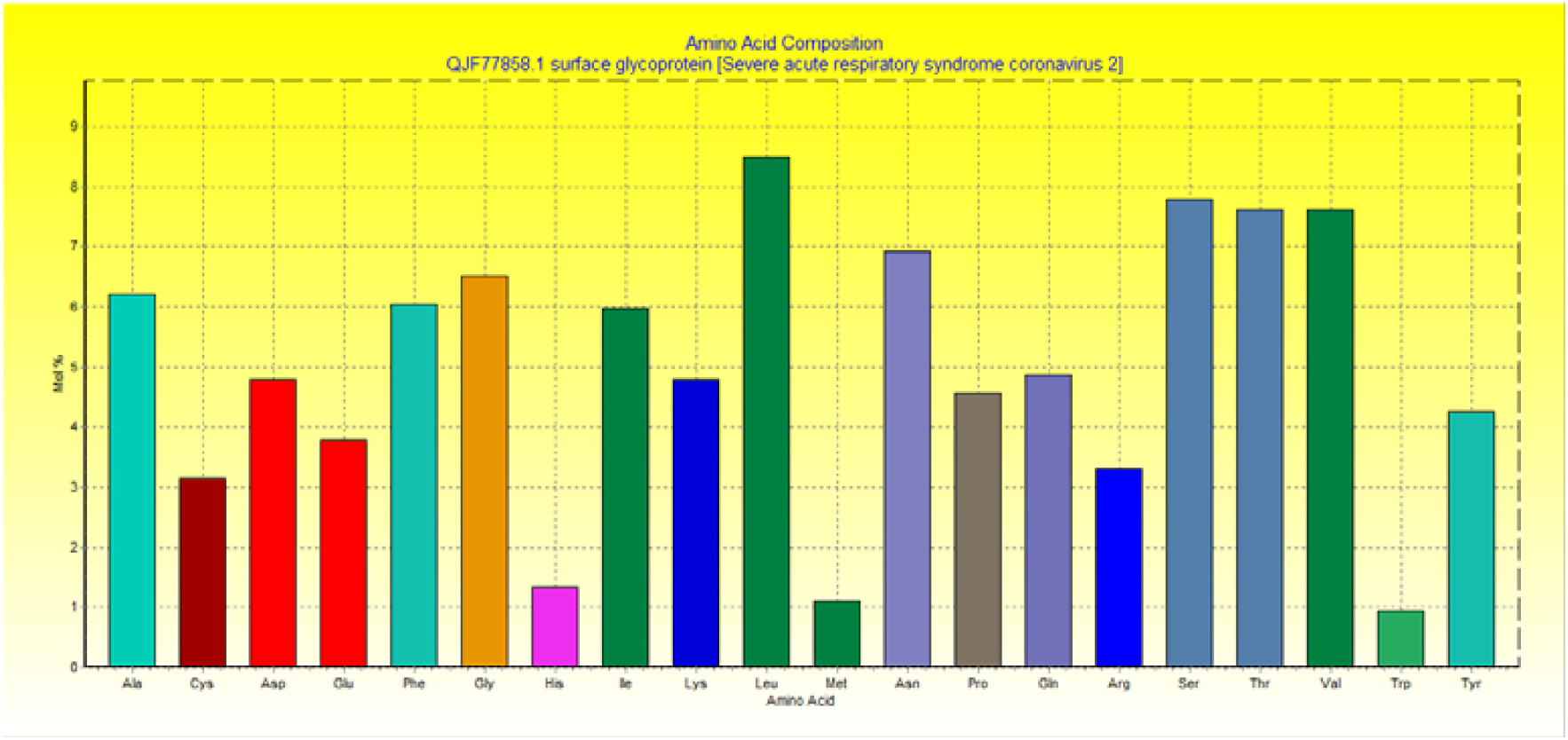
Amino acid composition of Surface glycoprotein. Amino acid distribution histogram for 12 mjor proteins of SARS-CoV-2 proteome

**Figure 2 (D).**
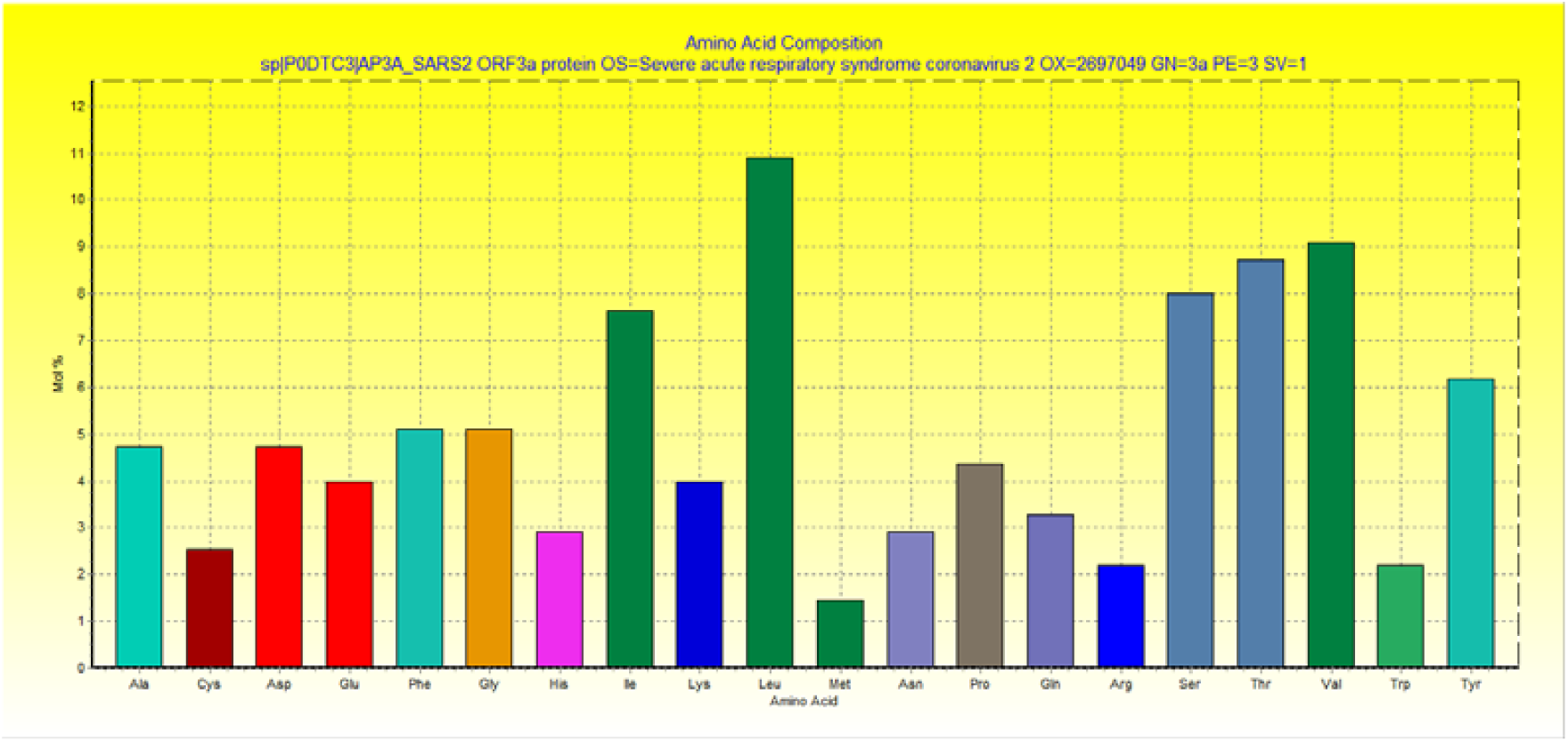
Amino acid composition of ORF 3a protein. Amino acid distribution histogram for 12 mjor proteins of SARS-CoV-2 proteome

**Figure 2 (E).**
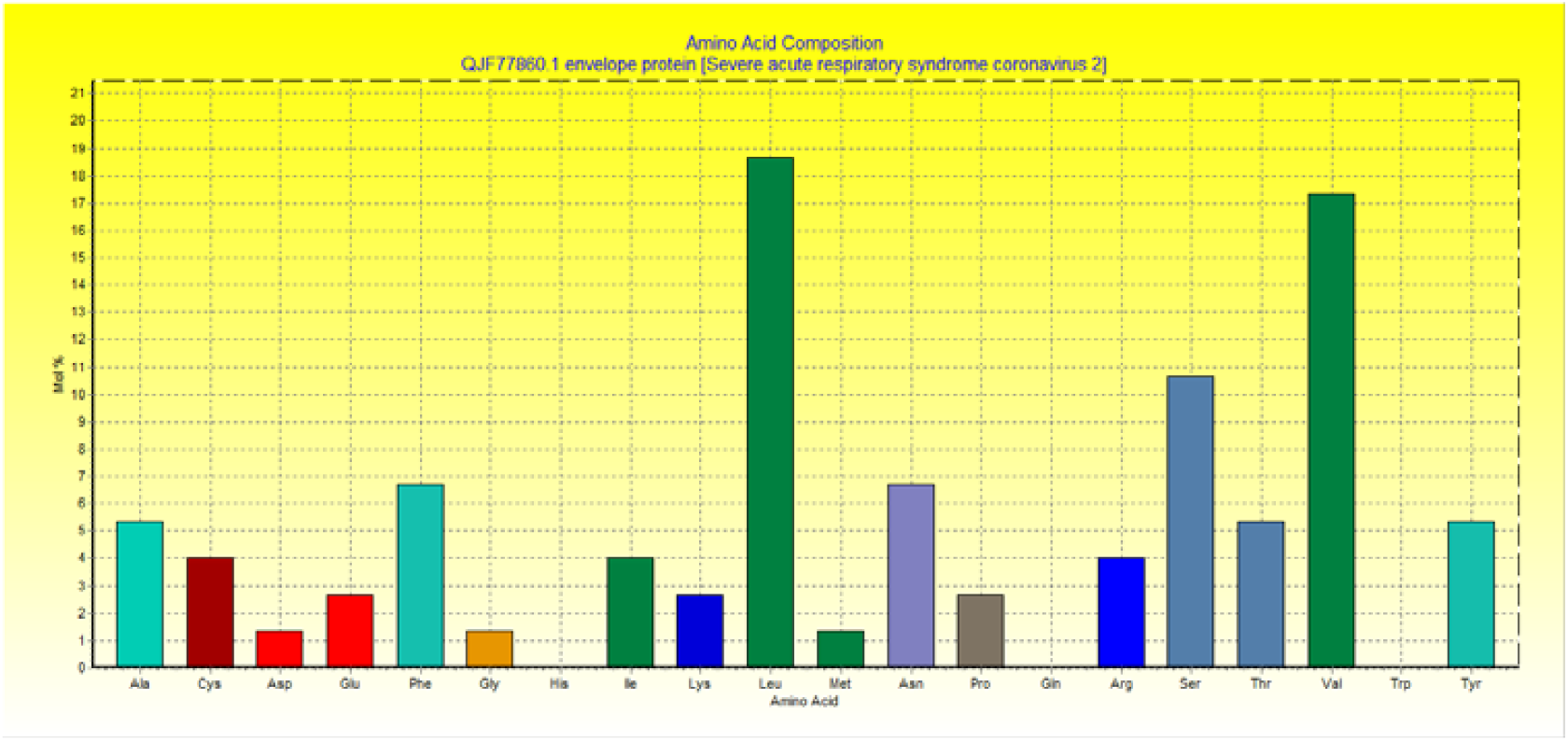
Amino acid composition of Envelope protein. Amino acid distribution histogram for 12 mjor proteins of SARS-CoV-2 proteome

**Figure 2 (F).**
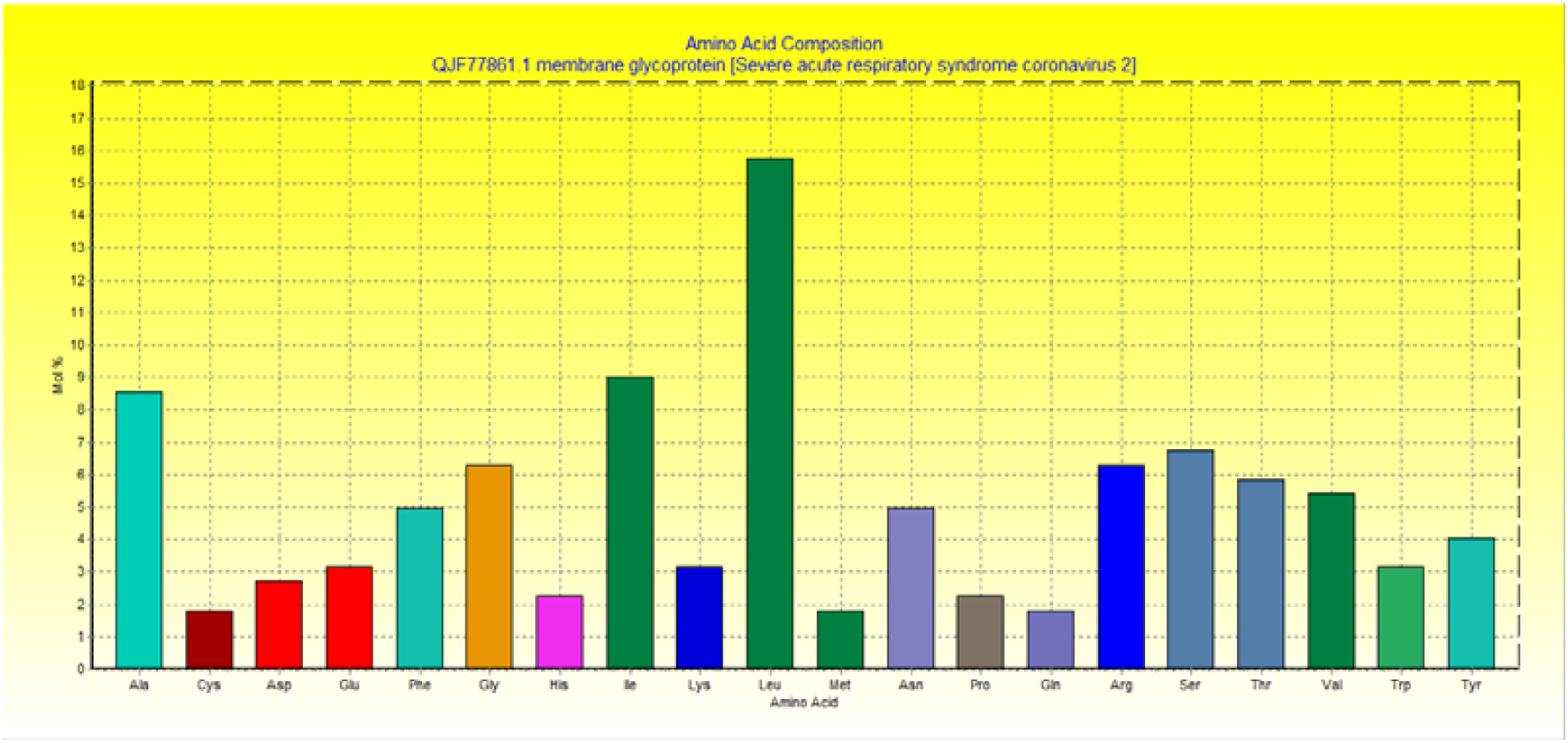
Amino acid composition of Membrane glycoprotein. Amino acid distribution histogram for 12 mjor proteins of SARS-CoV-2 proteome

**Figure 2 (G).**
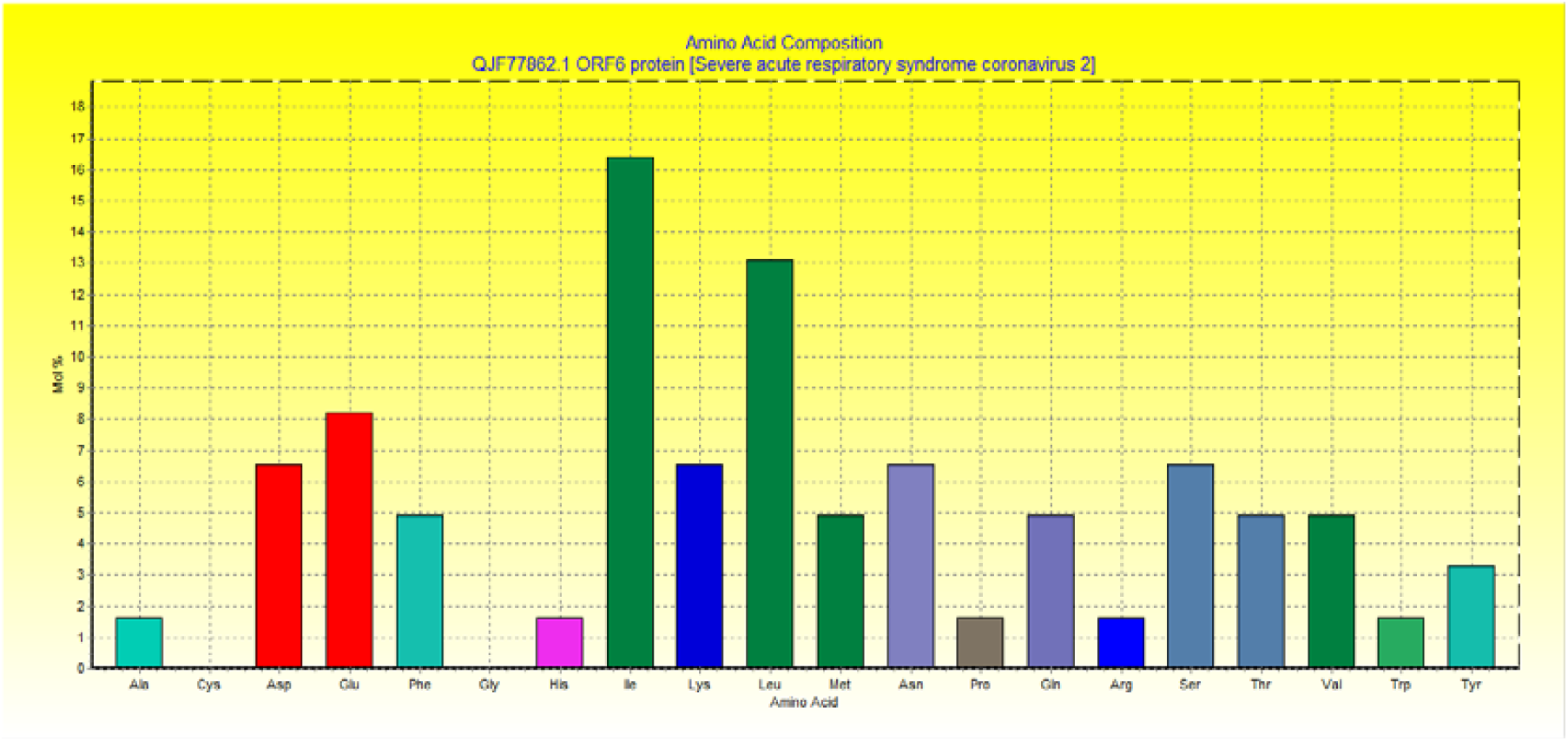
Amino acid composition of ORF6 protein. Amino acid distribution histogram for 12 mjor proteins of SARS-CoV-2 proteome

**Figure 2 (H).**
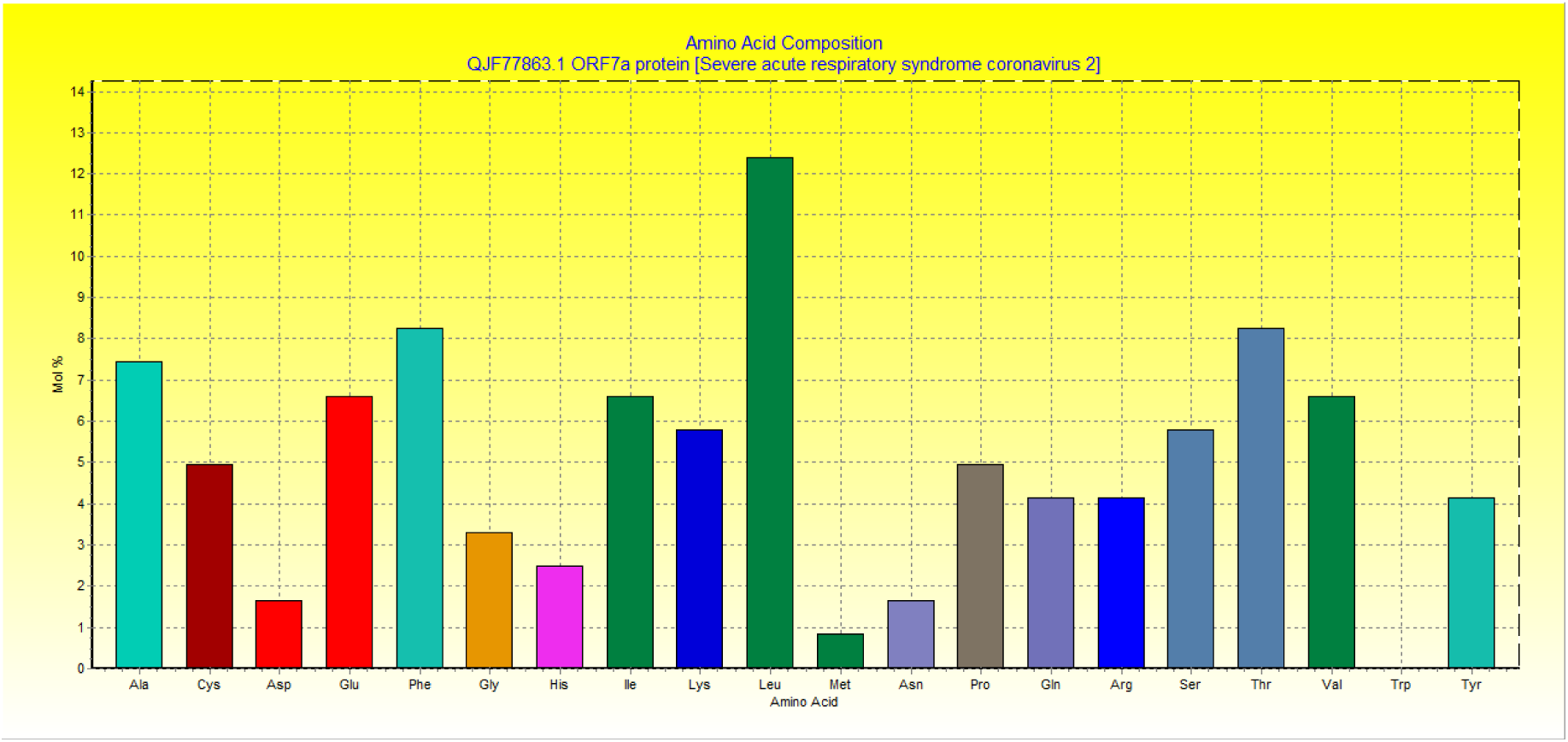
Amino acid composition of ORF 7a protein. Amino acid distribution histogram for 12 mjor proteins of SARS-CoV-2 proteome

**Figure 2 (I).**
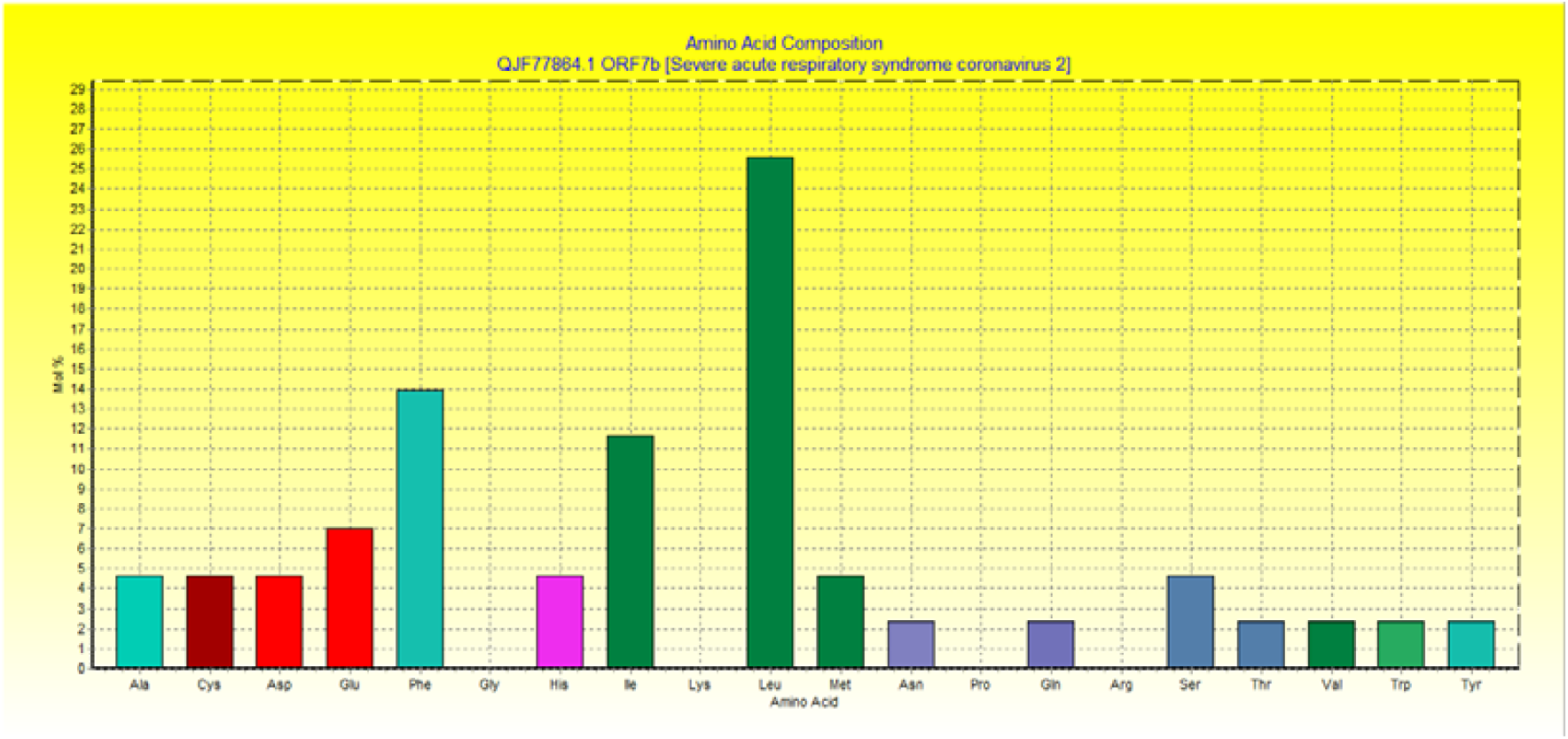
Amino acid composition of ORF7b protein. Amino acid distribution histogram for 12 mjor proteins of SARS-CoV-2 proteome

**Figure 2 (J).**
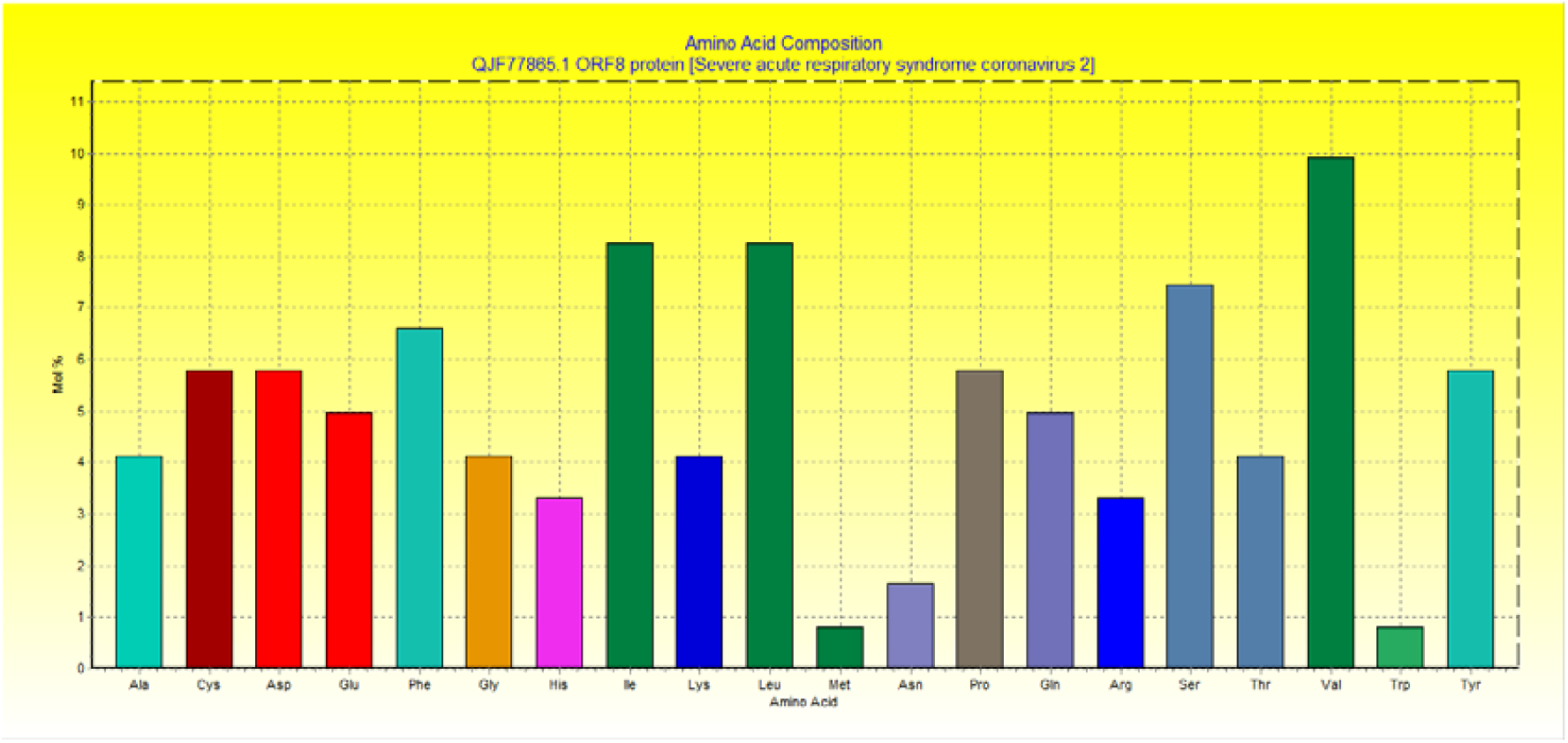
Amino acid composition of ORF8 protein. Amino acid distribution histogram for 12 mjor proteins of SARS-CoV-2 proteome

**Figure 2 (K).**
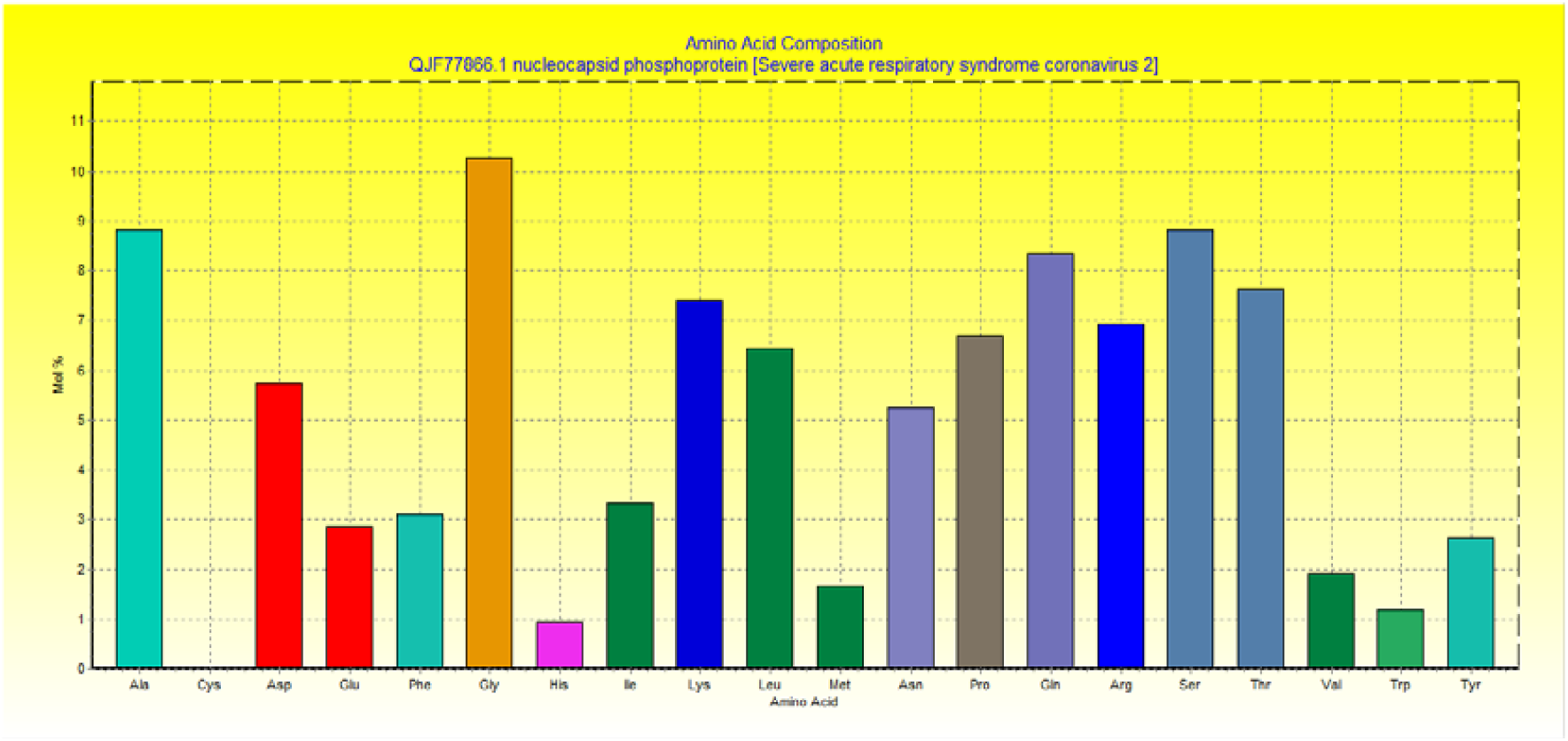
Amino acid composition of Nucleocapsid phosphoprotein. Amino acid distribution histogram for 12 mjor proteins of SARS-CoV-2 proteome

**Figure 2 (L).**
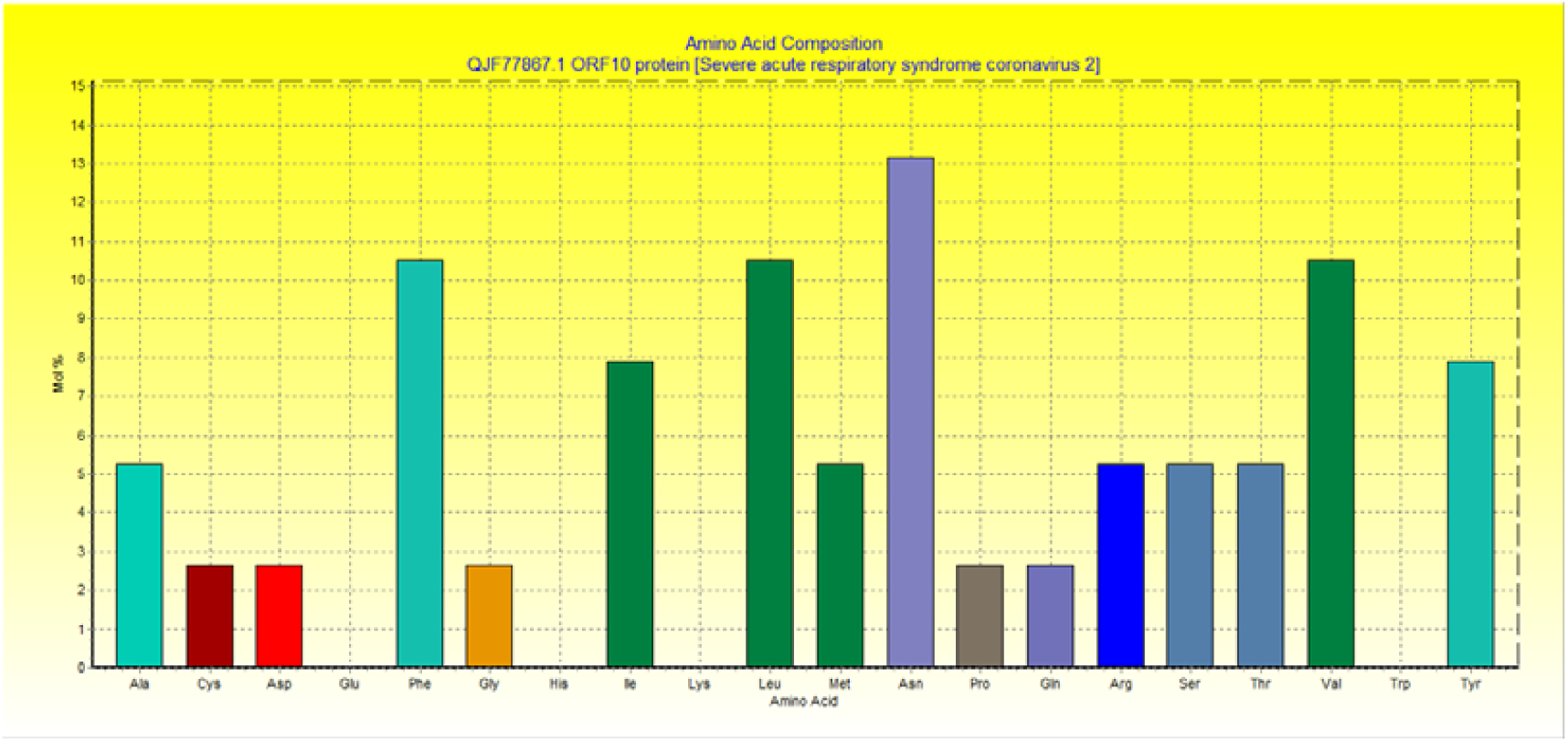
Amino acid composition of ORF10 protein. Amino acid distribution histogram for 12 mjor proteins of SARS-CoV-2 proteome

InterPro classification of protein families ORF1a polyprotein belongs to Non-structural protein NSP1, betacoronavirus (IPR021590) family. It is involved in biological processes-viral genome replication (*GO: 0019079*), viral protein processing (*GO: 0019082*) and proteolysis (*GO:* 0006508).

The molecular functions of ORF1a polyprotein are omega peptidase activity (*GO* :0008242), cysteine-type endopeptidase activity (*GO* :0004197), transferase activity (*GO* :0016740), RNA binding (*GO* :0003723), nucleic acid binding (*GO* :0003676), Zinc ion binding (*GO* :0008270), cysteine-type peptidase activity (*GO* :0008234), RNA-directed 5’-3’ RNA polymerase activity (*GO* :0003968), and single-stranded RNA binding (*GO* :0003727).

Hits for all PROSITE (release 2020_02) motifs on sequence of ORF1a (YP_009725295-1) has found 4 hits (Table 4)- (i) Macro domain profile :PS51154|MACRO (profile) Macro domain profile :1025 - 1194: score=18.014; (ii) Peptidase family C16 domain profile : PS51124|PEPTIDASE_C16 (profile) Peptidase family C16 domain profile :1634 - 1898: score=60.973; (iii) Coronavirus main protease (M-pro) domain profile : PS51442|M_PRO (profile) Coronavirus main protease (M-pro) domain profile : 3264 - 3569: score=154.193; (iv) Carbamoyl-phosphate synthase subdomain signature 2: >PS00867|CPSASE_2 (pattern) Carbamoyl-phosphate synthase subdomain signature 2 : 2061 - 2068: [confidence level: (−1)].

The ORF1ab polyprotein is protein complex of 15 proteins namely NSP1, NSP2, NSP3, NSP4, 3C-like proteinase, NSP6, NSP7, NSP8, NSP9, NSP10, RNA-dependent RNA polymerase, Helicase, 3’-to-5’ exonuclease, EndoRNAse and 2’-O-ribose methyltransferase. Orf1ab polyprotein is a multifunctional protein involved in the transcription and replication of viral RNAs. Predicted functions of SARS-CoV-2 proteome with respective ProFunc score has been listed in Table 4 (**Annexure-I; Supplementary file**).

### 1. Leader protein (NSP1) (PF11501)

Non-structural protein NSP1 (IPR021590) is the N-terminal cleavage product from the viral replicase that mediates RNA replication and processing (Almeida *et al*., 2007). ProMotif results revealed that the structure of NSP1 protein has **2** sheets, 5 beta hairpins,4 beta bulges,7 strands,5 helices,1 helix-helix interacs,22 beta turns and 3 gamma turns (Figure 3 A). Physicochemical parameter analysis computed that NSP1 has theoretical Isoelectric point (pI) 5.36, Instability index 28.83, Aliphatic index 89.72 and Grand average of hydropathicity - 0.378 (Table 3).

**Figure 3A.**
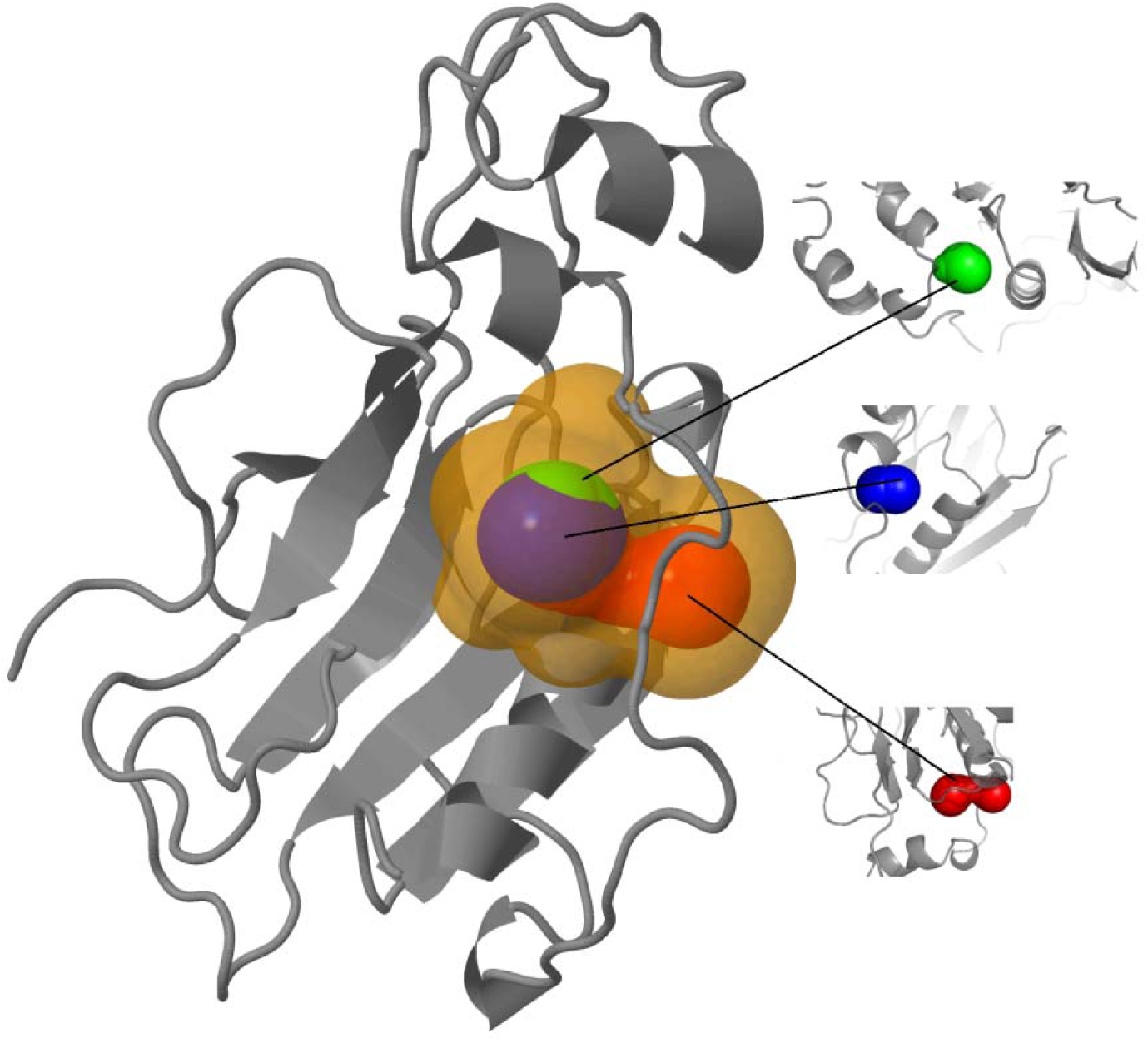
NSP1. The 15 predicted protein Structures of SARS-CoV-2 proteome along with estimated best tunnels.

**Figure 3B.**
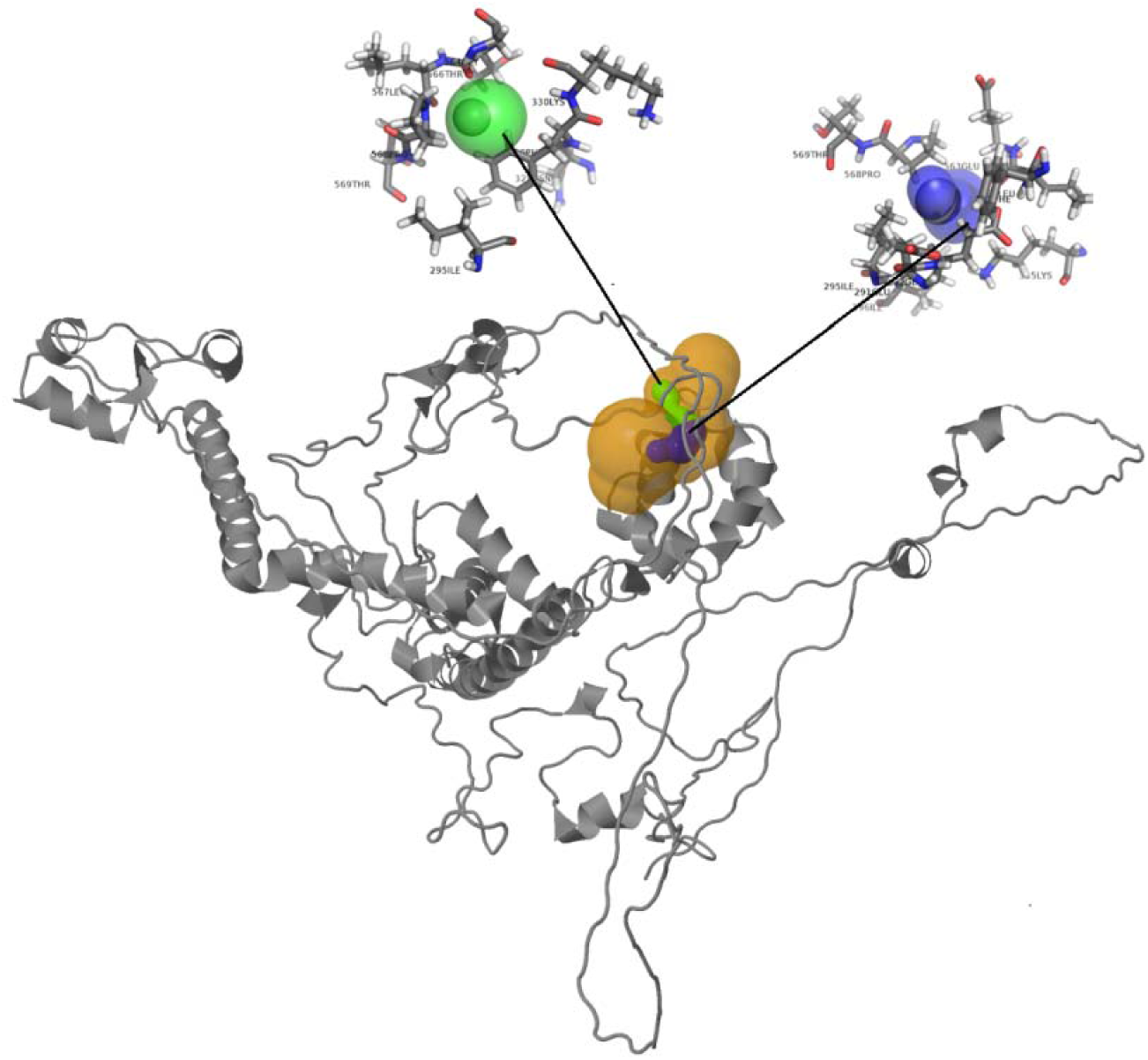
NSP2. The 15 predicted protein Structures of SARS-CoV-2 proteome along with estimated best tunnels.

**Figure 3C.**
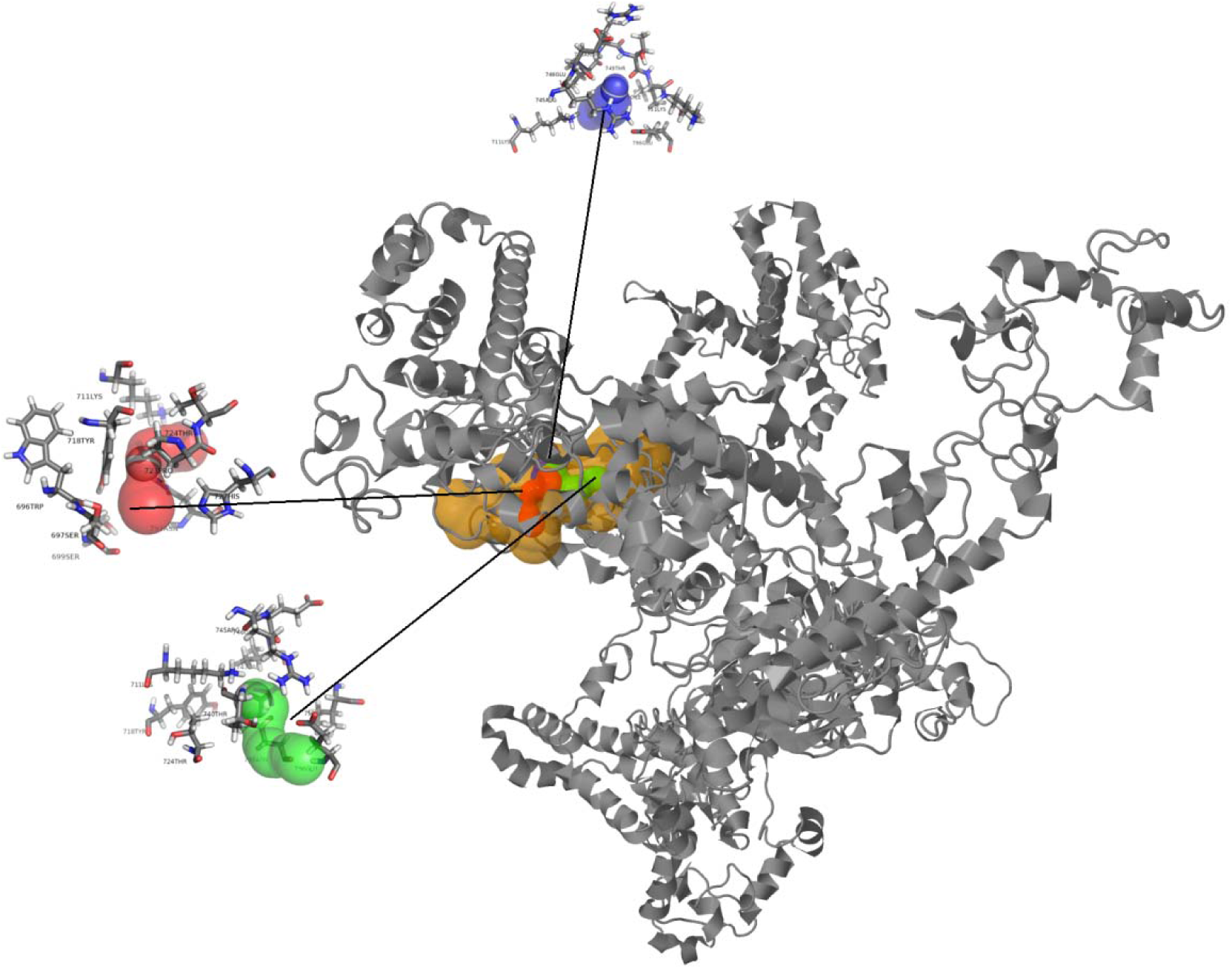
NSP3. The 15 predicted protein Structures of SARS-CoV-2 proteome along with estimated best tunnels.

**Figure 3D.**
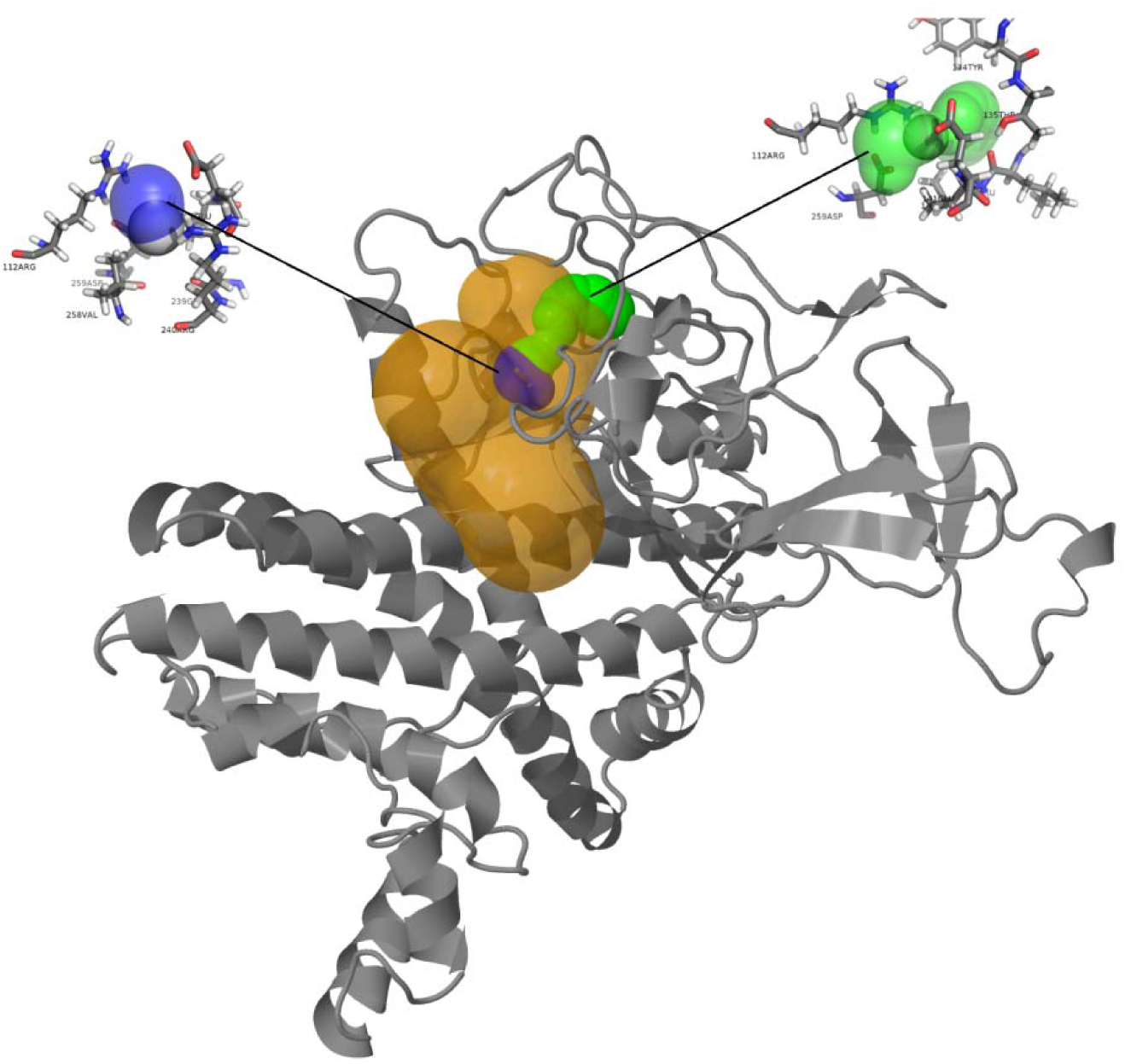
NSP4. The 15 predicted protein Structures of SARS-CoV-2 proteome along with estimated best tunnels.

**Figure 3E.**
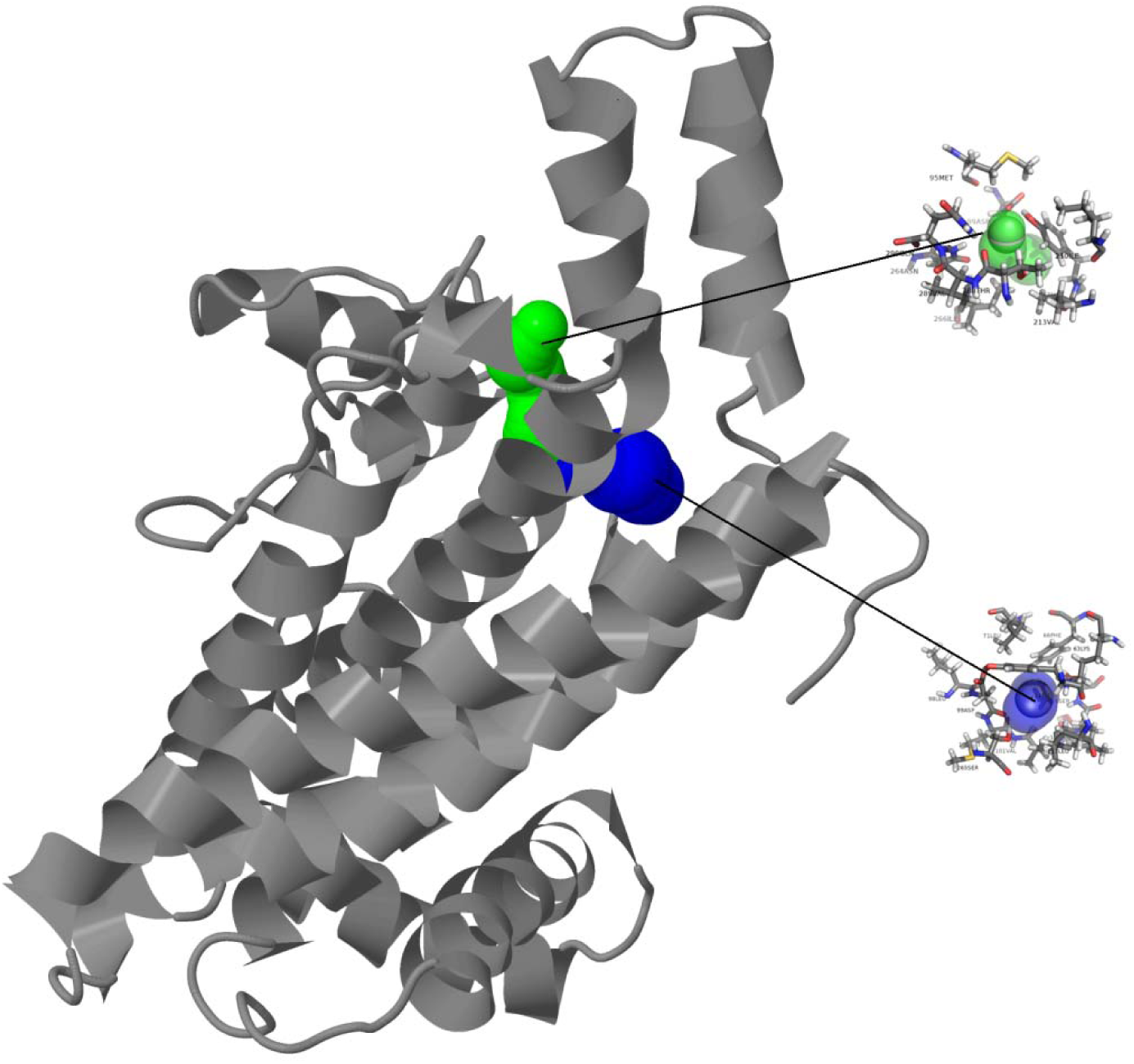
NSP6. The 15 predicted protein Structures of SARS-CoV-2 proteome along with estimated best tunnels.

**Figure 3F.**
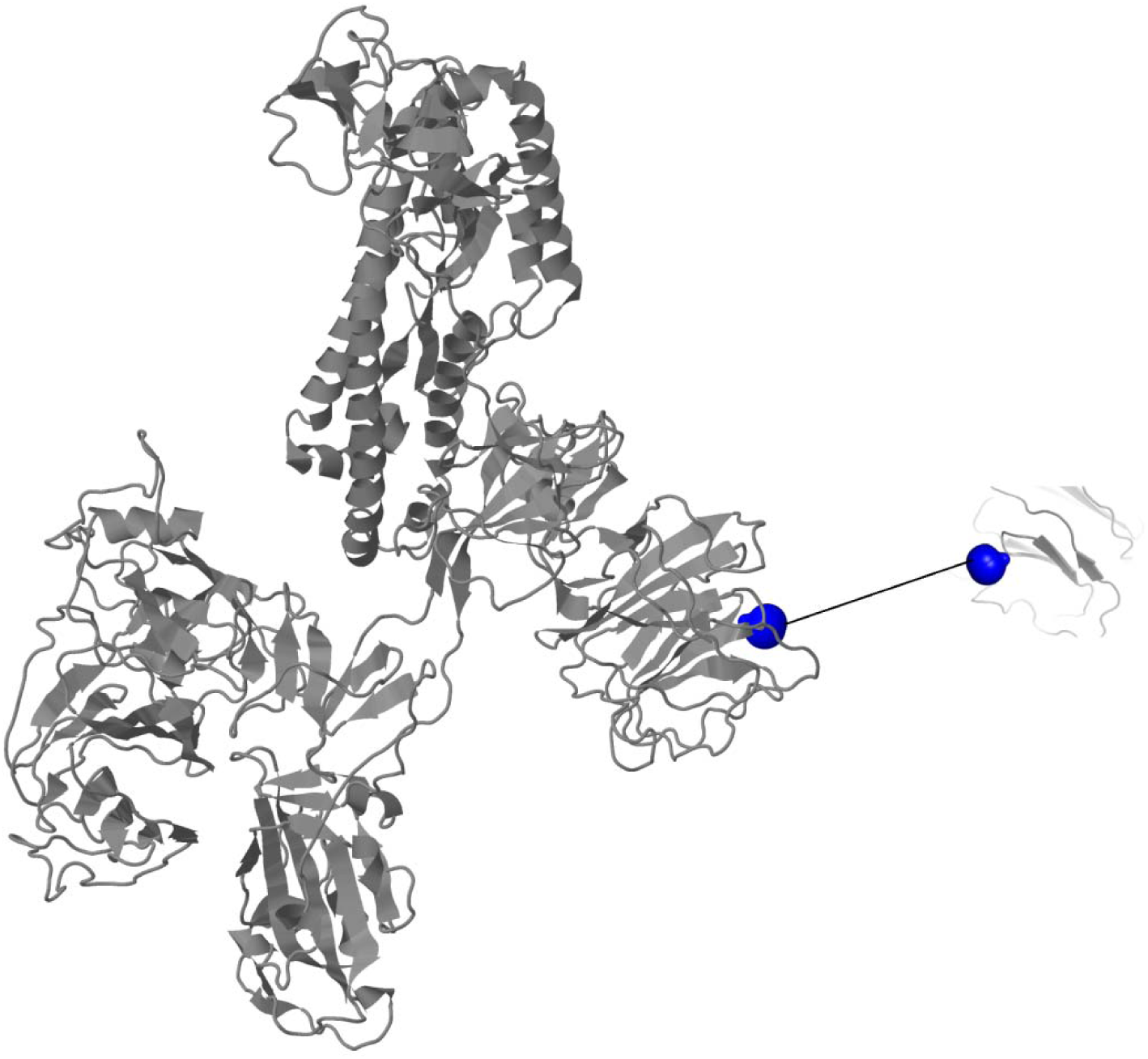
Surface Glycoprotein. The 15 predicted protein Structures of SARS-CoV-2 proteome along with estimated best tunnels.

**Figure 3G.**
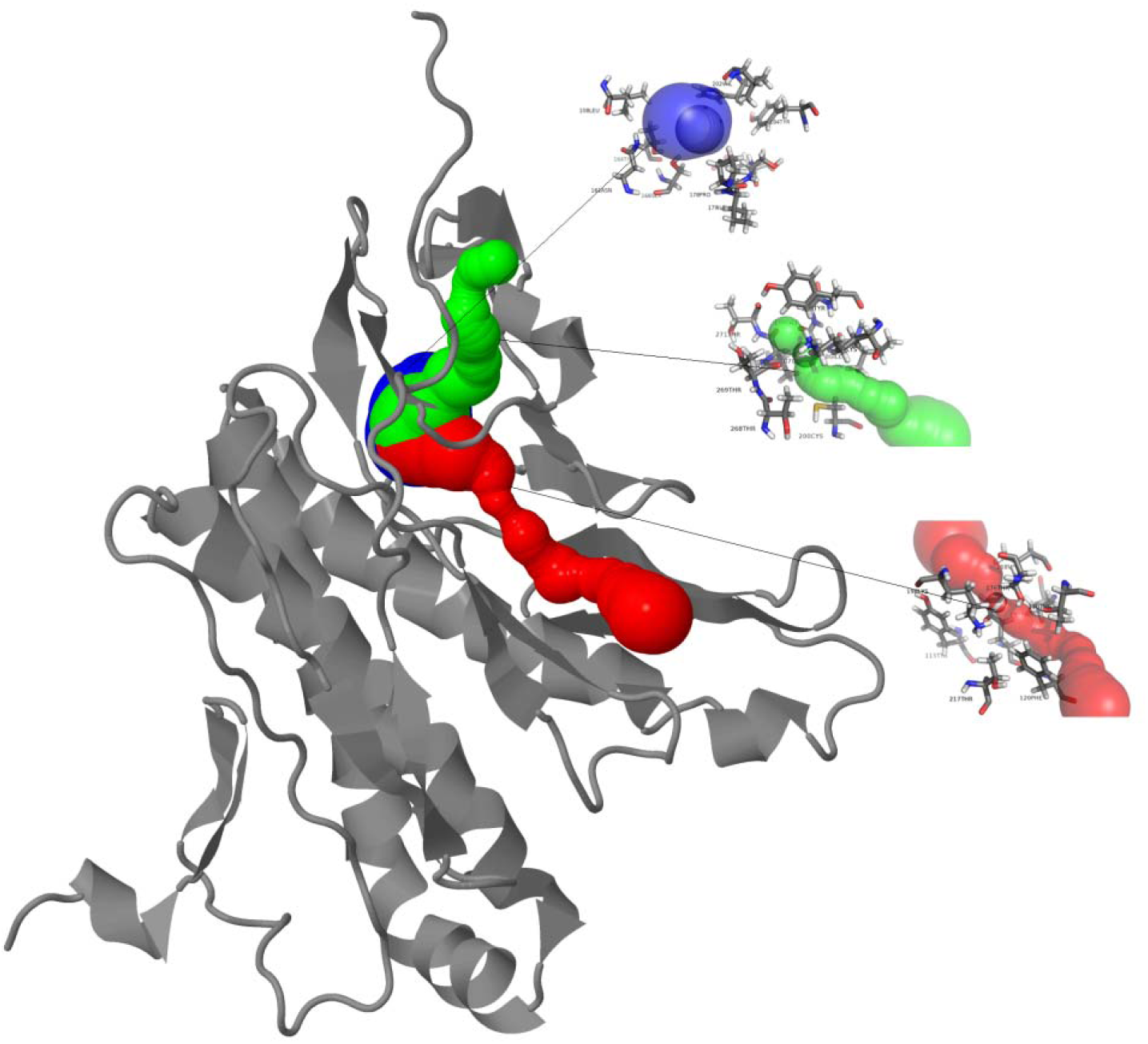
ORF3a protein. The 15 predicted protein Structures of SARS-CoV-2 proteome along with estimated best tunnels.

**Figure 3H.**
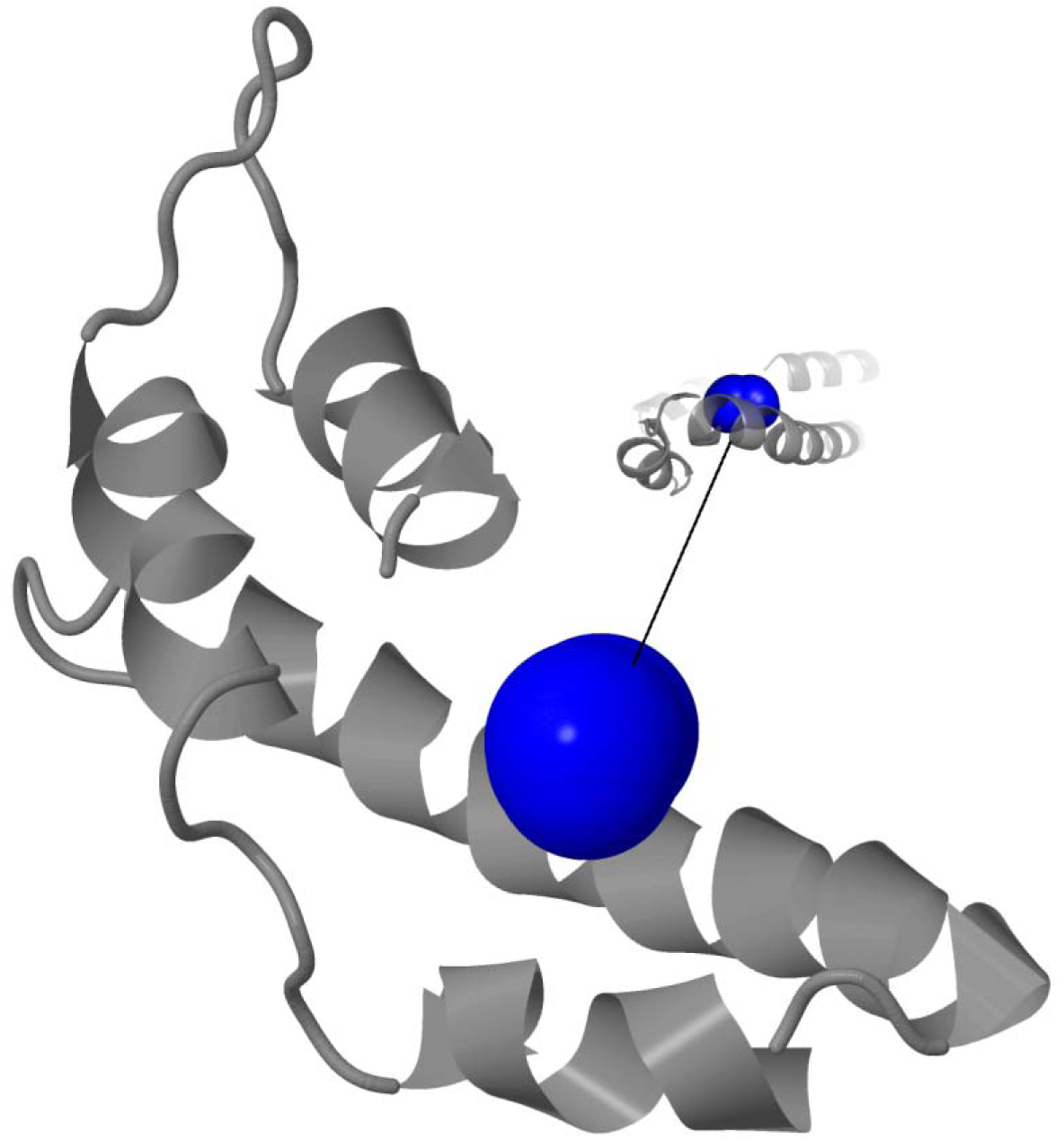
Envelope protein. The 15 predicted protein Structures of SARS-CoV-2 proteome along with estimated best tunnels.

**Figure 3I.**
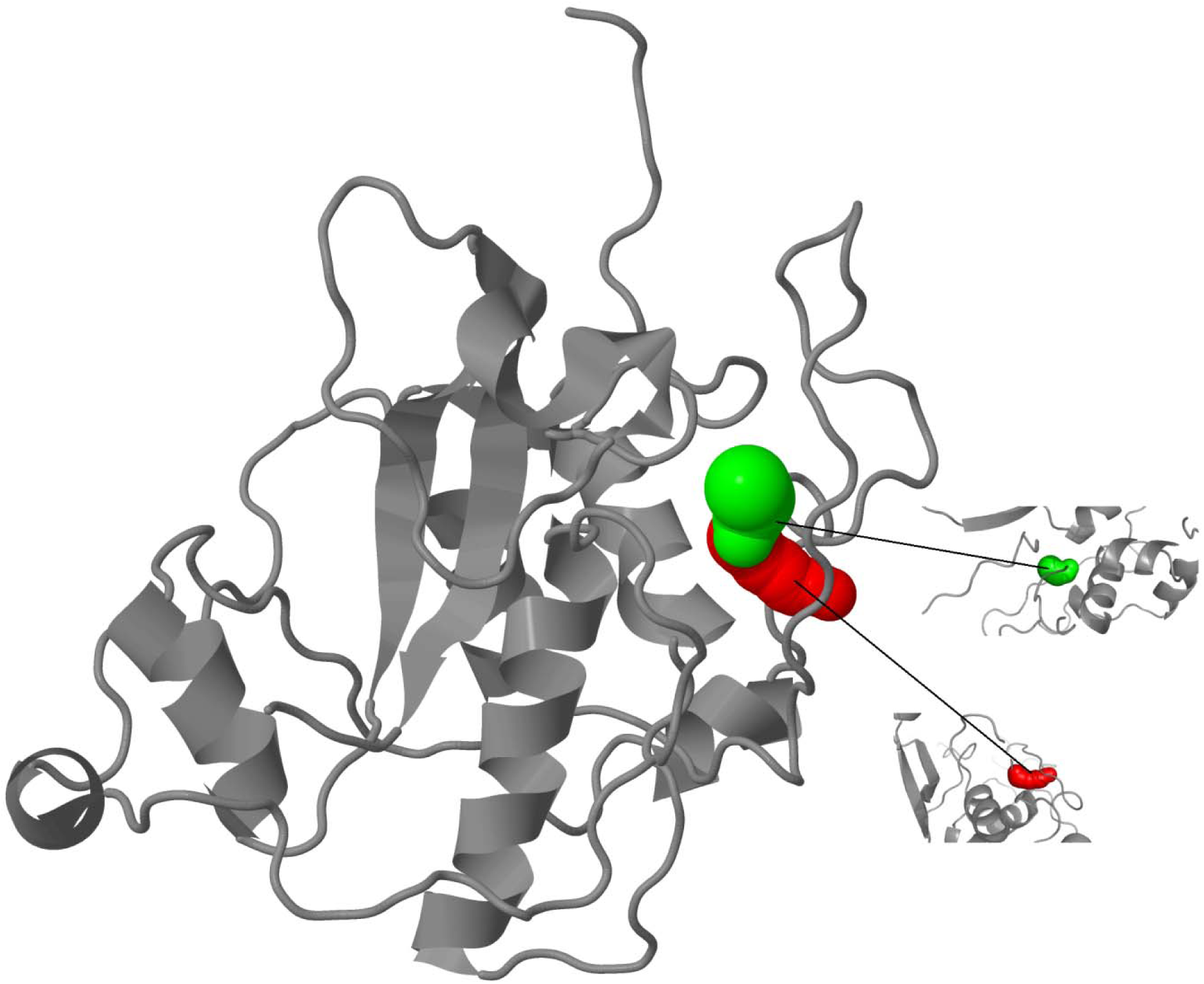
Membrane glycoprotein. The 15 predicted protein Structures of SARS-CoV-2 proteome along with estimated best tunnels.

**Figure 3J.**
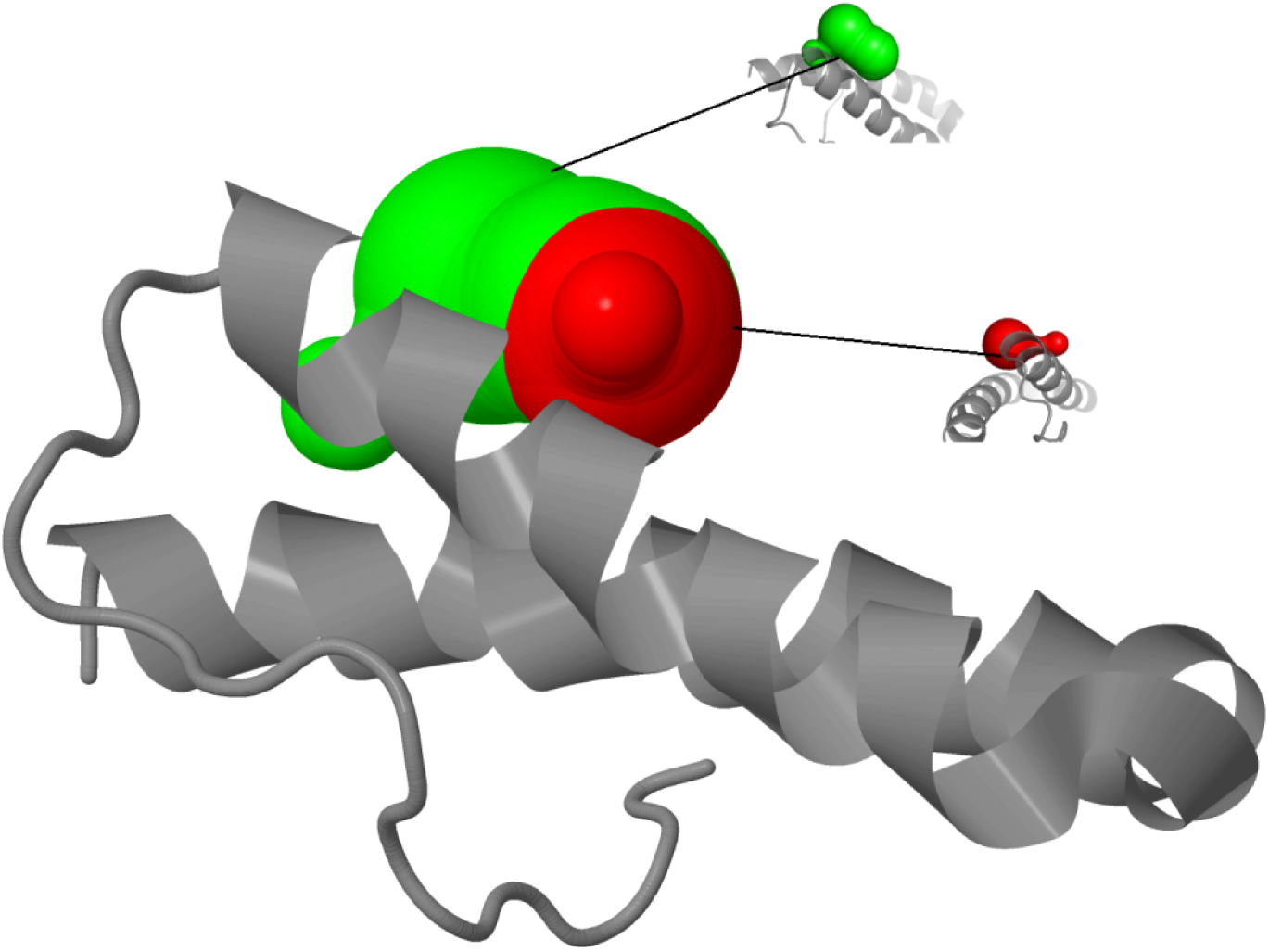
ORF6 protein. The 15 predicted protein Structures of SARS-CoV-2 proteome along with estimated best tunnels.

**Figure 3K.**
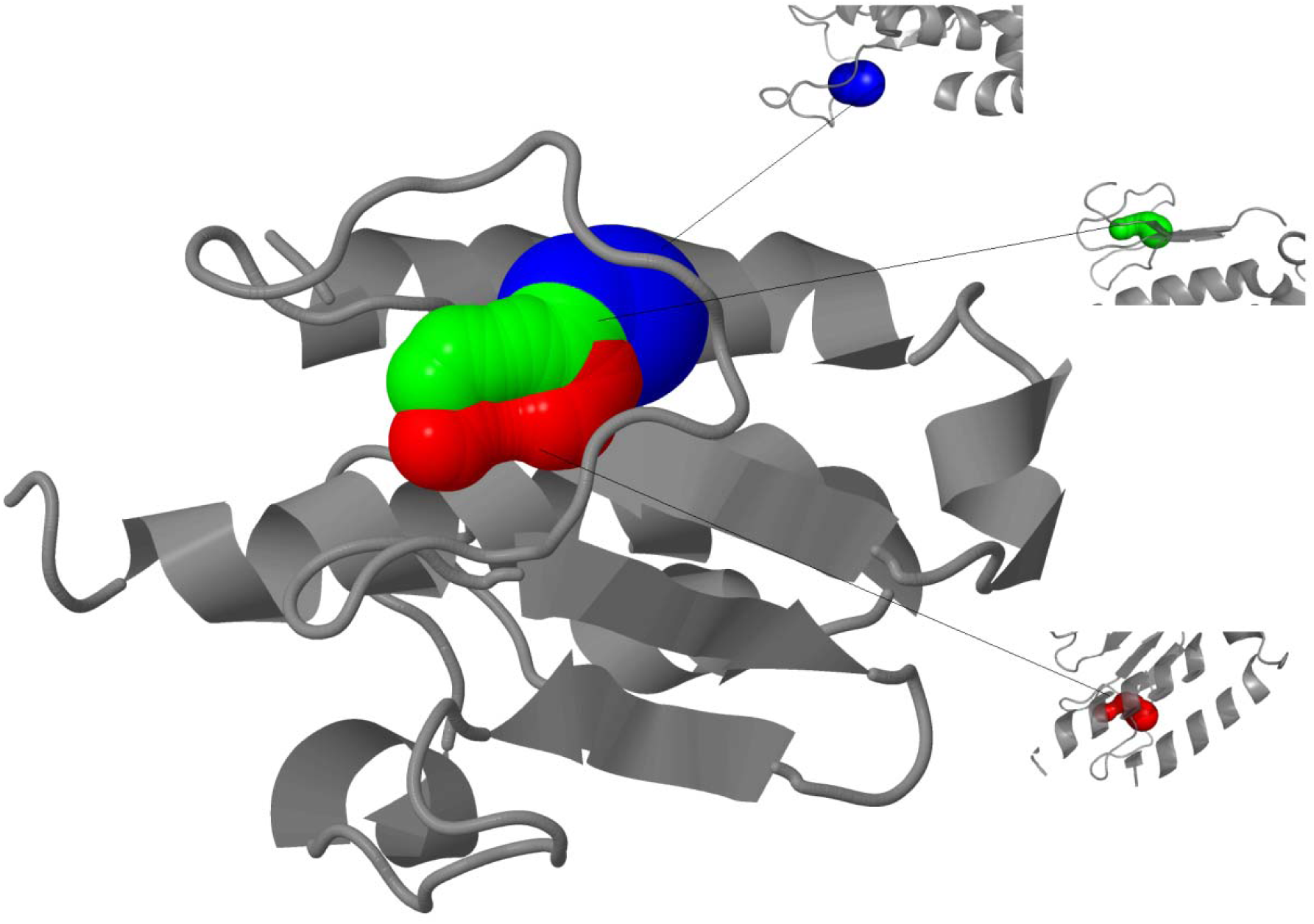
ORF7a protein. The 15 predicted protein Structures of SARS-CoV-2 proteome along with estimated best tunnels.

**Figure 3L.**
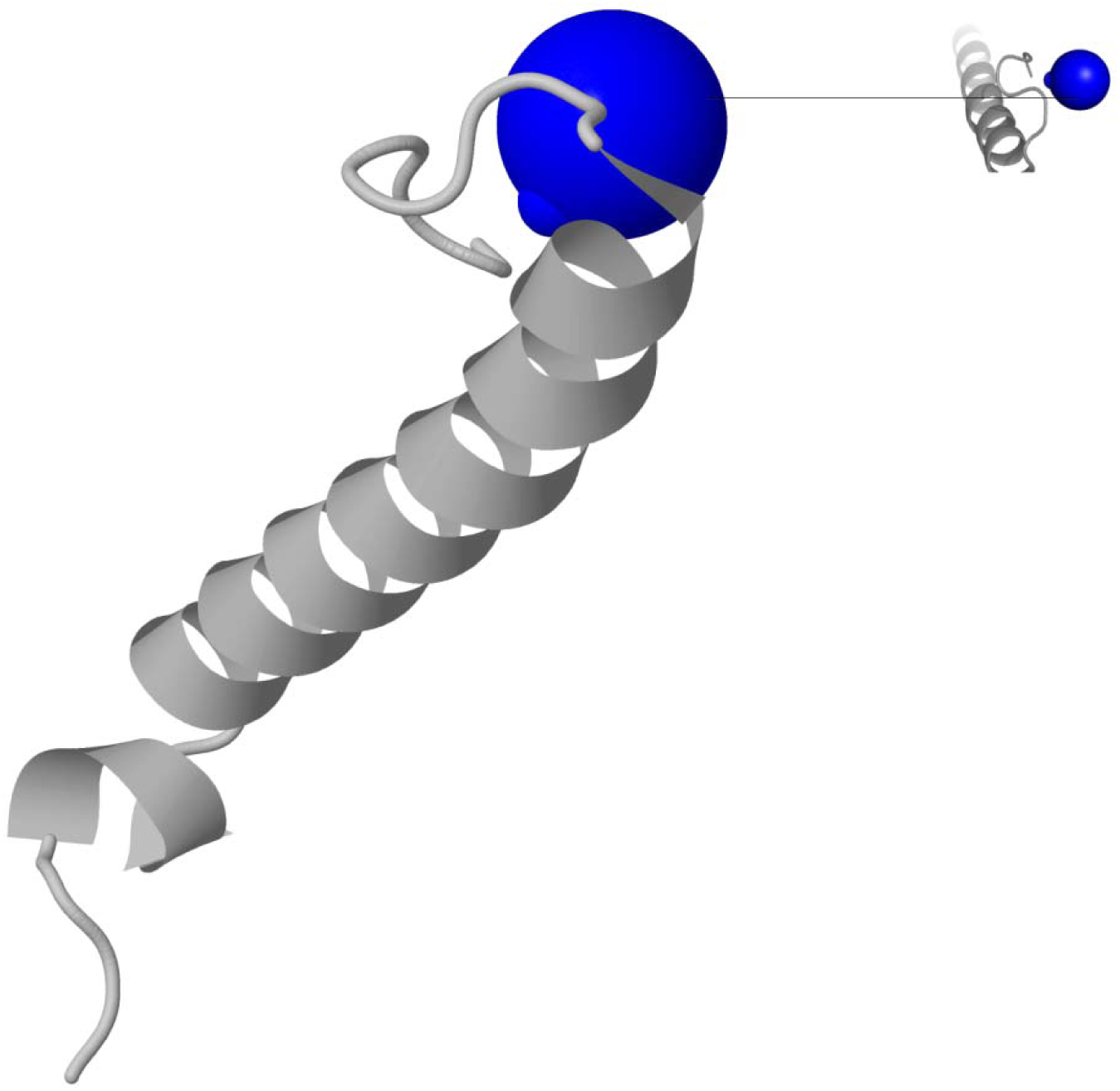
ORF 7b protein. The 15 predicted protein Structures of SARS-CoV-2 proteome along with estimated best tunnels.

**Figure 3M.**
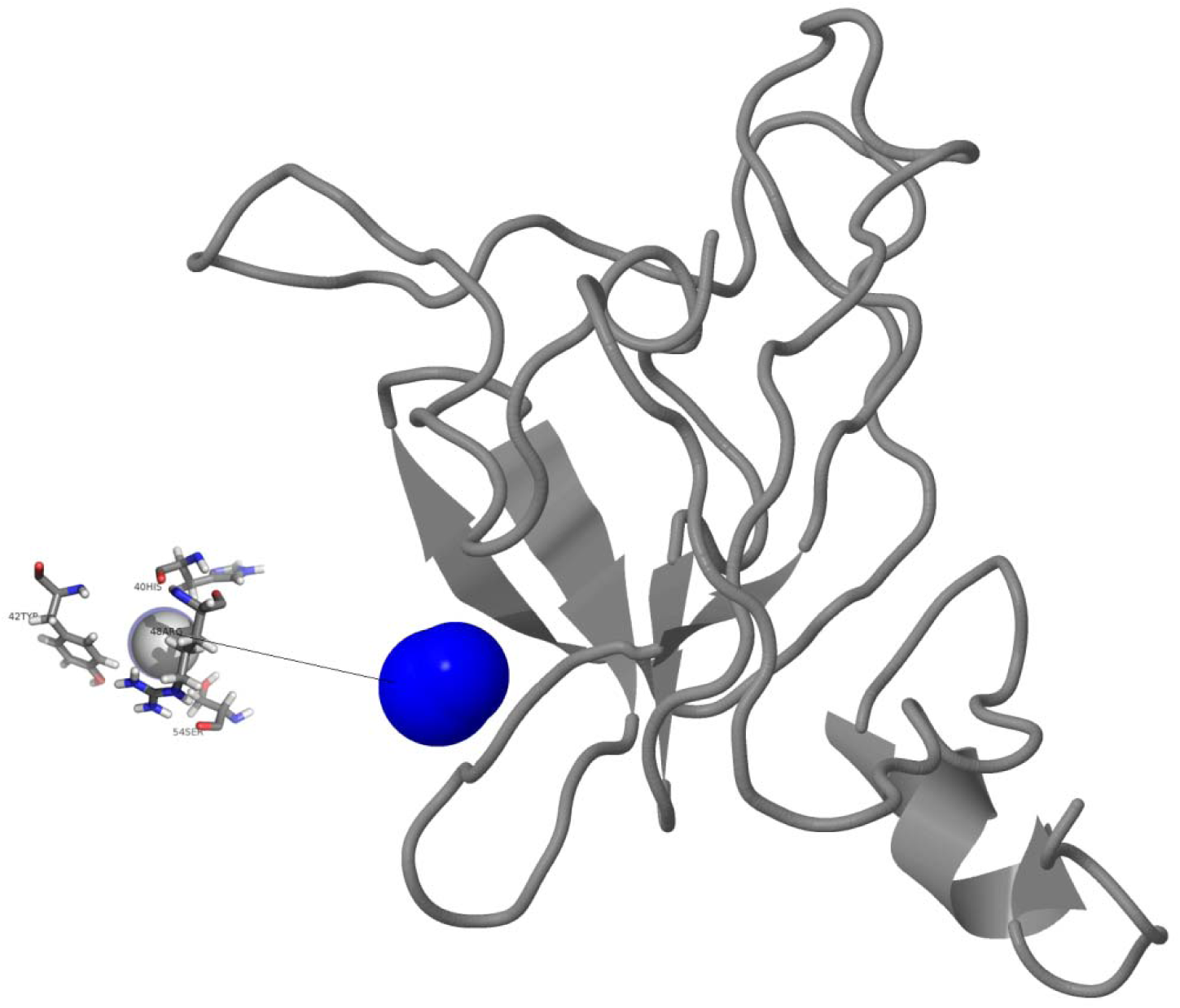
ORF 8 protein. The 15 predicted protein Structures of SARS-CoV-2 proteome along with estimated best tunnels.

**Figure 3N.**
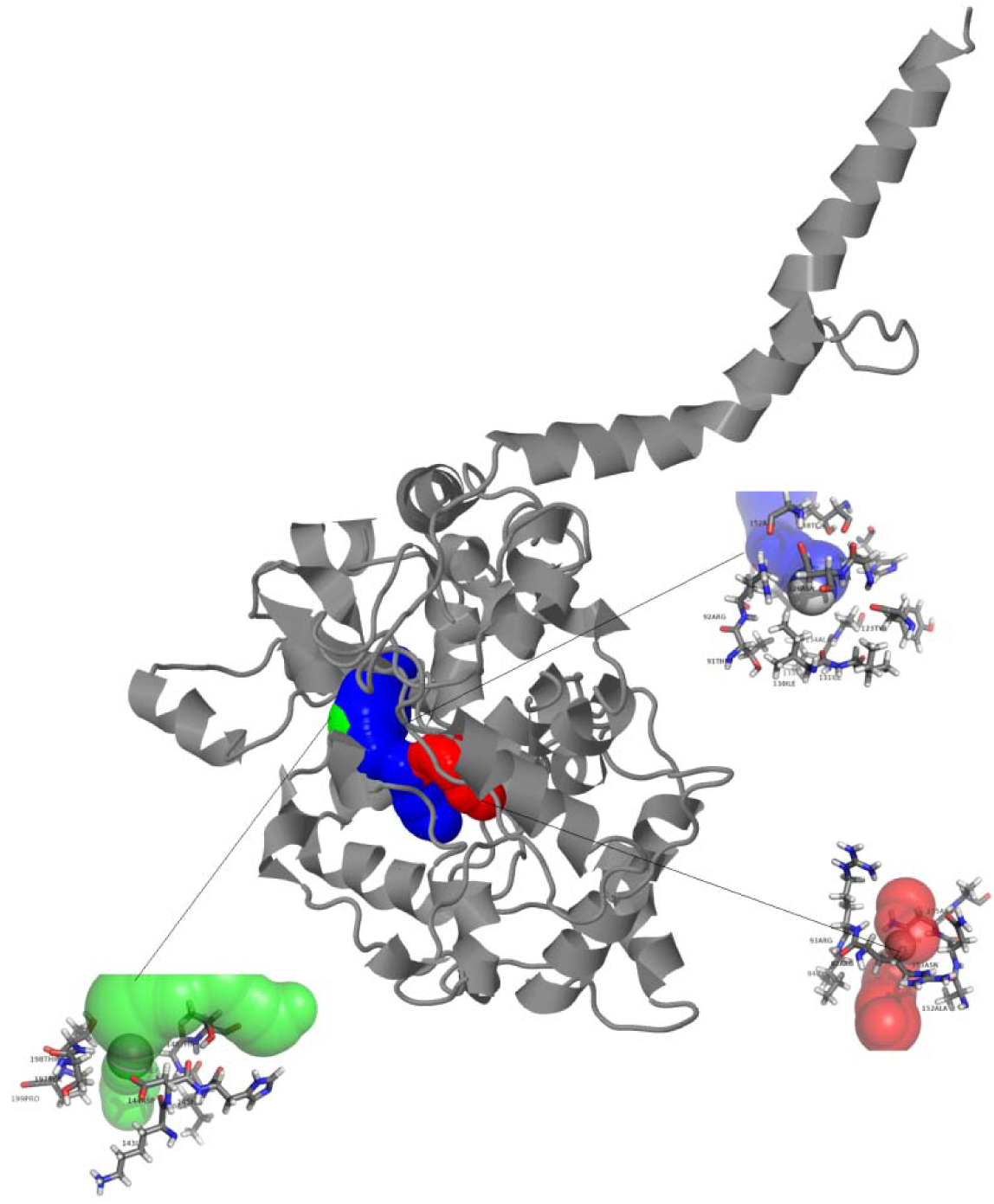
Nucleocapsid phosphoprotein. The 15 predicted protein Structures of SARS-CoV-2 proteome along with estimated best tunnels.

**Figure 3O.**
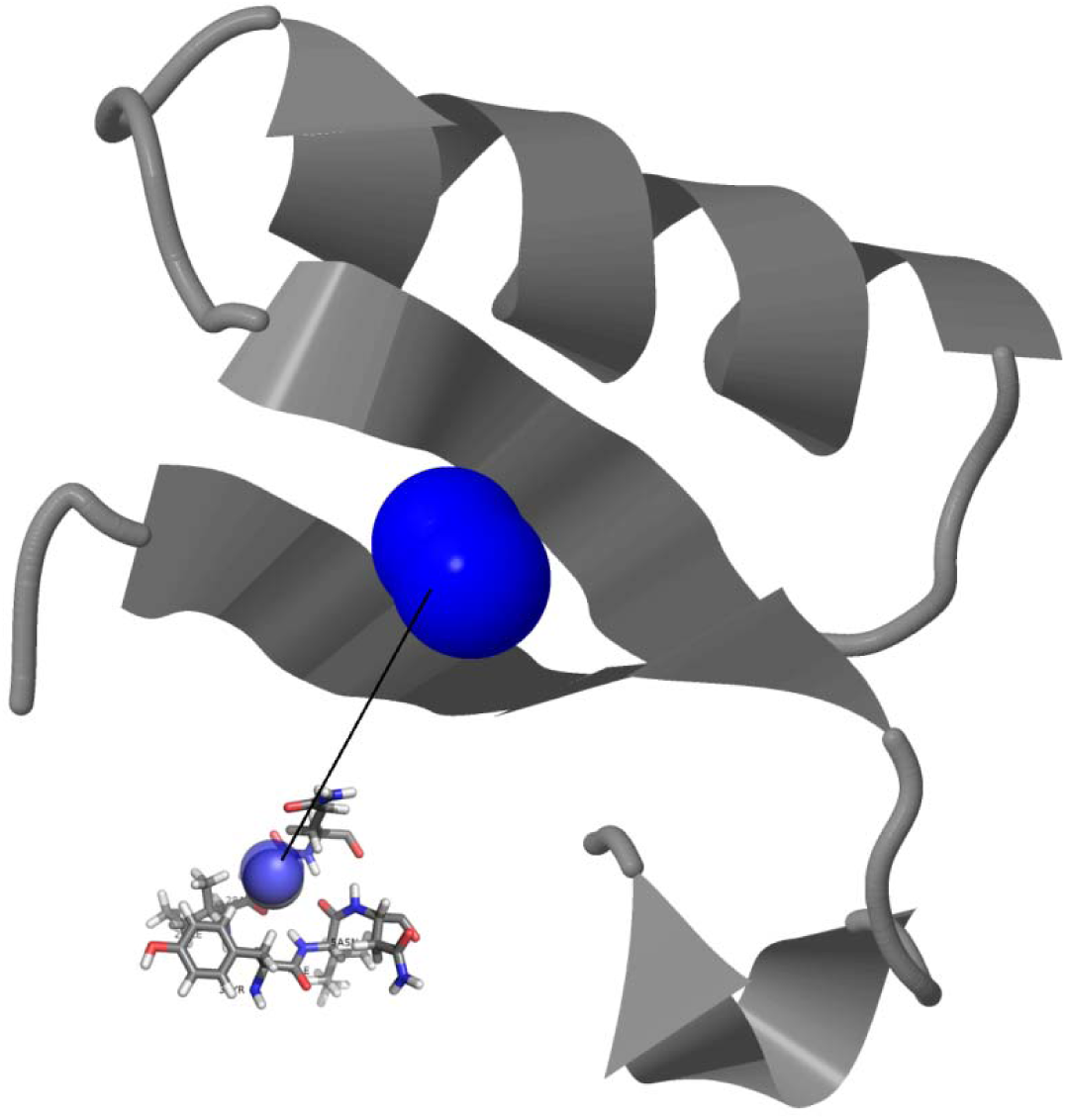
ORF10 protein. The 15 predicted protein Structures of SARS-CoV-2 proteome along with estimated best tunnels.

NSP1 binds to the 40S ribosomal subunit and inhibits translation, and it also induces a template-dependent endonucleolytic cleavage of host mRNAs (Kamitani *et al*., 2009). Structurally, NSP1 consists of a mixed parallel/antiparallel 6-stranded beta barrel with an alpha helix covering one end of the barrel and another helix alongside the barrel (Almeida *et al*., 2007). NSP1 also suppresses the host innate immune functions by inhibiting type I interferon expression and host antiviral signaling pathways (Almeida *et al*., 2007).

Tunnels analysis results have estimated three tunnels in NSP1. The best tunnel i.e.tunnel-1(blue) with bottleneck radius 1.9 Å, length 2.9 Å, distance to surface 2.4 Å, curvature 1.2, throughput 0.89 and number of residues 16; The tunnel-2 (green) with bottleneck radius 1.6 Å, length 7.3 Å, distance to surface 6.2 Å, curvature 1.2, throughput 0.81 and number of residues 22; The tunnel-3 (red) with bottleneck radius 1.1 Å, length 8.8 Å, distance to surface 7.3 Å, curvature 1.2, throughput 0.68 and number of residues 17 (Figure 3A).

### 2. Non-structural protein 2 (NSP2)

ProMotif evaluation demonstrated that the NSP2 has 2 sheets, 2 beta hairpins, 1 beta bulge, 4 strands; 23 helices; 15 helix-helix interacs; 40 beta turns; 10 gamma turns;1 disulphide (Figure 3B). NSP2 has pI at 6.25, Instability index 36.06, Aliphatic index 88.93 and Grand average of hydropathicity - 0.062 (Table 3). This protein may play a role in the modulation of host cell survival signaling pathway by interacting with host PHB and PHB2.

Tunnels analysis results have estimated two most active tunnels in NSP2. The tunnel-1(blue) with bottleneck radius 1.5 Å, length 5.4 Å, distance to surface 4.8 Å, curvature 1.1, throughput 0.82 and number of residues 19; The tunnel-2 (green) with bottleneck radius 1.1 Å, length 7.2 Å, distance to surface 6.6 Å, curvature 1.1, throughput 0.65 and number of residues 20 (Figure 3B).

### 3. Non-structural protein 3 (NSP3)

The structure of NSP3 has 6 sheets, 10 beta hairpins, 6 beta bulges, 23 strands, 139 helices, 130 helix-helix interacs, 169 beta turns, 45 gamma turns, and 6 disulphides after application of ProMotif (Figure 3C). NSP3 with pI 5.56 has Instability index 36.56, Aliphatic index 86.22 and Grand average of hydropathicity - 0.175 (Table 3).

Biological Process of NSP3 is proteolysis (GO: 0006508) and involves in molecular function such as single-stranded RNA binding activity (GO: 0003727), cysteine-type peptidase activity (GO: 0008234), RNA-directed 5’-3’ RNA polymerase activity (GO: 0003968) and nucleic acid binding (GO: 0003676).

A total of 7 tunnels have been estimated in the structure of NSP3. Out of which three are found to be with high throughput values. The tunnel-1(blue) with bottleneck radius 1.5 Å, length 6.3 Å, distance to surface 5.6 Å, curvature 1.1, throughput 0.80 and number of residues 15; The tunnel-2 (green) with bottleneck radius 1.7 Å, length 10.5 Å, distance to surface 8.5 Å, curvature 1.2, throughput 0.72 and number of residues 21; The tunnel-3 (red) with bottleneck radius 1.4 Å, length 11.1 Å, distance to surface 7.4 Å, curvature 1.5, throughput 0.68 and number of residues 20 (Figure 3B). (Figure 3C).

### 4. Non-structural protein 4 (NSP4)

The structure of NSP4 has 5 sheets, 6 beta hairpins, 3 beta bulges, 11 strands, 16 helices, 20 helix-helix interacs, 41 beta turns, 6 gamma turns (Figure 3D). NSP4 has pI at 7.16, Instability index 34.09, Aliphatic index 95.50 and Grand average of hydropathicity 0.343 (Table 3). NSP4 Participates in the assembly of virally-induced cytoplasmic double-membrane vesicles necessary for viral replication. This C-terminal domain (InterPro entry IPR032505) is predominantly alpha-helical, which may be involved in protein-protein interactions (Manolaridis *et al*., 2009).

Tunnels analysis results have estimated two most active tunnels in NSP 4. The tunnel-1(blue) with bottleneck radius 1.8 Å, length 1.9 Å, distance to surface 1.8 Å, curvature 1.0, throughput 0.91 and number of residues 7; The tunnel-2 (green) with bottleneck radius 1.5 Å, length 13.7 Å, distance to surface 9.4 Å, curvature 1.5, throughput 0.72 and number of residues 21 (Figure 3D).

### 5. 3C-like protein

Structure of 3C-like proteinase presented 2 sheets, 7 beta hairpins, 7 beta bulges, 13 strands, 8 helices, 9 helix-helix interacs, 28 beta turns and 2 gamma turns with pI 5.95 as well as Instability index 27.65, Aliphatic index 82.12 and Grand average of hydropathicity −0.019 (Table 3).

The biological process of 3C-like proteinase is the viral protein processing (*GO* :0019082). It resembles Peptidase*_C30* (PF05409) family which corresponds to Merops family C30. This polyprotein is a target of drug cinanserin, acting most likely via inhibition of the 3CL proteinase identified by deploying both a homology model and the crystallographic structure of the binding pocket of the enzyme (Cheng *et al.*, 2005).Cinanserin is an inhibitor of the 3C-Like Proteinase of Severe Acute Respiratory Syndrome Coronavirus and Strongly Reduces Virus Replication *in vitro* (Lili *et al*., 2005).

### 6. Non-structural protein 6 (NSP6)

NSP6 presented 1 sheet, 1 beta hairpin, 2 strands, 14 helices, 31 helix-helix interacs, 9 beta turns, 2 gamma turns (Figure 3E) along with pI value 9.11 Instability index 22.94, Aliphatic index 111.55 and Grand average of hydropathicity 0.790 (Table 3). Nsp6 may play a role in the initial induction of autophagosomes from host’s endoplasmic reticulum.

NS6 can increase the cellular gene synthesis and may induce apoptosis through c-Jun N-terminal kinase and Caspase-3 mediated stress (Cheng *et al*., 2015) could modulate host antiviral responses by inhibiting the synthesis and signalling of interferon-beta (IFN-beta) via two complementary pathways. One involves NS6 interaction with host N-Myc (and STAT) interactor (Nmi) protein inducing its degradation via ubiquitin proteasome pathway, suppressing Nmi enhanced IFN signalling. The other pathway suppresses the translocation of signal transducer and activator of transcription 1 (STAT1) and downstream IFN signalling (Cheng *et al*., 2015).

Tunnels analysis results have estimated two most active tunnels in Nsp 6 out of three tunnels. The tunnel-1(blue) with bottleneck radius 1.8 Å, length 7.8 Å, distance to surface 7.1 Å, curvature 1.1, throughput 0.81 and number of residues 20; The tunnel-2 (green) with bottleneck radius 1.4 Å, length 8.7 Å, distance to surface 7.0 Å, curvature 1.3, throughput 0.75 and number of residues 20 (Figure 3E).

### 7. Non-structural protein 7 (NSP7)

NSP7 is predominantly a alpha helical structure with 3 helices, 7 helix-helix interacs and 3 beta turns (PDB ID 7BV1_C). NSP7 has pI 5.18, Instability index 51.97, Aliphatic index 117.35 and Grand average of hydropathicity 0.199 (Table 3). NSP7 may have the function in activating RNA-synthesizing activity and it forms a hexadecamer with nsp8 that may participate in viral replication by acting as a primase (Xiao *et al*.,2012) InterPro scan has predicted its molecular functions as omega peptidase activity (*GO* :0008242), cysteine-type endopeptidase activity (*GO* :0004197) and transferase activity (*GO* :0016740). NSP 7 belongs to nsp7 (PF08716) has been implicated in viral RNA replication. Nonstructural Proteins 7 and 8 of Feline Coronavirus forms a 2:1 Heterotrimer that exhibits primer-independent RNA Polymerase Activity (Yibei *et al*., 2012).

### 8. Non-structural protein 8 (NSP8)

NSP8 has2 sheets, 2 beta hairpins, 1 beta bulge, 5 strands, 5 helices, 6 helix-helix interacs, 13 beta turns, 1 gamma turn (PDB ID 7BV1_B) carries pI value of 6.58, as well as Instability index 37.78, Aliphatic index 88.33 and Grand average of hydropathicity of −0.192 (Table 3). It forms a hexadecamer with nsp7 that may participate in viral replication by acting as a primase. Molecular Functions of NSP8 are scanned as transferase activity (GO: 0016740), cysteine-type endopeptidase activity (*GO* :0004197) and omega peptidase activity (*GO* :0008242).

NSP alone as a monomer structure may not be biologically relevant as it forms a hexadecameric supercomplex with nsp7 (Xiao *et al.*,2012). The dimensions of the central channel and positive electrostatic properties of the cylinder imply that it confers processivity on RNA-dependent RNA polymerase (Zhai *et al*., 2005).

### 9. Non-structural protein 9 (NSP9)

Nsp9 has single helix with 2 sheets, 5 beta hairpins, 4 beta bulges, 7 strands, 11 beta turns showed pI 9.10 along with Instability index 34.17, Aliphatic index 82.92 and Grand average of hydropathicity −0.227 (Table 3).It acts as a ssRNA-binding protein might be involved in viral replication and RNA binding activity (GO: 0003723). The NSP9 (PF08710) is a single-stranded RNA-binding viral protein likely to be involved in RNA synthesis (Egloff *et al*., 2004) and its structure comprises of a single beta barrel (Campanacci *et al*., 2003).

### 10. Non-structural protein 10 (NSP10)

The structure of NSP10 has exhibited 2 sheets, 1 beta hairpin, 5 strands, 6 helices, 3 helix-helix interacs, 13 beta turns, 1 gamma turn with pI value of 6.29 as well as Instability index 34.56, Aliphatic index 61.80 and Grand average of hydropathicity −0.068 (Table 3). It plays an essential role in viral mRNAs cap methylation.

The NSP 10 (PF09401) involves in biological process - viral genome replication (GO: 0019079) and RNA binding activity (GO: 0003723), zinc ion binding (GO: 0008270). A cluster of basic residues on the protein surface suggests a nucleic acid-binding function, interacting selectively and non-covalently with an RNA molecule or a portion thereof. NSP10 contains two zinc binding motifs and forms two anti-parallel helices which are stacked against an irregular beta sheet (Joseph *et al*., 2006). Nsp10 binds to nsp16 through an activation surface area in nsp10, and the resulting complex exhibits RNA cap (nucleoside-2’-O)-methyltransferase activity.

### 11. RNA-dependent RNA polymerase (Pol/RdRp)

Pol/RdRp showed 8 sheets, 8 beta hairpins, 1 psi loop, 2 beta bulges, 22 strands, 41 helices, 58 helix-helix interacs, 91 beta turns and 16 gamma turns, is associated with replication and transcription of the viral RNA genome. RNA-dependent RNA polymerase has pI value of 6.14 along with Instability index 28.32, Aliphatic index 78.43 and Grand average of hydropathicity −0.224 (Table 3).

The biological process of Pol/RdRp (*Corona_RPol_N* _PF06478) has been with transcription, DNA-templated (GO: 0006351), and viral RNA genome replication (*GO:* 0039694). The molecular functions are predicted as RNA-directed 5’-3’ RNA polymerase activity (*GO* : 0003968), RNA binding (*GO* :0003723) and ATP binding (*GO* :0005524). Coronavirus RPol N-terminus family covers the N-terminal region of the coronavirus RNA-directed RNA polymerase. The nsp7 and nsp8 activate and confer processivity to the RNA-synthesizing activity of Pol (Kirchdoerfer and Ward, 2019).

### 12. Helicase

The structure of Helicase has been shown with 8 sheets, 1 beta alpha beta unit, 7 beta hairpins, 5 beta bulges, 26 strands, 19 helices, 16 helix-helix interacs, 92 beta turns, 15 gamma turns along with pI 8.66, Instability index 33.31, Aliphatic index 84.49 and Grand average of hydropathicity −0.096 (Table 3).

Helicase protein has molecular functions-zinc ion binding (GO: 0008270) and ATP binding (GO: 0005524). Multi-functional protein Helicase is associated with a zinc-binding domain in N-terminus displaying RNA and DNA duplex-unwinding activities with 5’ to 3’ polarity. Activity of helicase is also dependent on magnesium (By Similarity).

### 13. 3’-to-5’ exonuclease/NSP14

3’-to-5’ exonuclease has 6 sheets, 8 beta hairpins, 3 beta bulges, 23 strands, 13 helices, 10 helix-helix interacs, 60 beta turns. 3’-to-5’ with pI 7.80, Instability index 28.85, Aliphatic index 78.96 and Grand average of hydropathicity −0.134 (Table 3). The two possible activities NSP14 include exoribonuclease activity acting on both ssRNA and dsRNA in a 3’ to 5’ direction and a N7-guanine methyltransferase activity.

### 14. endoRNAse/ NSP15

NSP15 demonstrated 7 sheets, 1 beta alpha beta unit, 9 beta hairpins, 6 beta bulges, 21 strands, 10 helices, 8 helix-helix interacs, 37 beta turns and 2 gamma turns with pI 5.06, Instability index 36.28, Aliphatic index 95.09 and Grand average of hydropathicity −0.076 (Table 3). endoRNAse is a Mn(2+)-dependent, uridylate-specific enzyme, which leaves 2’-3’-cyclic phosphates 5’ to the cleaved bond (Ricagno *et al*., 2006.)

### 15. 2’-O-ribose methyltransferase/NSP16

NSP16 carried 3 sheets, 3 beta alpha beta units, 1 beta hairpin, 2 beta bulges, 12 strands, 12 helices, 6 helix-helix interacs,15 beta turns and 4 gamma turns along with pI 7.59, Instability index 26.11, Aliphatic index 90.64 and Grand average of hydropathicity −0.086 (Table 3).

2’-O-ribose methyltransferase belongs to *NSP16* family (PF06460). The SARS-CoV RNA cap SAM-dependent (nucleoside-2’-O-)-methyltransferase (2’-O-MTase) is a heterodimer comprising SARS-CoV nsp10 and nsp16. Nsp16 adopts a typical fold of the S-adenosylmethionine-dependent methyltransferase (SAM) family as defined initially for the catechol O-MTase but it lacks several elements of the canonical MTase fold, such as helices B and C. The 2’-O-ribose methyltransferase (nsp16) topology matches those of dengue virus NS5 N-terminal domain and of vaccinia virus VP39 MTases (Chen *et al*., 2011).

### 16. Surface glycoprotein (spike glycoprotein)

Surface glycoprotein **(spike glycoprotein)** has presented 13 sheets, 18 beta hairpins, 18 beta bulges, 52 strands, 22 helices, 29 helix-helix interacs, 76 beta turns, 16 gamma turns and 12 disulphides (Figure 3F). Surface glycoprotein (Length = 1273 amino acids; Molecular Weight = 141.113kDa) is rich in Leucine (8.48%) and Serine (7.78) %). Surface glycoprotein has pI 6.32, Instability index 32.86, Aliphatic index 84.67 and Grand average of hydropathicity −0.077 (Figure 2C; Table 3).

Surface glycoprotein involves in two important biological processes i.e. receptor-mediated virion attachment to host cell (GO: 0046813) and membrane fusion (GO: 0061025). **Spike protein S1** attaches the virion to the cell membrane by interacting with host receptor, initiating the infection. Binding to human ACE2 and CLEC4M/DC-SIGNR receptors and internalization of the virus into the endosomes of the host cell induces conformational changes in the S glycoprotein. **Spike protein S2** mediates fusion of the virion and cellular membrane by acting as a class I viral fusion protein. **Spike protein S2’** Acts as a viral fusion peptide which is unmasked following S2 cleavage occurring upon virus endocytosis. It is a part of cellular components viral envelope (GO: 0019031) and integral component of membrane (GO: 0016021).

The estimated high throughput tunnel-1 (blue) in Surface glycoprotein is with bottleneck radius 1.1 Å, length 4.9 Å, distance to surface 3.0 Å, curvature 1.7, throughput 0.81 and number of residues 7 (Figure 3F).

### 17. ORF3a protein

ORF3a protein (**Papain-lke protease**) has 7 sheets, 8 beta hairpins, 2 beta bulges, 15 strands, 6 helices, 3 helix-helix interacs, 28 beta turns, 2 gamma turns and 1 disulphide (Figure 3G). ORF3a protein (Molecular Weight = 31121.29 Daltons) is rich in Leucine (10.91%), Valine (9.09%), Threonine (8.73%) and Serine (8.00%). ORF3a protein has theoretical Isoelectric point (pI) 5.55, Instability index 32.96, Aliphatic index 103.42 and Grand average of hydropathicity 0.275 (Figure 2D; Table 3).

The protein belongs to family Protein 3a, betacoronavirus (IPR024407). Protein 3a encoded by Orf3/3a, also known as X1, which forms homotetrameric potassium, sodium or calcium sensitive ion channels (viroporin) and may modulate virus release. It has also been shown to up-regulate expression of fibrinogen subunits FGA, FGB and FGG in host lung epithelial cells (Shen *et al*., 2005; Lu *et al*., 2006).

Tunnels analysis results have estimated three tunnels in ORF3a. The best tunnel i.e.tunnel-1(blue) with bottleneck radius 2.5 Å, length 5.8 Å, distance to surface 5.1 Å, curvature 1.1, throughput 0.90 and number of residues 22; The tunnel-2 (green) with bottleneck radius 1.1 Å, length 21.4 Å, distance to surface 18.3 Å, curvature 1.2, throughput 0.45 and number of residues 38, The tunnel-3 (red) with bottleneck radius 0.9 Å, length 27.0 Å, distance to surface 22.3 Å, curvature 1.2, throughput 0.37and number of residues 47 (Figure 3G).

3a protein is a pro-apoptosis-inducing protein. It localises to the endoplasmic reticulum (ER)-Golgi compartment. SARS-CoV causes apoptosis of infected cells through NLRP3 inflammasome activation, as ORF3a is a potent activator of the signals required for this activation, pro-IL-1beta gene transcription and protein maturation. This protein also promotes the ubiquitination of apoptosis-associated speck-like protein containing a caspase recruitment domain (ASC) mediated by its interaction with TNF receptor-associated factor 3 (TRAF3). The expression of ORF3a induces NF-kappa B activation and up-regulates fibrinogen secretion with the consequent high cytokine production (Yu *et al*., 2004; Lu *et al*., 2006). Another apoptosis mechanism described for this protein is the activation of the PERK pathway of unfolded protein response (UPR), which causes phosphorylation of eIF2alpha and leads to reduced translation of cellular proteins as well as the activation of pro-apoptotic downstream effectors (i.e ATF4, CHOP) (Minakshi *et al*., 2009).

### 18. Envelope protein (E protein)

Envelope protein (E protein) has 4 helices, 2 helix-helix interacs and 3 beta turns (Figure 3H). Envelope protein (Molecular Weight = 8364.59 Daltons) is rich in Leucine (18.67%) and Valine (17.33%). Envelope protein has pI 8.57, Instability index 38.68, Aliphatic index 144.00 and Grand average of hydropathicity 1.128 (Figure 2E; Table 3).

The Envelope protein belongs to family of Envelope small membrane protein, coronavirus (IPR003873) and Envelope small membrane protein, betacoronavirus (IPR043506). It plays a central role in virus morphogenesis and assembly. Biological Process of Envelope protein is pore formation by virus in membrane of host cell (GO: 0039707). The estimated high throughput tunnel-1 (blue) in envelope protein is with bottleneck radius 2.4 Å, length 4.2 Å, distance to surface 3.4 Å, curvature 1.2, throughput 0.92 and number of residues 7 (Figure 3H).

E proteins are well conserved among Coronavirus strains. They are small, integral membrane proteins involved in several aspects of the virus’ life cycle, such as assembly, budding, envelope formation, and pathogenesis (Schoeman and Fielding, 2019). E protein acts as a viroporin by oligomerizing after insertion in host membranes to create a hydrophilic pore that allows ion transport (Madan *et al*., 2005; Surya *et al*., 2015).

SARS-CoV E protein forms a Ca^2+^ permeable channel in the endoplasmic reticulum Golgi apparatus intermediate compartment (ERGIC)/Golgi membranes. The E protein ion channel activity alters Ca^2+^ homeostasis within cells boosting the activation of the NLRP3 inflammasome, which leads to the overproduction of IL-1beta. SARS-CoV overstimulates the NF-kappaB inflammatory pathway and interacts with the cellular protein syntenin, triggering p38 MARK activation. These signalling cascades result in exacerbated inflammation and immunopathology (Nieto-Torres *et al*., 2015).

### 19. Membrane glycoprotein (M protein)

Membrane glycoprotein has 9 helices, 8 helix-helix interacs, 28 beta turns and 11 gamma turns (Figure 3I). Membrane glycoprotein (Molecular Weight = 25145.16 Daltons) Leucine (15.77%) and Isoleucine (9.01%). Membrane glycoprotein presented pI 9.51, Instability index 39.14, Aliphatic index 120.86 and Grand average of hydropathicity 0.446 (Figure 2F; Table 3).

Biological process of M protein is viral life cycle (GO: 0019058); includes attachment and entry of the virus particle, decoding of genome information, translation of viral mRNA by host ribosomes, genome replication, and assembly and release of viral particles containing the genome. Out of the four estimated tunnels in Membrane glycoprotein, the two high throughput tunnels are tunnel-1 (green) is with bottleneck radius 1.7 Å, length 3.9 Å, distance to surface 3.3 Å, curvature 1.2, throughput 0.87 and number of residues 17; tunnel-2 (red) is with bottleneck radius 1.5 Å, length 7.4 Å, distance to surface 6.4 Å, curvature 1.2, throughput 0.76 and number of residues 18; (Figure 3I).

M protein is a component of the viral envelope that plays a central role in virus morphogenesis and assembly via its interactions with other viral proteins. Protein family membership of M protein includes FM matrix/glycoprotein, coronavirus (IPR002574). This family consists of various coronavirus matrix proteins which are transmembrane glycoproteins (Armstrong *et al*., 1984).

The membrane (M) protein is the most abundant structural protein and defines the shape of the viral envelope. It is also regarded as the central organiser of coronavirus assembly, interacting with all other major coronaviral structural proteins. M proteins play a critical role in protein-protein interactions (as well as protein-RNA interactions) since virus-like particle (VLP) formation in many CoVs requires only the M and envelope (E) proteins for efficient virion assembly (Ujike and Taguch, 2015).

Interaction of spike (S) with M is necessary for retention of S in the ER-Golgi intermediate compartment (ERGIC)/Golgi complex and its incorporation into new virions, but dispensable for the assembly process. Binding of M to nucleocapsid (N) proteins stabilises the nucleocapsid (N protein-RNA complex), as well as the internal core of virions, and, ultimately, promotes completion of viral assembly. Together, M and E protein make up the viral envelope and their interaction is sufficient for the production and release of virus-like particles (VLPs) (Schoeman and Fielding, 2019).

### 20. ORF6 protein

ORF6 protein shows 3 helices, 1 helix-helix interact, 2 beta turns and 1 gamma turn (Figure 3 J). ORF6 protein (Molecular Weight = 7272.15 Daltons) is rich in Isoleucine (16.39%) and Leucine (13.11%). ORF6 with pI 4.60, Instability index 31.16, Aliphatic index 130.98 and Grand average of hydropathicity 0.233 (Figure 2G; Table 3).

The ORF6 protein belongs to the protein family Non-structural protein NS6, betacoronavirus (IPR022736). Proteins in this family are typically between 42 to 63 amino acids in length, highly conserved among SARS-related coronaviruses (Geng *et al*., 2005).

Tunnels analysis results have shown four tunnels in the structure of ORF6 protin. Two of the mostly throughput tunnels are tunnel-1(green) with bottleneck radius 1.6 Å, length 12.7 Å, distance to surface 8.7 Å, curvature 1.0, throughput 0.78 and number of residues 17; The tunnel-2 (red) with bottleneck radius 1.2 Å, length 7.6 Å, distance to surface 6.6 Å, curvature 1.1, throughput 0.75 and number of residues 12 (Figure 3J).

### 21. ORF7a protein

Protein 7a (X4 like protein) is a non-structural protein which is dispensable for virus replication in cell culture. Structurally it consists of 1 sheet, 2 beta hairpins, 3 strands 5 helices, 4 helix-helix interacs, 8 beta turns, 1 gamma turn (Figure 3K).ORF7a protein (Molecular Weight = 13743.47 Daltons) is rich in Leucine (12.40%) Threonine (8.26 %) and Phenylalanine (8.26%). ORF7a protein has shown pI 8.23, Instability index 48.66, Aliphatic index 100.74 and Grand average of hydropathicity 0.318 (Figure 2H; Table 3).

Protein 7a (SARS coronavirus X4 like protein) (Pfam: PF08779 SARS_X4) is a unique type I transmembrane protein (Nelson *et al*., 2005). It has been suggested that it has binding activity to integrin I domains (Hänel *et al*., 2006). It contains a motif which has been demonstrated to mediate COPII dependent transport out of the endoplasmic reticulum, and the protein is targeted to the Golgi apparatus (InterPro IPR01488) (Pekosz *et al*., 2006).

Tunnels analysis results have estimated three tunnels in ORF7a protein. The best tunnel i.e.tunnel-1(blue) with bottleneck radius 2.6 Å, length 1.8 Å, distance to surface 1.8 Å, curvature 1.0, throughput 0.94 and number of residues 16; The tunnel-2 (green) with bottleneck radius 1.8 Å, length 6.9 Å, distance to surface 6.0 Å, curvature 1.2, throughput 0.80 and number of residues 17, The tunnel-3 (red) with bottleneck radius 1.3 Å, length 9.1 Å, distance to surface 7.2 Å, curvature 1.3, throughput 0.74 and number of residues 18 (Figure 3K).

### 22. ORF 7b protein

Structure of ORF7b has presented 2 helices, 1 helix-helix interact,1 beta turn and 1 gamma turn (Figure 3L). ORF7b protein (Molecular Weight = 5179.98 Daltons) is rich in Leucine (25.58%) and Phenylealanine (13.95%) having pI 4.17, Instability index 50.96, Aliphatic index 156.51 and Grand average of hydropathicity 1.449 (Figure 2I; Table 3).

ORF7b has Protein family membership Non-structural protein 7b, SARS-like (IPR021532) (also known as accessory protein 7b, NS7B, ORF7b, and 7b) from human SARS coronavirus (SARS-CoV) and similar betacoronaviruses (Pekosz *et al*., 2006). It consists of an N-terminal, a C-terminal and a transmembrane domain, the latter is essential to retain the protein in the Golgi compartment (Schaecher *et al*., 2007, 2008). Despite it being named as “non-structural”, it has been reported to be a structural component of SARS-CoV virions and an integral membrane protein (Schaecher *et al*., 2007).

The estimated high throughput tunnel-1 (blue) in ORF7b protein is with bottleneck radius 1.2 Å, length 3.3 Å, distance to surface 3.3 Å, curvature 1.0, throughput 0.89 and number of residues 4 (Figure 3L).

### 23. ORF8 protein

ORF8 protein has 3 sheets, 1 beta bulge, 10 strands, 15 beta turns and 1 gamma turn (Figure 3M). ORF8 protein (Molecular Weight = 13830.33 Daltons) is rich in Valine (9.92%) Leucine (8.26%) and Isoleucine (8.26%). And presented pI 5.42, Instability index 45.79, Aliphatic index 97.36 and Grand average of hydropathicity 0.219 (Figure 2J; Table 3).

ORF8 protein belongs to the family Non-structural protein NS8, betacoronavirus (IPR022722).This family of proteins includes the accessory proteins encoded by the ORF8 in coronaviruses, also known as accessory protein 8, or non-structural protein 8 (NS8). This is distinct from NSP8, which in encoded on the replicase polyprotein. This protein has two conserved sequence motifs: EDPCP and INCQ. It may modulate viral pathogenicity or replication in favour of human adaptation. ORF8 was suggested as one of the relevant genes in the study of human adaptation of the virus (Keng *et al*., 2006; Law *et al*., 2006).

The estimated high throughput tunnel-1 (blue) in ORF 8 protein is with bottleneck radius 1.9 Å, length 1.0 Å, distance to surface 1.0 Å, curvature 1.0, throughput 0.93 and number of residues 9 (Figure 3M).

### 24. Nucleocapsid phosphoprotein

Nucleocapsid phosphoprotein has demonstrated 1 sheet, 1 beta hairpin, 2 strands, 30 helices, 27 helix-helix interacs, 31 beta turns, 11 gamma turns (Figure 3N). Nucleocapsid phosphoprotein (Molecular Weight = 45623.27 Daltons) is rich in Glycine (10.26%), Alanine (8.83%) and Serine (8.83%) along with pI 10.07, Instability index 55.09, Aliphatic index 52.53 and Grand average of hydropathicity −0.971 (Figure 2K; Table 3).

The Nucleocapsid SARS-COV-2 (IPR001218) is the member of protein family Nucleocapsid protein, coronavirus (IPR001218) and Nucleocapsid protein, betacoronavirus (IPR043505). Coronavirus (CoV) nucleocapsid (N) proteins have 3 highly conserved domains. The N-terminal domain (NTD) (N1b), the C-terminal domain (CTD) (N2b) and the N3 region. The N1b and N2b domains from SARS CoV, infectious bronchitis virus (IBV), human CoV 229E and mouse hepatic virus (MHV) display similar topological organisations. N proteins form dimers, which are asymmetrically arranged into octamers via their N2b domains. The protein is cellular component of viral nucleocapsid (GO: 0019013).

Tunnels analysis results have estimated three tunnels in Nucleocapsid phosphoprotein. The best tunnel i.e.tunnel-1(blue) with bottleneck radius 2.0 Å, length 19.6 Å, distance to surface 14.9 Å, curvature 1.3, throughput 0.70 and number of residues 40; The tunnel-2 (green) with bottleneck radius 1.8 Å, length 25.1 Å, distance to surface 15.7 Å, curvature 1.6, throughput 0.63 and number of residues 45, The tunnel-3 (red) with bottleneck radius 1.3 Å, length 20.0 Å, distance to surface 8.1 Å, curvature 2.5, throughput 0.58 and number of residues 34 (Figure 3N).

Domains N1b and N2b are linked by another domain N2a that contains an SR-rich region (rich in serine and arginine residues). A priming phosphorylation of specific serine residues by an as yet unknown kinase, triggers the subsequent phosphorylation by the host glycogen synthase kinase-3 (GSK-3) of several residues in the SR-rich region. This phosphorylation allows the N protein to associate with the RNA helicase DDX1 permitting template read-through, and enabling the transition from discontinuous transcription of sub genomic mRNAs (sgmRNAs) to continuous synthesis of longer sgmRNAs and genomic RNA (gRNA). Production of gRNA in the presence of N oligomers may promote the formation of ribonucleoprotein complexes, and the newly transcribed sgmRNA would guarantee sufficient synthesis of structural proteins (Wu et *al*., 2014; Cong *et al*., 2017, 2020).

It has been shown that N proteins interact with nonstructural protein 3 (NSP3) and thus are recruited to the replication-transcription complexes (RTCs). In MHV, the N1b and N2a domains mediate the binding to NSP3 in a gRNA-independent manner. At the RTCs, the N protein is required for the stimulation of gRNA replication and sgmRNA transcription. It remains unclear, however, how and why the N protein orchestrates viral RNA synthesis. The cytoplasmic N-terminal ubiquitin-like domain of NSP3 and the SR-rich region of the N2a domain of the N protein may be important for this interaction. The direct association of N protein with RTCs is a critical step for MHV infection (Cong *et al*., 2020). Sequence comparison of the N genes of five strains of the coronavirus mouse hepatitis virus suggests a three domain structure for the nucleocapsid protein (Parker and Masters, 1990).

### 25. ORF10 protein

Structure wise ORF10 has shown 1 sheet, 1 beta alpha beta unit, 2 strands, 1 helix, 2 beta turns (Figure 3O).ORF10 protein (Molecular Weight = 4449.01 Daltons) is rich in Asperagine (13.16%), Leucine (10.53%), and Phenylalanine (10.53%) as well as carries pI 7.93, Instability index 16.06, Aliphatic index 107.63 and Grand average of hydropathicity 0.637 (Figure 2L; Table 3). Protein family membership has not been predicted for ORF 10 protein. The estimated high throughput tunnel-1 (blue) in ORF 10 protein is with bottleneck radius 1.5 Å, length 1.0 Å, distance to surface 1.0 Å, curvature 1.0, throughput 0.89 and number of residues 6 (Figure 3O).

### Hetero-Oligomeric Complexes

Nsp7-nsp8 hexadecamer may possibly confer processivity to the polymerase, may be by binding to dsRNA or by producing primers utilized by the latter. Experimental evidence for SARS-CoV that nsp7 and nsp8 activate and confer processivity to the RNA-synthesizing activity of Polymerase (Subissi *et al*., 2014; Kirchdoerfer and Ward, 2019). Nsp10 plays a pivotal role in viral transcription by stimulating nsp14 3’-5’ exoribonuclease activity. Nsp10 plays a pivotal role in viral transcription by stimulating nsp16 2’-O-ribose methyltransferase activity. Spike protein S1 binds to human ACE2, initiating the infection. CoV attaches to the target cells with the help of spike protein–host cell protein interaction (angiotensin converting enzyme-2 [ACE-2] interaction in SARS-CoV (Li *et al*., 2003) and dipeptidyl peptidase-4 [DPP-4] in MERS-CoV (Mubarak *et al*., 2019). After the receptor recognition, the virus genome with its nucleocapsid is released into the cytoplasm of the host cells. The viral genome contains ORF1a and ORF1b genes, which produce two PPs that are pp1a and pp1b (te Velthuis *et al*., 2016) which help to take command over host ribosome for their own translation process(Stobart *et al*., 2013).

The Instability index value of SARS-COV-2 ranged between 16.06 (ORF10 protein) and 51.97 (NSP7), which classifies ORF10 protein as most stable and NSP7 as most unstable protein. The proteins namely ORF7a protein (48.66), ORF7b protein (50.96), NSP7 (51.97), ORF 8 protein (45.79) and Nucleocapsid phosphoprotein (55.09) are unstable as per the instability index. The rest of the proteins are showing stability as per the instability index (Table 3).Except Nucleocapsid phosphoprotein and ORF10 protein all other proteins of SARS-COV-2 are rich in Lucine (Figure 2).The aliphatic index of SARS-COV2 ranged between 61.80 (NSP10) and 156.51 (ORF 7b protein), which indicates most thermostability in ORF 7b protein (Table 3).The Grand average of hydropathicity (GRAVY) value of NSP4 (0.343), NSP6 (0.790), NSP7 (0.199), ORF3a protein (0.275), Envelpe protein (1.128), Membrane glycoprotein (0.446), ORF6 protein (0.233), ORF7a protein (0.318), ORF 7b protein (1.449), ORF 8 protein (0.219) and ORF10 protein (0.637) indicate that these proteins are hydrophobic in nature. The all other proteins are hydrophilic (Table 3).

Tunnels are access paths connecting the interior of molecular systems with the surrounding environment. The presence of tunnels in proteins influences their reactivity, as they determine the nature and intensity of the interaction that these proteins can take part in (Jaiswal *et al*., 2012). Tunnel analysis of the newly predicted structures of the present study have estimated the presence of multiple tunnels in NSP1, NSP3, NSP6, ORF3a protein, ORF6 protein, membrane glycoprotein, ORF7a protein and Nucleocapsid phosphoprotein. The presence of multiple tunnels in these proteins may take key role in a large number of transport pathways for small ligands influencing their reactivity. It has been experimentally demonstrated that the tunnels and their properties can define many important protein characteristics like substrate specificity, enantioselectivity, stability and activity (Brezovsky *et al.*, 2016). The detail of the structure verification report has been deposited to Modelarchive and will be available to download alongwith the structures (https://www.modelarchive.org/doi/10.xxxx/).

## CONCLUSION

The RNA genome of SARS-CoV-2 has 29.9 Kb nucleotides, encoding for 29 proteins, though one may not get expressed. Studying these different components of the virus, as well as how they interact with human cells is already yielding some clues, but much remains to be explored. The present study reported theoretical modeling of 15 proteins, *In-silico* sequence-based and structure-based functional characteration of full SARS-CoV-2 proteome based on the NCBI reference sequence NC_045512 (29903 bp ss-RNA). Presence of large number of tunnels in NSP1, NSP3, NSP6, ORF3a protein, ORF6 protein, membrane glycoprotein, ORF7a protein and Nucleocapsid phosphoprotein indicates their high reactivity. The theoretical structures along with statistical verification reports have been successfully deposited to Model Archive. The 15 theoretical structures would perhaps be useful for the scientific community for advanced computational analysis on interactions of each protein for detailed functional analysis of active sites towards structure based drug designing or to study potential vaccines, if at all, towards preventing epidemics and pandemics in absence of complete experimental structure.

## ACKNOWLEDGEMENTS

The authors are grateful to DBT-Govt. of India for supporting Bioinformatics Laboratory (under DBT-Star College scheme) at Post Graduate Department of Zoology, Darrang College, Tepur, Assam. The authors are thankful to the Principal, Darrang College (Gauhati University), Tezpur (Assam) India and Head of the Post Graduate Department of Zoology, Darrang College and the University of Science and Technology, Meghalaya, India for supporting the research laboratory facility.

## REFERENCES

Altschul, S.F., Madden, T.L., Schaffer, A.A., Zhang, J., Zhang, Z., Miller, W. and Lipman, D.J. 1997. Gapped BLAST and PSI-BLAST: a new generation of protein database search programs. Nuc. Ac. Res., 25(17): 3389–3402.

Armstrong J, Niemann H, Smeekens S, Rottier P, Warren G. 1984. Sequence and topology of a model intracellular membrane protein, E1 glycoprotein, from a coronavirus. Nature 308:751-2, View article PMID: 6325918.

Brezovsky J., Babkova P., Degtjarik O., Fortova A., Gora A., Iermak I., Rezacova P., Dvorak P., Smatanova I.K., Prokop Z. et al. .. Engineering a de Novo Transport Tunnel. ACS Catal. 2016; 6:7597–7610.

Campanacci V, Egloff MP, Longhi S, Ferron F, Rancurel C, Salomoni A, Durousseau C, Tocque F, Bremond N, Dobbe JC, Snijder EJ, Canard B, Cambillau C. 2003. Structural genomics of the SARS coronavirus: cloning, expression, crystallization and preliminary crystallographic study of the Nsp9 protein. Acta Crystallogr D Biol Crystallogr; 59:1628-1631.:PUBMED:12925794 EPMC:12925794.

Chang CK, Jeyachandran S, Hu NJ, Liu CL, Lin SY, Wang YS, et al. 2016. Structure-based virtual screening and experimental validation of the discovery of inhibitors targeted towards the human coronavirus nucleocapsid protein. Mol Biosyst; 12:59–66.

Chen Y, Su C, Ke M, Jin X, Xu L, Zhang Z, Wu A, Sun Y, Yang Z, Tien P, Ahola T, Liang Y, Liu X, Guo D. 2011. Biochemical and structural insights into the mechanisms of SARS coronavirus RNA ribose 2’-O-methylation by nsp16/nsp10 protein complex. PLoS Pathog. 7:e1002294.: PUBMED:22022266 EPMC:22022266.

Cheng W, Chen S, Li R, Chen Y, Wang M, Guo D. 2015. Severe acute respiratory syndrome coronavirus protein 6 mediates ubiquitin-dependent proteosomal degradation of N-Myc (and STAT) interactor. Virol Sin 30, 153-61; PMID: 25907116.

Chiu, S.S., Chan, K.H., Chu, K.W., Kwan, S.W., Guan, Y., Poon, L.L., and Peiris, J.S. 2005. Human coronavirus NL63 infection and other coronavirus infections in children hospitalized with acute respiratory disease in Hong Kong, China. Clin. Infect. Dis. 40, 1721–1729.

Cong Y, Kriegenburg F, de Haan CAM, Reggiori F. 2017. Coronavirus nucleocapsid proteins assemble constitutively in high molecular oligomers. Sci Rep 7, 5740, PMID: 28720894.

Cong Y, Ulasli M, Schepers H, Mauthe M, V’kovski P, Kriegenburg F, Thiel V, de Haan CAM, Reggiori F. 2020. Nucleocapsid Protein Recruitment to Replication-Transcription Complexes Plays a Crucial Role in Coronaviral Life Cycle. J. Virol. 94, PMID: 31776274.

Coronaviridae Study Group of the International Committee on Taxonomy of Viruses. 2020. The species Severe acute respiratory syndrome-related coronavirus: classifying 2019-nCoV and naming it SARS-CoV-2. Nat. Microbiol. 5, 536–544.

de Groot, R. J. et al. in Virus Taxonomy, Ninth Report of the International Committee on Taxonomy of Viruses (eds King, A. M. Q. et al.) 806–828 (Elsevier Academic Press, 2012).

DeLano, W. L. 2002. Pymol: An open-source molecular graphics tool. CCP4 Newsletter On Protein Crystallography, 40, 82–92.

Egloff MP, Ferron F, Campanacci V, Longhi S, Rancurel C, Dutartre H, Snijder EJ, Gorbalenya AE, Cambillau C, Canard B. 2004. The severe acute respiratory syndrome-coronavirus replicative protein nsp9 is a single-stranded RNA-binding subunit unique in the RNA virus world., Proc Natl Acad Sci U S A. 101:3792-3796.PUBMED:15007178 EPMC:15007178.

Eswar, N., Webb, B., Marti-Renom, M.A., Madhusudhan, M.S., Eramian, D., Shen, M.Y., Pieper, U. and Sali, A. 2006. Comparative Protein Structure Modelling Using MODELLER. Ed: Coligan JE, Dunn BM, Speicher DW, Wingfield PT. Current Protocols in Protein Science Unit. 2.9 1–31.

Finn R.D., Clements J., Eddy S.R. 2011. HMMER Web Server: Interactive Sequence Similarity Searching. R. D. Finn, J. Clements, S. R. Eddy. Nucleic Acids Research, 39:W29–37.

Fiser, A., Do, R.K. and Sali, A. 2000. Modeling of loops in protein structures. Protein Science, 9: 1753–1773.

Gasteiger, E., Hoogland, C., Gattiker, A., Duvaud, S., Wilkins, M.R., Appel, R.D. and Bairoch, A. 2005. Protein Identification and Analysis Tools on the ExPASy Server; (In) John M. Walker (ed): The Proteomics Protocols Handbook, Humana Press pp. 571–607.

Geng H, Liu YM, Chan WS, Lo AW, Au DM, Waye MM, Ho YY. 2005. The putative protein 6 of the severe acute respiratory syndrome-associated coronavirus: expression and functional characterization. FEBS Lett. 579, 6763-8. PMID: 16310783

Haas J., Gumienny R., Barbato A., Ackermann F., Tauriello G., Bertoni M., Studer G., Smolinski A., Schwede T. 2019, Introducing “best single template” models as reference baseline for the Continuous Automated Model Evaluation (CAMEO). Proteins. 87, 1378-1387. [DOI: 10.1002/prot.25815].

Hänel K, Stangler T, Stoldt M, Willbold D. 2006. “Solution structure of the X4 protein coded by the SARS related coronavirus reveals an immunoglobulin like fold and suggests a binding activity to integrin I domains”. J. Biomed. Sci. 13 (3): 281–93. PMID 16328780. doi: 10.1007/s11373-005-9043-9.

Huang, C. et al. 2020. Clinical features of patients infected with 2019 novel coronavirus in Wuhan, China. Lancet 395, 497–506.

Jaiswal, D., Vařeková, R. S., Ionescu, C. M., Sehnal, D., & Koča, J. (2012). Searching for tunnels of proteins – comparison of approaches and available software tools. Journal of Cheminformatics, 4(Suppl 1), P60. https://doi.org/10.1186/1758-2946-4-S1-P60

Jean, A., Quach, C., Yung, A., and Semret, M. 2013. Severity and outcome associated with human coronavirus OC43 infections among children. Pediatr. Infect. Dis. J. 32, 325–329.

Jiang, S., Shi, Z., Shu, Y., Song, J., Gao, G.F., Tan, W., and Guo, D. 2020. A distinct name is needed for the new coronavirus. Lancet 395, 949.

Joseph JS, Saikatendu KS, Subramanian V, Neuman BW, Brooun A, Griffith M, Moy K, Yadav MK, Velasquez J, Buchmeier MJ, Stevens RC, Kuhn P. 2006. Crystal structure of nonstructural protein 10 from the severe acute respiratory syndrome coronavirus reveals a novel fold with two zinc-binding motifs. J Virol. 80:7894-7901.: PUBMED:16873246 EPMC:16873246.

Keng CT, Choi YW, Welkers MR, Chan DZ, Shen S, Gee Lim S, Hong W Tan. 2006. The human severe acute respiratory syndrome coronavirus (SARS-CoV) 8b protein is distinct from its counterpart in animal SARS-CoV and down-regulates the expression of the envelope protein in infected cells. YJ. Virology 354, 132-42, PMID: 16876844.

Kirchdoerfer, R.N. and Ward, A.B. 2019. Structure of the SARS-CoV nsp12 polymerase bound to nsp7 and nsp8 co-factors. Nature Communications, 10(1), pp.1–9.

Lai, M.M.C., Perlman, S., and Anderson, L.J. 2007. Coronaviridae. In Fields Virology, D.M. Knipe and P.M. Howley, eds. (Lippincott Williams & Wilkins), pp. 1305–1335.

Laskowski, R.A., Watson, J.D. and Thornton, J.M. 2005. ProFunc: a server for predicting protein function from 3D structure. Nucleic Acids Res., 33: W89–W93.

Law PY, Liu YM, Geng H, Kwan KH, Waye MM, Ho YY. 2006. Expression and functional characterization of the putative protein 8b of the severe acute respiratory syndrome-associated coronavirus. FEBS Lett. 580, 3643-8, PMID: 16753150

Li W, Moore MJ, Vasilieva N, Sui J, Wong SK, Berne MA, et al. 2003. Angiotensin-converting enzyme 2 is a functional receptor for the SARS coronavirus. Nature 426:450–4.

Lili Chen, Chunshan Gui, Xiaomin Luo, Qingang Yang, Stephan Günther, Elke Scandella, Christian Drosten, Dongl u Bai, Xichang He, Burkhard Ludewig, Jing Chen, Haibin Luo, Yiming Yang, Yifu Yang, Jianping Zou, Vol ker Thiel, Kaixian Chen, Jianhua Shen, Xu Shen, Hualiang Jiang. 2005. Journal of Virology May 2005, 79 (11) 7095–7103; DOI: 10.1128/JVI.79.11.7095-7103.

Lu W, Zheng BJ, Xu K, Schwarz W, Du L, Wong CK, Chen J, Duan S, Deubel V, Sun B. 2006. Severe acute respiratory syndrome-associated coronavirus 3a protein forms an ion channel and modulates virus release. Proc. Natl. Acad. Sci. U.S.A. 103, 12540-5, View article PMID: 16894145

Lu, G., and Liu, D. 2012. SARS-like virus in the Middle East: a truly bat-related coronavirus causing human diseases. Protein Cell 3, 803–805.

Lu, G., Wang, Q., and Gao, G.F. 2015. Bat-to-human: spike features determining ‘host jump’ of coronaviruses SARS-CoV, MERS-CoV, and beyond. Trends Microbiol. 23, 468–478.

Madan V, Garcia Mde J, Sanz MA, Carrasco L.; 2005. Viroporin activity of murine hepatitis virus E protein. FEBS Lett. 579, 3607-12, PMID: 15963987.

Manolaridis I, Wojdyla JA, Panjikar S, Snijder EJ, Gorbalenya AE, Berglind H, Nordlund P, Coutard B, Tucker PA. 2009. Structure of the C-terminal domain of nsp4 from feline coronavirus. Acta Crystallogr D Biol Crystallogr.; 65:839-846.: PUBMED:19622868 EPMC:19622868.

Minakshi R, Padhan K, Rani M, Khan N, Ahmad F, Jameel S. 2009. The SARS Coronavirus 3a protein causes endoplasmic reticulum stress and induces ligand-independent downregulation of the type 1 interferon receptor. PLoS ONE 4, e8342. View article PMID: 20020050

Mubarak A, Alturaiki W, Hemida MG. 2019. Middle East Respiratory Syndrome Coronavirus (MERS-CoV): Infection, Immunological Response, and Vaccine Development. J Immunol Res.;2019:1–11.

Nelson CA, Pekosz A, Lee CA, Diamond MS, Fremont DH. 2005. “Structure and intracellular targeting of the SARS-coronavirus Orf7a accessory protein.”. Structure. 13 (1): 75–85. PMID 15642263. doi: 10.1016/j.str.2004.10.010.

Nieto-Torres JL, Verdia-Baguena C, Jimenez-Guardeno JM, Regla-Nava JA, Castano-Rodriguez C, Fernandez-Delgado R, Torres J, Aguilella VM, Enjuanes L. 2015. Severe acute respiratory syndrome coronavirus E protein transports calcium ions and activates the NLRP3 inflammasome. Virology 485, 330-9, PMID:26331680.

Parker, M. M. & Masters, P. S. 1990. Sequence comparison of the N genes of five strains of the coronavirus mouse hepatitis virus suggests a three domain structure for the nucleocapsid protein. Virology 179, 463-468.[CrossRef]

Paules CI, Marston HD, Fauci AS. Coronavirus infections-More than just the common cold. JAMA 2020;323:707.

Pearson, W.R. 1991. Searching protein sequence libraries: comparison of the sensitivity and selectivity of the Smith-Waterman and FASTA algorithms. Genomics, 11(3): 635–50.

Pekosz A, Schaecher SR, Diamond MS, Fremont DH, Sims AC, Baric RS. 2006. “Structure, expression, and intracellular localization of the SARS-CoV accessory proteins 7a and 7b.”. Adv Exp Med Biol. 581: 115–20. PMID 17037516. doi: 10.1007/978-0-387-33012-9_20.

Ricagno S, Egloff MP, Ulferts R, et al. Crystal structure and mechanistic determinants of SARS coronavirus nonstructural protein 15 define an endoribonuclease family. Proceedings of the National Academy of Sciences of the United States of America. 2006 Aug;103(32):11892–11897. DOI: 10.1073/pnas.0601708103.

Schaecher SR, Diamond MS, Pekosz A. 2008. The transmembrane domain of the severe acute respiratory syndrome coronavirus ORF7b protein is necessary and sufficient for its retention in the Golgi complex. J. Virol. 82, 9477-91. PMID: 18632859

Schaecher SR, Mackenzie JM, Pekosz A. 2007. The ORF7b protein of severe acute respiratory syndrome coronavirus (SARS-CoV) is expressed in virus-infected cells and incorporated into SARS-CoV particles. J. Virol. 81, 718-31, PMID: 17079322

Schoeman D, Fielding BC. 2019. Coronavirus envelope protein: current knowledge. Virol. J. 16, 69, PMID: 31133031.

Sehnal, D., Svobodová Vařeková, R., Berka, K. et al. 2013. MOLE 2.0: advanced approach for analysis of biomacromolecular channels. J Cheminform 5, 39 https://doi.org/10.1186/1758-2946-5-39.

Shen S, Lin PS, Chao YC, Zhang A, Yang X, Lim SG, Hong W, Tan YJ. 2005. The severe acute respiratory syndrome coronavirus 3a is a novel structural protein. Biochem. Biophys. Res. Commun. 330, 286-92, View article PMID: 15781262

Siddell, S. G. et al. 2008. Additional changes to taxonomy ratified in a special vote by the International Committee on Taxonomy of Viruses. Arch. Virol. 164, 943–946 (2019).

Stourac, J., Vavra, O., Kokkonen, P., Filipovic, J., Pinto, G., Brezovsky, J., Damborsky, J., Bednar, D., 2019: Caver Web 1.0: Identification of Tunnels and Channels in Proteins and Analysis of Ligand Transport. Nucleic Acids Research 47: W414–W422.

Stobart CC, Sexton NR, Munjal H, Lu X, Molland KL, Tomar S, et al. 2013. Chimeric exchange of coronavirus nsp5 proteases (3CLpro) identifies common and divergent regulatory determinants of protease activity. J Virol. 87:12611–8.

Subissi, L., Posthuma, C.C., Collet, A., Zevenhoven-Dobbe, J.C., Gorbalenya, A.E., Decroly, E., Snijder, E.J., Canard, B. and Imbert, I., 2014. One severe acute respiratory syndrome coronavirus protein complex integrates processive RNA polymerase and exonuclease activities. Proceedings of the National Academy of Sciences, 111(37), pp.E3900–E3909.

Surya W, Li Y, Verdia-Baguena C, Aguilella VM Torres. 2015. MERS coronavirus envelope protein has a single transmembrane domain that forms pentameric ion channels. J. Virus Res. 201, 61-66. View article PMID: 25733052.

te Velthuis AJ, van den Worm SH, Snijder EJ. 2012. The SARS-coronavirus nsp7+nsp8 complex is a unique multimeric RNA polymerase capable of both de novo initiation and primer extension. Nucleic Acids Res 40:1737–47.

Ujike M, Taguchi F. 2015. Incorporation of spike and membrane glycoproteins into coronavirus virions. Viruses 7, 1700-25, PMID: 25855243.

Webb B., Sali A.. 2016. Comparative Protein Structure Modeling Using Modeller. Current Protocols in Bioinformatics 54, John Wiley & Sons, Inc., 5.6.1-5.6.37.

Woo PC, Lau SK, Lam CS, et al. 2012. Discovery of seven novel Mammalian and avian coronaviruses in the genus deltacoronavirus supports bat coronaviruses as the gene source of alphacoronavirus and betacoronavirus and avian coronaviruses as the gene source of gammacoronavirus and deltacoronavirus. J Virol.;86(7):3995D4008. doi: 10.1128/JVI.06540-11

Wu CH, Chen PJ, Yeh SH. 2014. Nucleocapsid phosphorylation and RNA helicase DDX1 recruitment enables coronavirus transition from discontinuous to continuous transcription. Cell Host Microbe 16, 462-72,. PMID: 25299332

Wu, F., Zhao, S., Yu, B., Chen, Y.-M., Wang, W., Song, Z.-G., Hu, Y., Tao, Z.-W., Tian, J.-H., Pei, Y.-Y., et al. 2020. A new coronavirus associated with human respiratory disease in China. Nature 579, 265–269.

Yang H, Bartlam M, Rao Z. Drug design targeting the main protease, the Achilles’ heel of coronaviruses. Curr Pharm Des 2006;12:4573–90.

Yibei Xiao, Qingjun Ma, Tobias Restle, Weifeng Shang, Dmitri I. Svergun, Rajesh Ponnusamy, Georg Sczakiel, Rolf Hilgenfeld J Virol. 2012 Apr; 86(8): 4444–4454. doi: 10.1128/JVI.06635-11.

Yu CJ, Chen YC, Hsiao CH, Kuo TC, Chang SC, Lu CY, Wei WC, Lee CH, Huang LM, Chang MF, Ho HN, Lee FJ. 2004. Identification of a novel protein 3a from severe acute respiratory syndrome coronavirus. FEBS Lett. 565:111-116. PUBMED:15135062 EPMC:15135062.

Zhai Y, Sun F, Li X, Pang H, Xu X, Bartlam M, Rao Z. 2005. Insights into SARS-CoV transcription and replication from the structure of the nsp7-nsp8 hexadecamer. Nat Struct Mol Biol. 12:980-986.: PUBMED:16228002 EPMC:16228002.

Zhou, P., Yang, X.-L., Wang, X.-G., Hu, B., Zhang, L., Zhang, W., Si, H.-R., Zhu, Y., Li, B., Huang, C.-L., et al. 2020). A pneumonia outbreak associated with a new coronavirus of probable bat origin. Nature 579, 270–273.

Zhu, N., Zhang, D., Wang, W., Li, X., Yang, B., Song, J., Zhao, X., Huang, B., Shi, W., Lu, R., et al. 2020. A novel coronavirus from patients with pneumonia in China, 2019. N. Engl. J. Med. 382, 727–733.

Ziebuhr, J. et al. Proposal 2017.013S. A.v1. Reorganization of the family Coronaviridae into two families, Coronaviridae (including the current subfamily Coronavirinae and the new subfamily Letovirinae) and the new family Tobaniviridae (accommodating the current subfamily Torovirinae and three other subfamilies), revision of the genus rank structure and introduction of a new subgenus rank. (ICTV, 2017); https://ictv.global/proposal/2017.Nidovirales/.

Ziebuhr, J. et al. Proposal 2019.021S.Ac.v1. Create ten new species and a new genus in the subfamily Orthocoronavirinae of the family Coronaviridae and five new species and a new genus in the subfamily Serpentovirinae of the family.

