## Supplementary material for "*In silico* Proteome analysis of Severe acute respiratory syndrome coronavirus 2 (SARS-CoV-2)": Table 4 and Figure 4: Table 4 ProFunc results.docx

**Annexure-1**

**Table 4.** Predicted functions of SARS-CoV-2 proteome with respective ProFunc score (shown within parenthesis)

| **PROTEIN NAME** | **Summary of predicted function** | | | |
| --- | --- | --- | --- | --- |
|  | **Protein name terms** | **Gene Ontology (GO) terms** | | |
|  |  | **Cellular component** | **Biological process** | **Biochemical function** |
| NSP1  ([YP_009725297.1](https://www.ncbi.nlm.nih.gov/protein/1802476805)) | **bound** (1.75) **human** (1.00) **streptococcus** (1.00) **nucleoside** (0.77) **nmr** (0.70) **nonstructural** (0.70) **nsp1** (0.70) **sars coronavirus** (0.70) | **cytoplasm** (0.50) **cytosol** (0.50) **cell** (0.50) **cell part** (0.50) | **metabolic process** (1.75) **catabolic process** (1.26) **primary metabolic process** (1.26) **cellular process** (1.00) | **catalytic activity** (2.21) **binding** (1.26) **transferase activity** (1.25) **metal ion binding** (0.89) |
| NSP2  ([YP_009725298.1](https://www.ncbi.nlm.nih.gov/protein/1802476806)) | **domain** (1.06) **dehydrogenase** (1.00) **binding** (0.89) **variant** (0.88) **bound** (0.84) **zp-c domain** (0.56) **bmrr** (0.55) **myelin** (0.54) | **cytoplasm** (2.57) **cell** (2.57) **cell part** (2.57) **intracellular** (2.57) | **metabolic process** (3.52) **primary metabolic process** (2.40) **cellular process** (1.69) **cellular metabolic process** (1.69) | **binding** (4.05) **catalytic activity** (3.12) **metal ion binding** (2.06) **ion binding** (2.06) |
| NSP3  ([YP_009725299.1](https://www.ncbi.nlm.nih.gov/protein/1802476807)) | **ubiquitin** (3.09) **papain-like protease** (2.16) **domain** (1.95) **sars-cov** (1.32) **enzyme** (1.20) **coronavirus** (1.19) **virus** (1.10) **ubiquitin carboxyl-terminal hydrolase** (1.05) | **lysosome** (0.70) **cytosol** (0.70) **cell** (0.70) **cell part** (0.70) | **metabolic process** (1.22) **proteolysis** (0.99) **catabolic process** (0.99) **macromolecule catabolic process** (0.99) | **metal ion binding** (0.83) **binding** (0.83) **ion binding** (0.83) **cation binding** (0.83) |
| NSP4  ([YP_009725300.1](https://www.ncbi.nlm.nih.gov/protein/1802476808)) | **pneumoniae** (1.06) **domain** (0.86) **chemokine receptor** (0.83) **penicillin-binding** (0.55) **pbp-2b** (0.55) **streptococcus pneumoniae strain** (0.55) **cytochrome** (0.50) **ba3** (0.50) | **cell** (3.46) **cell part** (3.46) **membrane** (2.21) **membrane part** (2.21) | **metabolic process** (4.66) **cellular process** (4.25) **cellular metabolic process** (3.86) **primary metabolic process** (2.76) | **binding** (3.91) **catalytic activity** (2.74) **metal ion binding** (2.56) **ion binding** (2.56) |
| 3C-like proteinase  ([YP_009725301.1](https://www.ncbi.nlm.nih.gov/protein/1802476809)) | **protease** (9.00) **main** (3.26) **domain** (2.90) **main protease** (2.72) **coronavirus** (1.84) **virus** (1.80) **proteinase** (1.57) **staphylococcus aureus** (1.51) | **plasma membrane** (0.97) **membrane** (0.97) **integral to membrane** (0.97) **outer membrane\-bounded periplasmic space** (0.97) | **proteolysis** (3.16) **metabolic process** (3.16) **catabolic process** (3.16) **macromolecule catabolic process** (3.16) | **peptidase activity** (3.16) **serine\-type peptidase activity** (3.16) **hydrolase activity** (3.16) **catalytic activity** (3.16) |
| NSP6  ([YP_009725302.1](https://www.ncbi.nlm.nih.gov/protein/1802476810)) | **domain** (1.76) **human** (1.47**n-terminal domain** (0.83) **human class** (0.77) **metabotropic glutamate receptor** (0.77) | **cell** (3.97) **cell part** (3.97) **intracellular** (2.36) **intracellular part** (2.36) | **cellular process** (2.88) **metabolic process** (2.86) **primary metabolic process** (2.17) **cellular metabolic** | **binding** (3.71) **catalytic activity** (3.02) **protein binding** (2.21) **hydrolase activity** (1.85) |
| NSP7  ([YP_009725303.1](https://www.ncbi.nlm.nih.gov/protein/1802476811)) | **virus** (2.35) **polymerase** (2.21) **rna** (1.88) **virus rna** (1.20) **rna polymerase** (1.17) **rna-dependent rna polymerase** (1.00) **human** (0.99) **sars-cov super** (0.90) | **virion** (0.50) | **glutamate catabolic process via L\-citramalate** (0.50) **anaerobic glutamate catabolic process** (0.50) **cellular process** (0.50) **cellular metabolic process** (0.50) | **binding** (0.78) **catalytic activity** (0.78) **protein binding** (0.50) **isomerase activity** (0.50) |
| NSP8  ([YP_009725304.1](https://www.ncbi.nlm.nih.gov/protein/1802476812)) | **virus** (2.35) **polymerase** (2.21) **rna** (1.88) **virus rna** (1.20) **rna polymerase** (1.17) **rna-dependent rna polymerase** (1.00) **human** (0.99) **sars-cov super** (0.90) | **virion** (0.50) | **glutamate catabolic process via L\-citramalate** (0.50) **anaerobic glutamate catabolic process** (0.50) **cellular process** (0.50) **cellular metabolic process** (0.50) | **binding** (0.78) **catalytic activity** (0.78) **protein binding** (0.50) **isomerase activity** (0.50) |
| NSP9  ([YP_009725305.1](https://www.ncbi.nlm.nih.gov/protein/1802476813)) | **nsp9** (1.60) **domain** (1.20) **coronavirus** (1.10) **sars-coronavirus** (0.90) **sars coronavirus nsp9** (0.90) **sars-cov nsp9 g104e** (0.90) **receptor** (0.81) **bound** (0.81) | **cytoplasm** (1.33) **cell** (1.33) **cell part** (1.33) **intracellular** (1.33) | **metabolic process** (1.70) **catabolic process** (1.70) **primary metabolic process** (1.70) **cellular process** (1.34) | **hydrolase activity** (1.76) **catalytic activity** (1.76) **hydrolase activity\, acting on ester bonds** (1.30) **binding** (1.22) |
| NSP10  ([YP_009725306.1](https://www.ncbi.nlm.nih.gov/protein/1802476814)) | **sars coronavirus** (2.11) **methyltransferase** (2.01) **nsp10** (1.80) **dehydrogenase** (1.39) **nsp16 nsp10 sars coronavirus** (0.90) **nsp10/nsp16** (0.90) **reovirus core** (0.90) **chikungunya virus nsp2 protease** (0.90) | **cell** (1.01) **cell part** (1.01) **intracellular** (1.01) **intracellular part** (1.01) | **metabolic process** (4.63) **cellular process** (3.74) **cellular metabolic process** (3.74) **methylation** (2.48) | **catalytic activity** (4.63) **binding** (3.73) **methyltransferase activity** (2.48) **transferase activity** (2.48) |
| RNA-dependent RNA polymerase  ([YP_009725307.1](https://www.ncbi.nlm.nih.gov/protein/1802476815)) | **human** (1.64) **domain** (1.00) **phospholipase** (1.00) **aps kinase** (0.52) **penicillium chrysogenum ternary** (0.52) **adp** (0.52) **repeats** (0.50) **chicken brain alpha spectrin** (0.50) | **cell** (1.70) **cell part** (1.70) **intracellular** (1.70) **intracellular part** (1.70) | **cellular process** (2.20) **metabolic process** (2.13) **cellular metabolic process** (1.70) **primary metabolic process** (1.70) | **catalytic activity** (2.63) **binding** (2.62) **metal ion binding** (1.41) **ion binding** (1.41) |
| Helicase  ([YP_009725308.1](https://www.ncbi.nlm.nih.gov/protein/1802476816)) | **helicase** (5.47) **human** (1.89) **mitochondrial** (1.38) **f1-atpase** (1.26) | **cell** (4.53) **cell part** (4.53) **intracellular** (4.53) **intracellular part** (4.53) | **cellular process** (5.07) **cellular metabolic process** (4.53) **nucleobase\, nucleoside\, nucleotide and nucleic acid metabolic process** (4.53) **metabolic process** (4.53) | **nucleotide binding** (5.07) **ATP binding** (5.07) **binding** (5.07) **purine nucleotide binding** (5.07) \| |
| 3'-to-5' exonuclease  ([YP_009725309.1](https://www.ncbi.nlm.nih.gov/protein/1802476817)) | **dna** (3.37) **polymerase** (3.37) **dna polymerase** (2.69) **exonuclease** (1.78) **rnase** (1.70) **type dna polymerase** (1.40) **angstrom** (1.40) **coli** (1.29) | **cytoplasm** (3.00) **cell** (3.00) **cell part** (3.00) **intracellular** (3.00) | **cellular process** (7.14) **cellular metabolic process** (7.14) **cellular macromolecule metabolic process** (7.14) **metabolic process** (7.14) | **catalytic activity** (7.14) **binding** (6.43) **nuclease activity** (6.32) **hydrolase activity** (6.32) |
| endoRNAse/ NSP15  ([YP_009725310.1](https://www.ncbi.nlm.nih.gov/protein/1802476818)) | **bound** (0.97) **sars** (0.90) **mutation mhv coronavirus non-structural** (0.90) **nsp15 f307l** (0.90) **xendou splicing independent snorna** (0.90) **splicing independent snorna processing** (0.90) **independent snorna processing endoribonuclease** (0.90) | **cell** (3.16) **cell part** (3.16) **intracellular** (2.86) **intracellular part** (2.86) | **cellular process** (3.17) **cellular metabolic process** (2.85) **metabolic process** (2.85) **primary metabolic process** (2.44) | **binding** (3.44) **catalytic activity** (2.40) **metal ion binding** (2.24) **ion binding** (2.24) |
| 2'-O-ribose methyltransferase  ([YP_009725311.1](https://www.ncbi.nlm.nih.gov/protein/1802476819)) | **sars coronavirus** (2.11) **methyltransferase** (2.01) **nsp10** (1.80) **dehydrogenase** (1.39) **nsp16 nsp10 sars coronavirus** (0.90) **nsp10/nsp16** (0.90) **reovirus core** (0.90) **chikungunya virus nsp2 protease** (0.90) | **cell** (1.01) **cell part** (1.01) **intracellular** (1.01) **intracellular part** (1.01) | **metabolic process** (4.63) **cellular process** (3.74) **cellular metabolic process** (3.74) **methylation** (2.48) | **catalytic activity** (4.63) **binding** (3.73) **methyltransferase activity** (2.48) **transferase activity** (2.48) |
| Surface glycoprotein  (QJF77858.1) | **coronavirus** (4.75) **domain** (4.33) **spike** (3.39) **receptor** (3.27) **receptor-binding domain** (2.59) **sars** (2.58) **human** (1.85) **sars coronavirus** (1.71) | **cytoplasm** (0.50) **cell** (0.50) **cell part** (0.50) **intracellular** (0.50) | **glycine biosynthetic process** (0.50) **one\-carbon compound metabolic process** (0.50) **cellular amino acid biosynthetic process** (0.50) **glycine biosynthetic process from serine** (0.50) | **glycine hydroxymethyltransferase activity** (0.50) **methyltransferase activity** (0.50) **transferase activity** (0.50) **pyridoxal phosphate binding** (0.50) |
| ORF3a protein  (QJF77859.1) | **dna** (1.78) **dna glycosylase** (1.01) **mycobacterium tuberculosis** (0.87) **bound** (0.87) **onconase double** (0.52) **p2y12 receptor** (0.50) **2mesadp** (0.50) **spontaneously- assembled amp stack** (0.50) | **cell** (1.81) **cell part** (1.81) **cytoplasm** (0.88) **intracellular** (0.88) | **metabolic process** (2.99) **cellular process** (2.48) **cellular metabolic process** (2.48) **primary metabolic process** (2.07) | **binding** (3.04) **catalytic activity** (2.76) **metal ion binding** (2.30) **ion binding** (2.30) |
| Envelope protein  (QJF77860.1) | **domain** (0.91) **human** (0.53) **cca-adding enzyme** (0.53) **pyruvate carboxylase** (0.50) **rhizobium etli** (0.50) **golgi complex-targeting signal coronavirus** (0.50) **complex-targeting signal coronavirus envelope** (0.50) **full-length human mitochondrial cca-adding** (0.50) | **intracellular** (0.95) **cell** (0.95) **cell part** (0.95) **intracellular part** (0.95) | **metabolic process** (0.95) **tRNA 3'\-terminal CCA addition** (0.53) **tRNA processing** (0.53) **tRNA 3'\-end processing** (0.53) | **binding** (1.42) **catalytic activity** (1.42) **nucleotide binding** (1.05) **ATP binding** (1.05) |
| Membrane glycoprotein  (QJF77861.1) | **peptide** (1.35) **coli** (1.35) **transporter** (1.27) **domain** (0.80) **xylella fastidiosa** (0.79) **oppa** (0.61) **viral** (0.51) **rna polymerase** (0.51) | **cell** (3.76) **cell part** (3.76) **membrane** (2.06) **integral to membrane** (1.67) | **establishment of localization** (2.10) **transport** (2.10) **localization** (2.10) **metabolic process** (2.01) | **binding** (3.58) **catalytic activity** (2.79) **protein binding** (2.43) **nucleotide binding** (2.08) |
| ORF6 protein  (QJF77862.1) | **domain** (1.74) **human** (1.36) **binding** (1.21) **binding domain** (1.00) **bound** (1.00) **synthase** (0.89) **nmr** (0.82) **region** (0.82) | **cell** (2.02) **cell part** (2.02) **cytoplasm** (1.64) **intracellular** (1.64) | **cellular process** (2.15) **cellular metabolic process** (2.15) **metabolic process** (2.15) **cellular biosynthetic process** (1.28) | **catalytic activity** (2.48) **binding** (2.03) **metal ion binding** (1.28) **ion binding** (1.28) |
| ORF7a protein  (QJF77863.1) | **methionine** (0.90) **human** (0.88) **sars-coronavirus orf7a accessory** (0.50) **human mterf4-nsun4** (0.50) **sars coronavirus orf coded** (0.50) **chimeric** (0.50) **5-ht1b-bril** (0.50) **ergotamine psi community** (0.50) | **cell** (2.79) **cell part** (2.79) **macromolecular complex** (1.62) **cytoplasm** (1.57) | **cellular process** (2.72) **metabolic process** (2.32) **macromolecule metabolic process** (1.93) **biopolymer metabolic process** (1.93) | **binding** (3.78) **catalytic activity** (2.43) **metal ion binding** (1.92) **ion binding** (1.92) |
| ORF7b protein  (QJF77864.1) | **domain** (1.21) **biosynthesis** (1.19) **human** (0.84) **phenazine biosynthesis** (0.76) **stearoyl-acyl carrier** (0.70) **desaturase** (0.70) **castor seeds** (0.70) **76a** (0.50) | **cell** (0.90) **cell part** (0.90) **protein farnesyltransferase complex** (0.50) **intracellular** (0.50) | **cellular process** (1.26) **metabolic process** (0.89) **biological regulation** (0.87) **protein farnesylation** (0.50) | **binding** (1.63) **metal ion binding** (1.26) **catalytic activity** (1.26) **ion binding** (1.26) |
| ORF8 protein  (QJF77865.1) | **domain** (1.98) **receptor** (1.33) **cell** (1.00) **cerevisiae** (0.86) **n-terminal** (0.80) **human** (0.78) **malt1** (0.53) **malt1 paracaspase p21 form** (0.50) | **cell** (2.70) **cell part** (2.70) **cytoplasm** (1.47) **intracellular** (1.47) | **cellular process** (1.95) **localization** (1.57) **metabolic process** (1.55) **establishment of localization** (1.23) | **binding** (1.85) **protein binding** (1.57) **catalytic activity** (1.44) **transferase activity** (0.90) |
| Nucleocapsid phosphoprotein  (QJF77866.1) | **domain** (3.83) **nucleocapsid** (2.44) **coronavirus nucleocapsid** (1.75) **dimerization domain** (1.74) **ubiquitin** (1.03) **human** (0.91) **oligomerization domain sars coronavirus** (0.90) **domain sars coronavirus nucleocapsid** (0.90 | **cytoplasm** (0.50) **cell** (0.50) **cell part** (0.50) **intracellular** (0.50) | **cellular process** (0.83) **cellular metabolic process** (0.83) **cellular biosynthetic process** (0.83) **metabolic process** (0.83) | **metal ion binding** (1.19) **binding** (1.19) **ion binding** (1.19) **cation binding** (1.19) |
| ORF10 protein  (QJF77867.1) | **domain** (2.03) **human** (1.24) **isomerase domain** (1.00) **domain human** (0.90) **angstrom** (0.80) **synthase** (0.50) **xylose isomerase domain** (0.50) **planctomyces limnophilus** (0.50) | **cell** (3.03) **cell part** (3.03) **intracellular** (2.61) **intracellular part** (2.61) | **metabolic process** (4.32) **cellular process** (3.61) **cellular metabolic process** (3.61) **primary metabolic process** (2.80) | **binding** (4.46) **catalytic activity** (3.96) **metal ion binding** (2.89) **ion binding** (2.89) |
