## Supplementary figures and images for "*In silico* Proteome analysis of Severe acute respiratory syndrome coronavirus 2 (SARS-CoV-2)"

### Figure 4A_NSP1.png

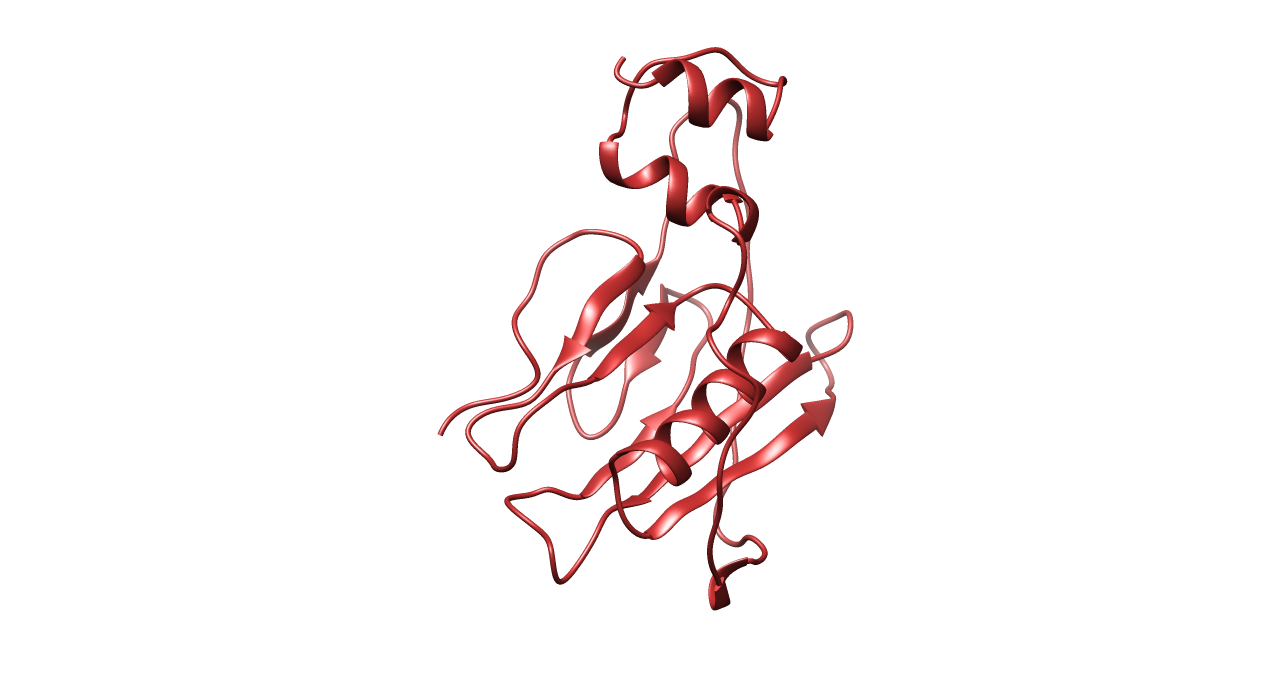

### Figure 4B_NSP2.png

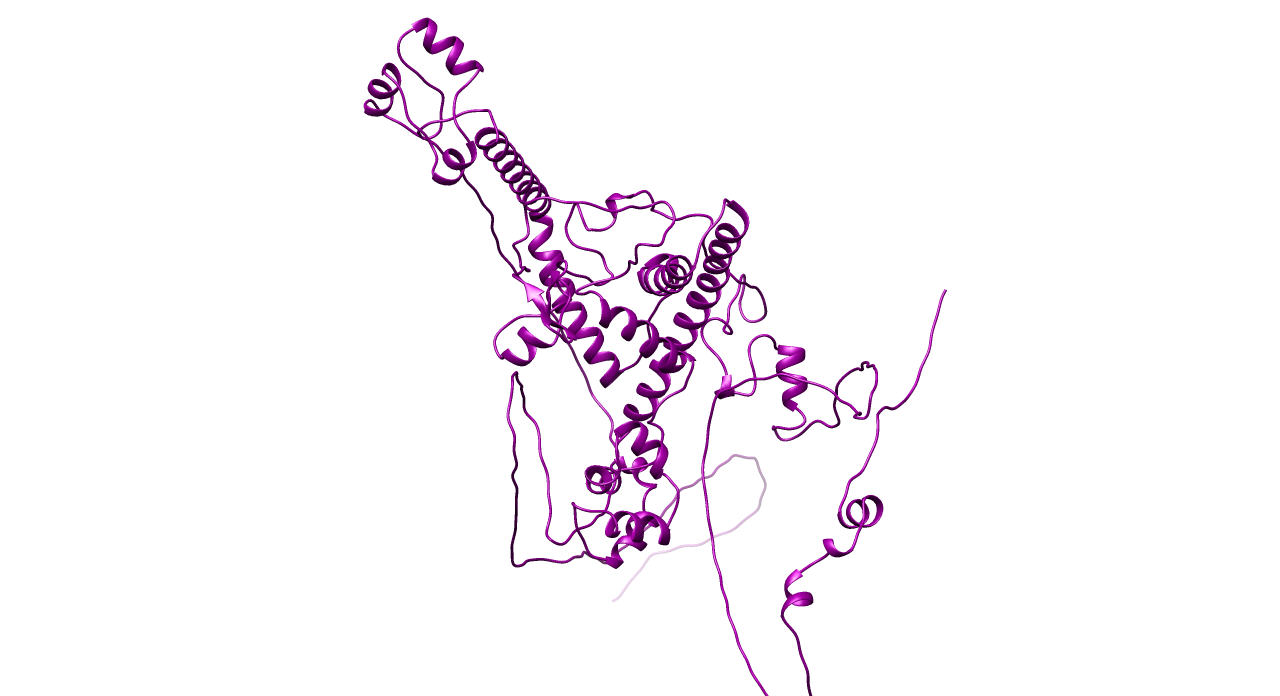

### Figure 4C_NSP3.png

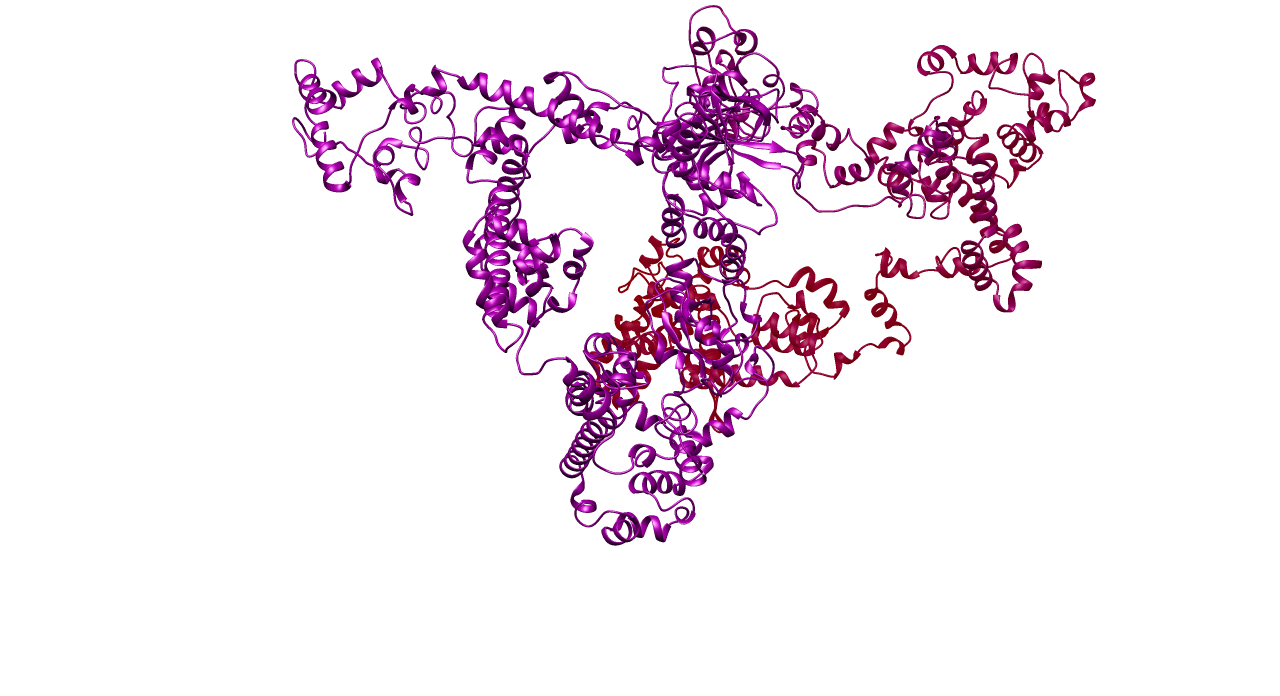

### Figure 4D_NSP4.png

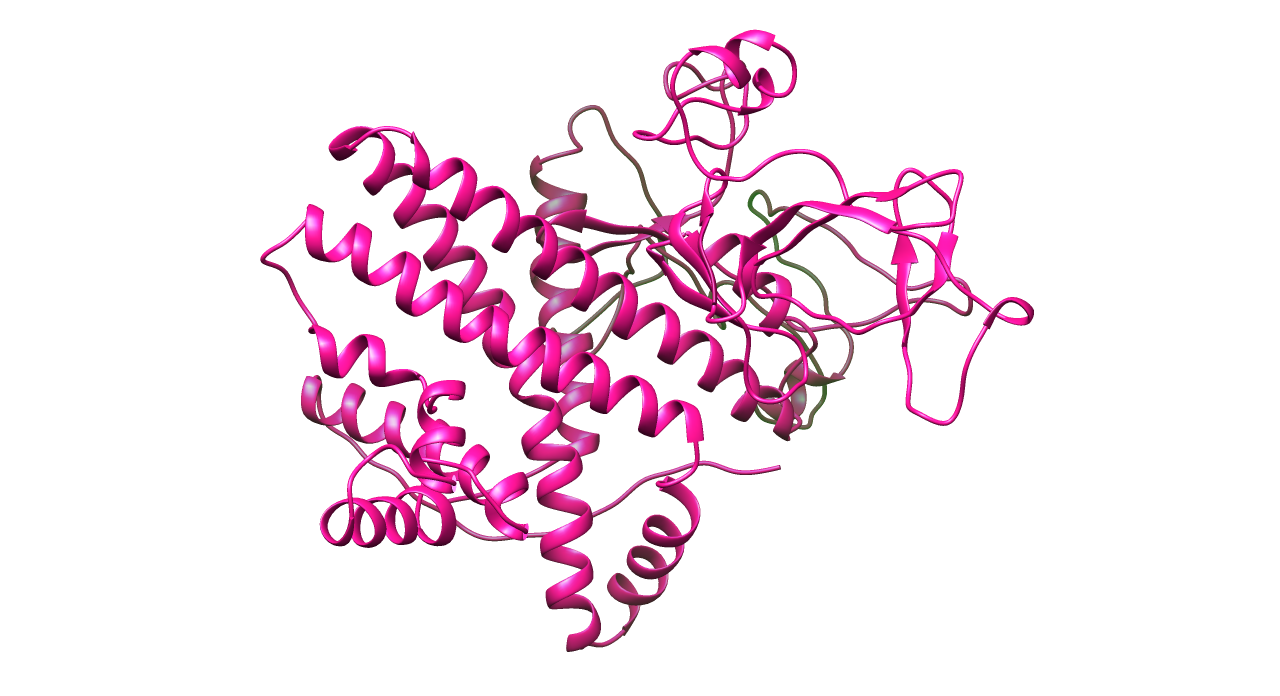

### Figure 4E_NSP6.png

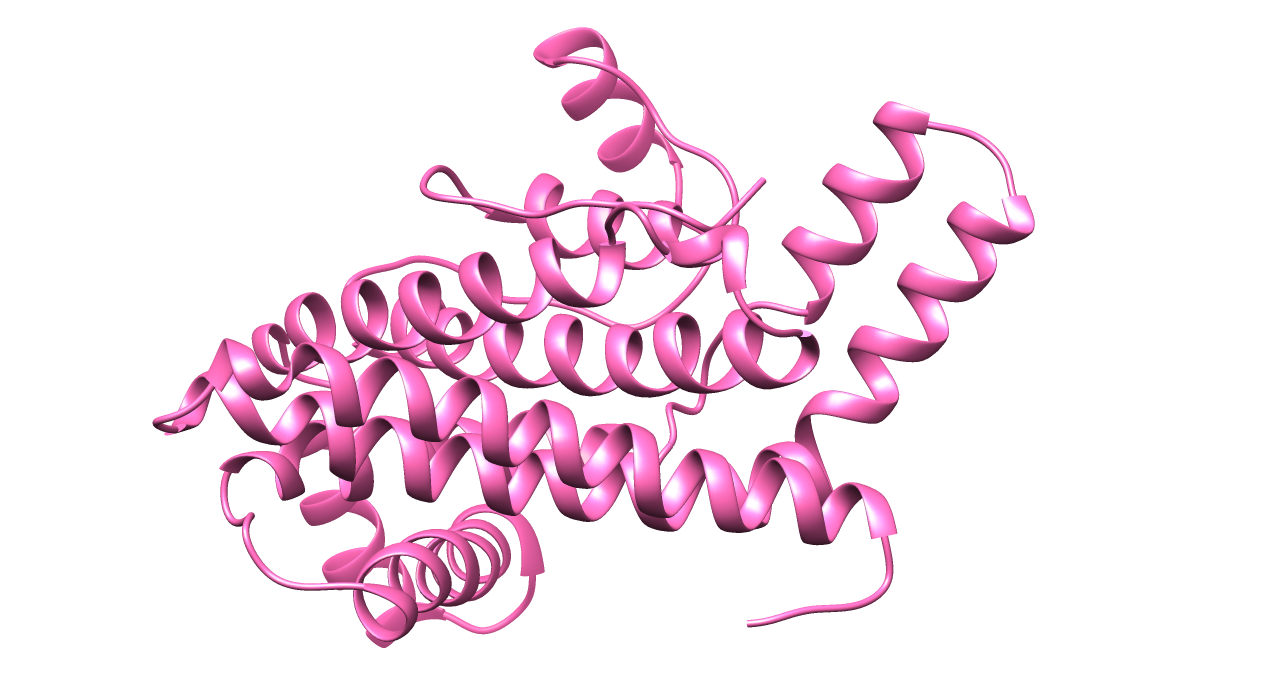

### Figure 4F_Surface glycoprotein.png

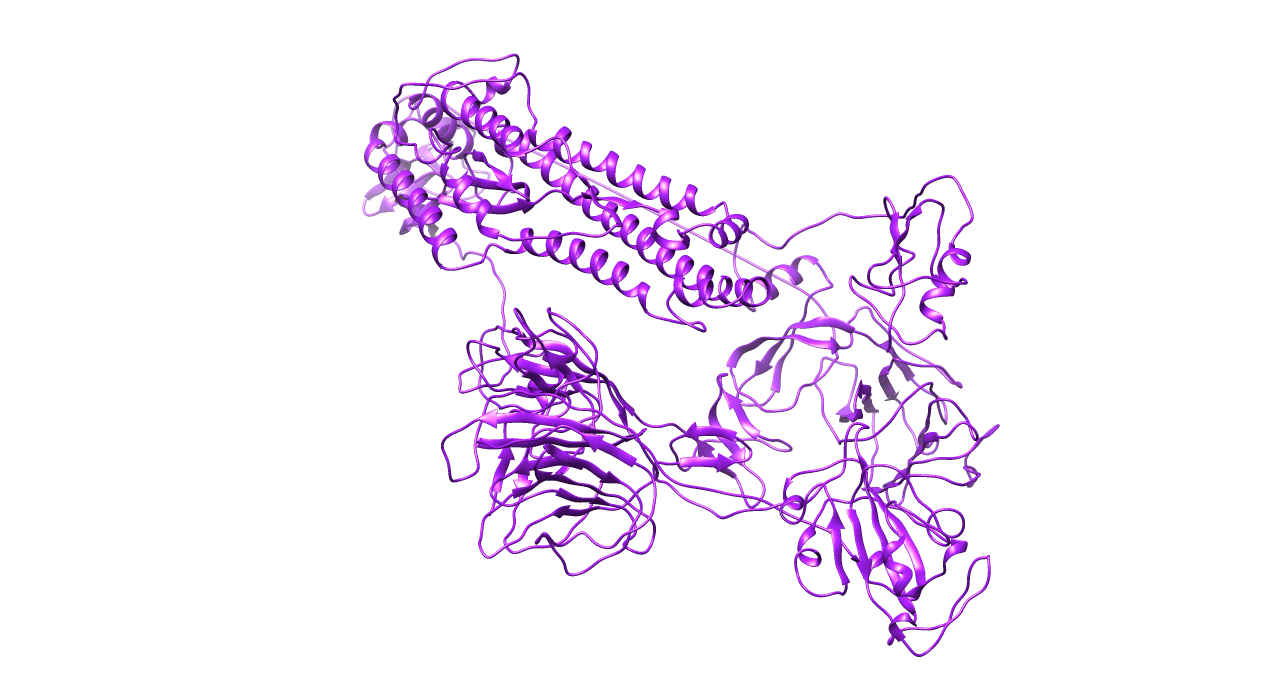

### Figure 4G_ORF3a protein.png

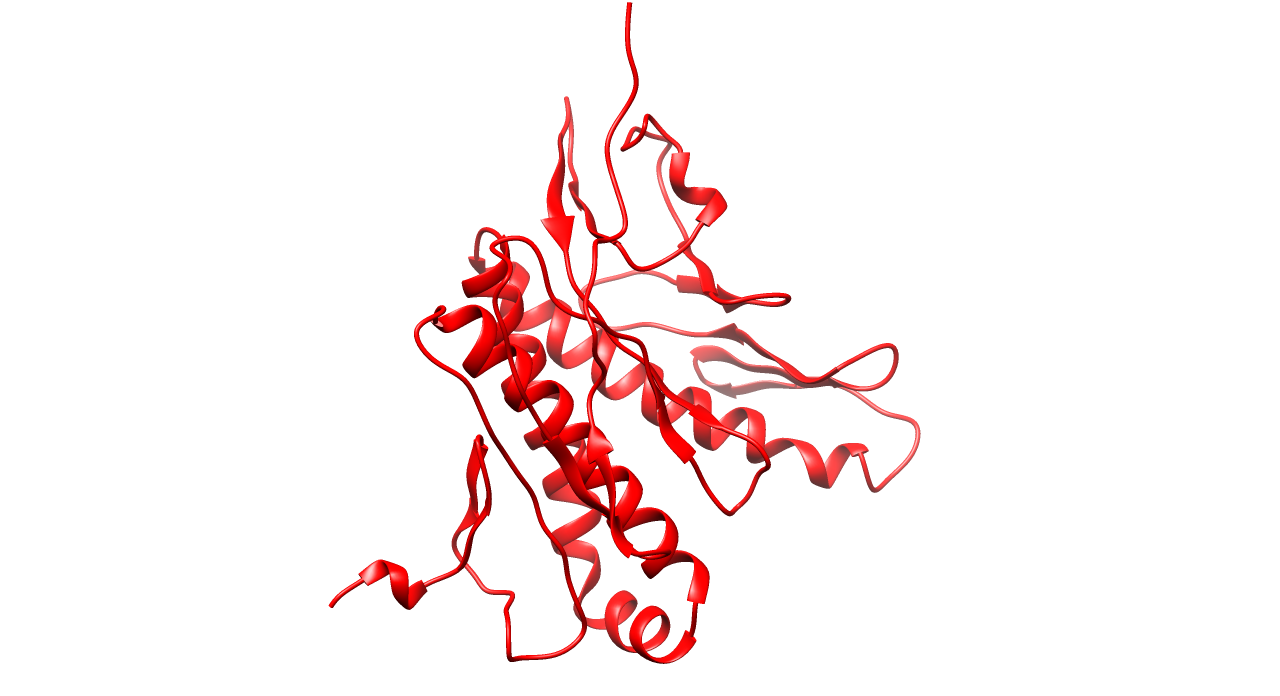

### Figure 4H_Envelope protein.png

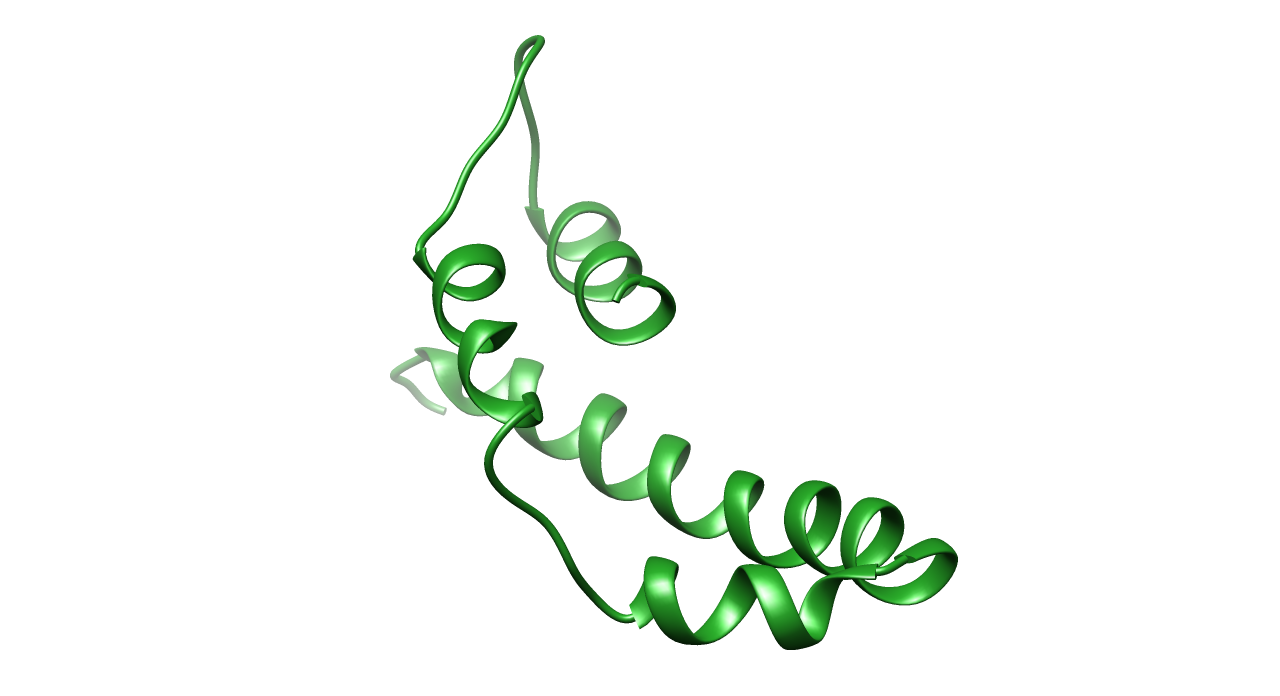

### Figure 4I_Membrane glycoprotein.png

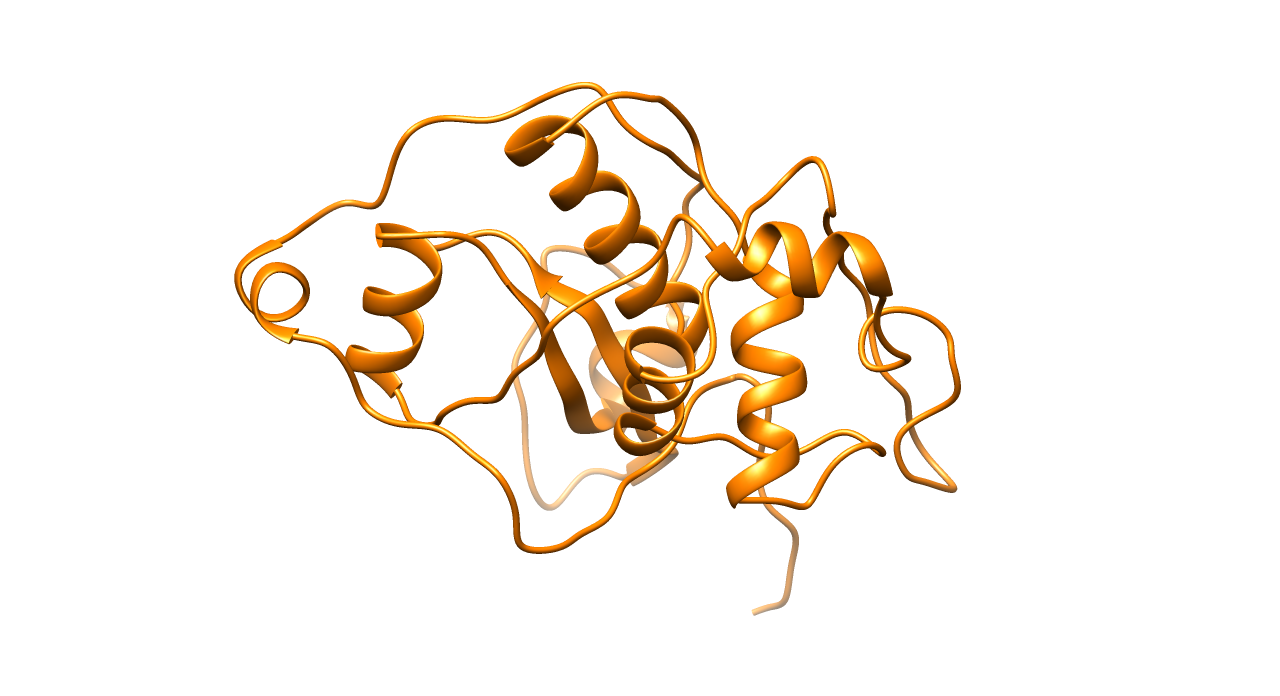

### Figure 4J_ORF6 protein.png

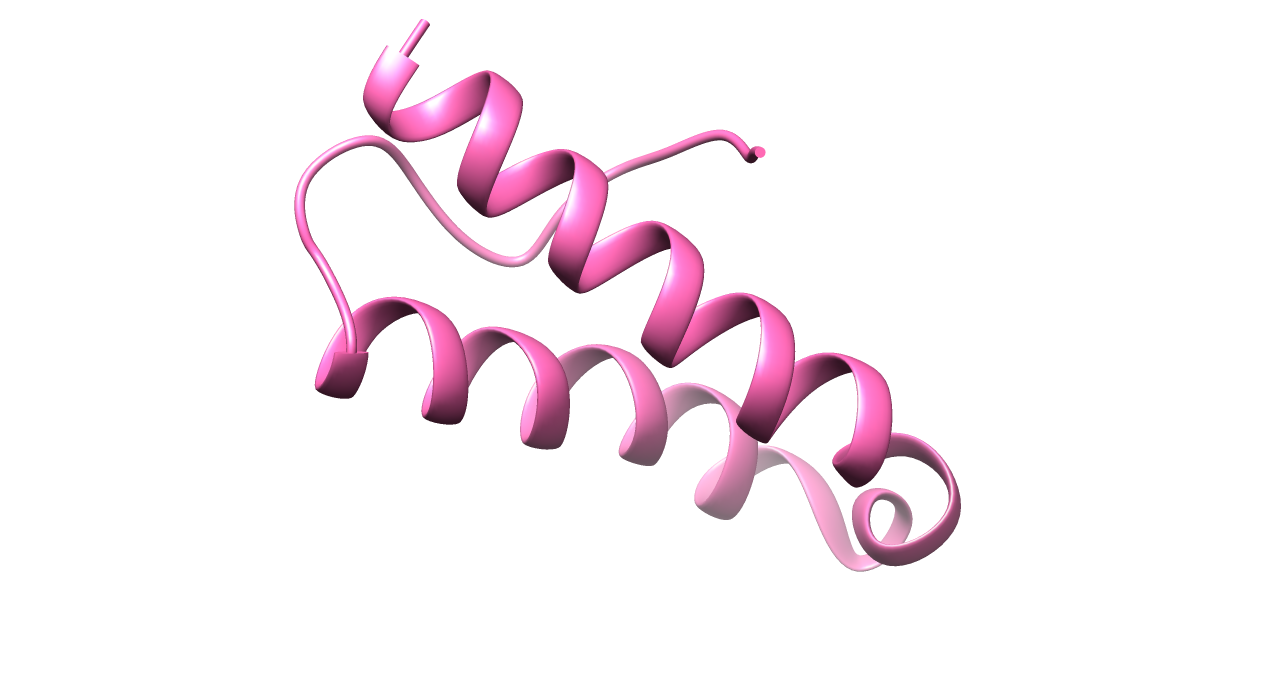

### Figure 4K_ORF7a protein.png

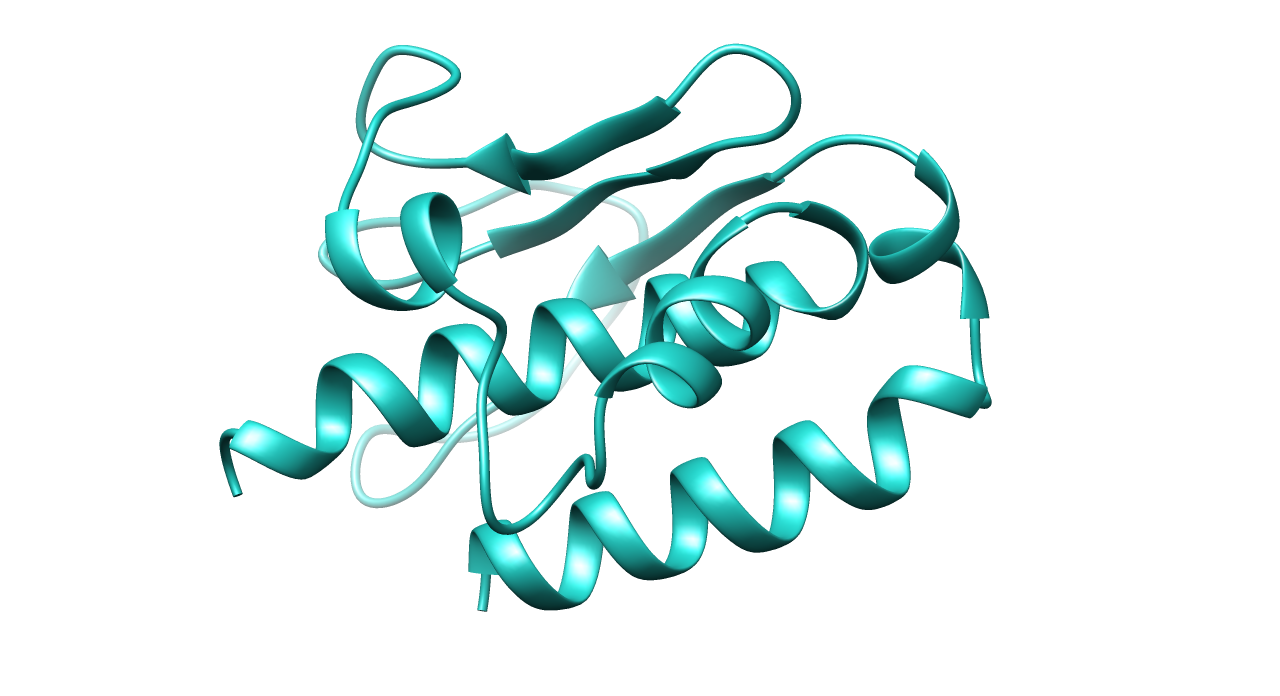

### Figure 4L_ORF7b protein.png

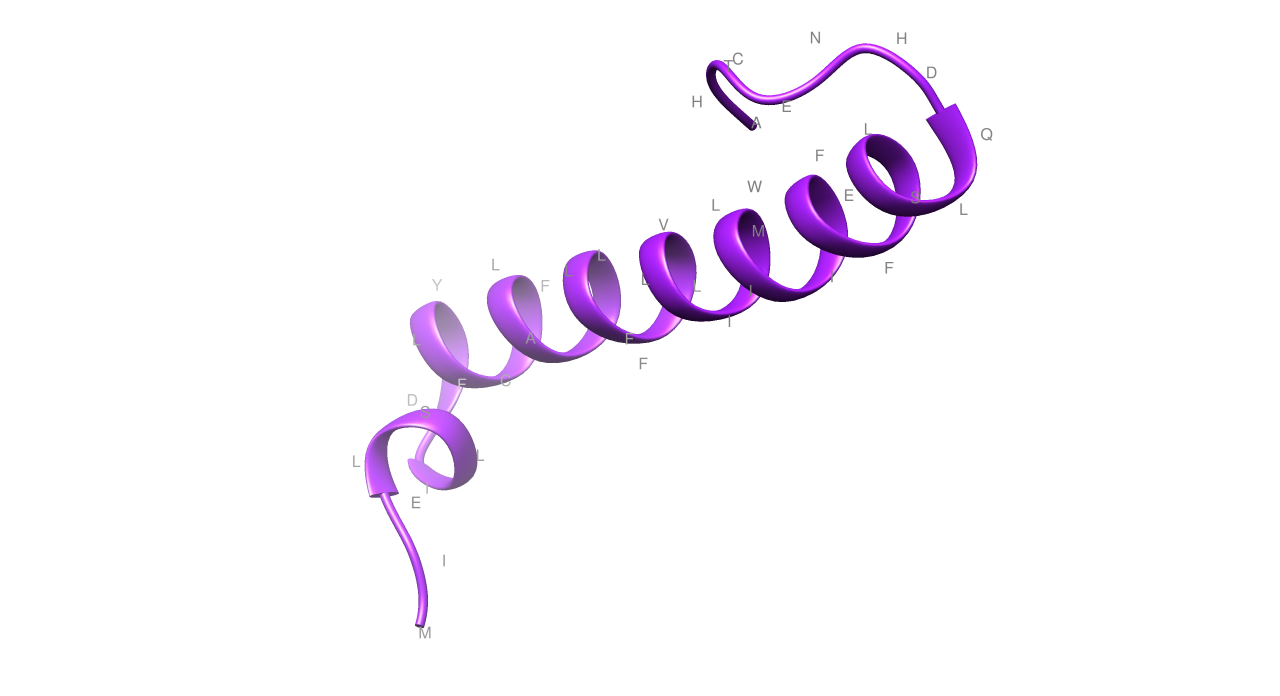

### Figure 4M_ORF8 protein.png

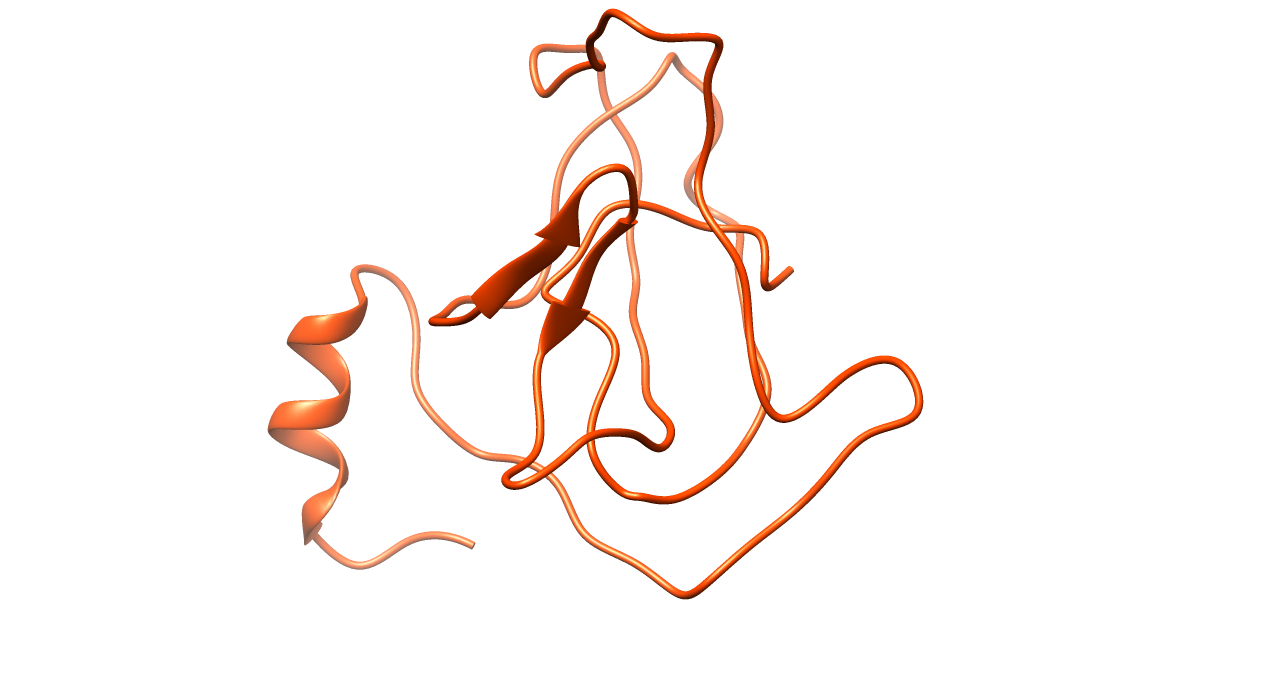

### Figure 4N_Nucleocapsid phosphoprotein.png

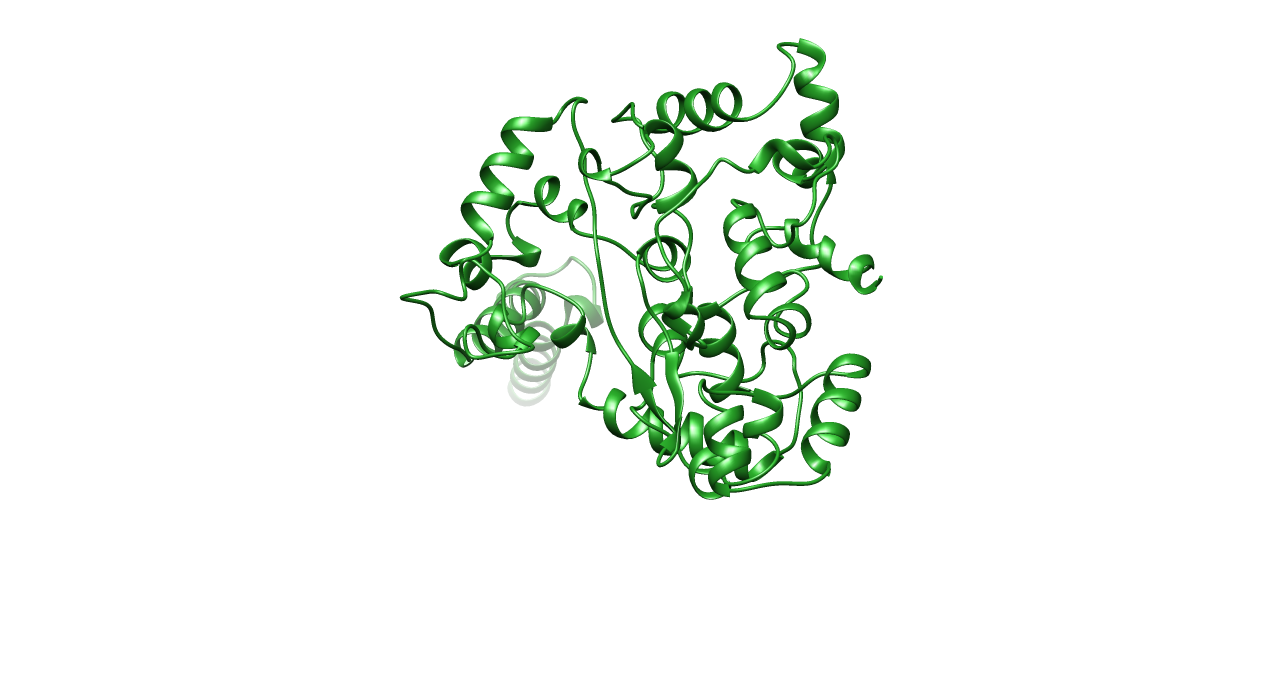

### Figure 4O_ORF10 protein.png

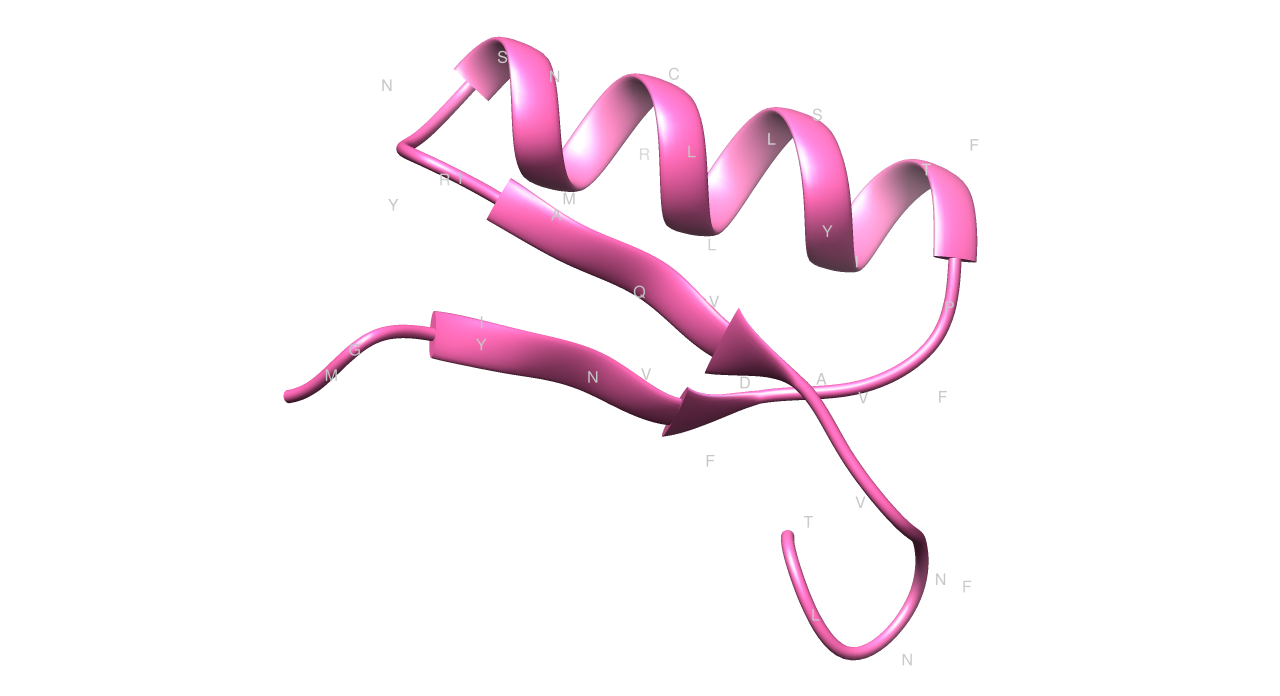
